# *Escherichia coli* SPFH membrane microdomain proteins HflKC contribute to aminoglycoside and oxidative stress tolerance

**DOI:** 10.1101/2022.07.25.501378

**Authors:** Aimee K. Wessel, Yutaka Yoshii, Alexander Reder, Rym Boudjemaa, Magdalena Szczesna, Jean-Michel Betton, Joaquin Bernal-Bayard, Christophe Beloin, Daniel Lopez, Uwe Völker, Jean-Marc Ghigo

**Affiliations:** Institut Pasteur, Université de Paris-Cité, CNRS UMR6047, Genetics of Biofilms Laboratory F-75015 Paris, France; Department of Functional Genomics, Interfaculty Institute for Genetics and Functional Genomics, University Medicine Greifswald, 17487 Greifswald, Germany; Abbelight, 191 avenue Aristide Briand, 94230 Cachan; Institut Pasteur, Université de Paris-Cité, UMR UMR6047, Stress adaptation and metabolism in enterobacteria 75015 Paris, France; Universidad Autonoma de Madrid, Centro Nacional de Biotecnologia, Campus de Cantoblanco, Calle Darwin 3, 28049 Madrid, España; Centre for Bacteriology Resistance Biology, Imperial College London, London, SW7 2AZ, UK; Departamento de Genética, Facultad de Biología, Universidad de Sevilla, Apartado 1095, 41080 Sevilla, Spain

**Keywords:** Membrane microdomains, SPFH proteins, Flotillin, Lipid raft, Stress tolerance, *Escherichia coli*

## Abstract

Many eukaryotic membrane-dependent functions are often spatially and temporally regulated by membrane microdomains (FMMs) also known as lipid rafts. These domains are enriched in polyisoprenoid lipids and scaffolding proteins belonging to the Stomatin, Prohibitin, Flotillin, and HflK/C (SPFH) protein superfamily that was also identified in Gram-positive bacteria. By contrast, little is still known about FMMs in Gram-negative bacteria. In *Escherichia coli* K12, 4 SPFH proteins, YqiK, QmcA, HflK, and HflC, were shown to localize in discrete polar or lateral inner-membrane locations, raising the possibility that *E. coli* SPFH proteins could contribute to the assembly of inner-membrane FMMs and the regulation of cellular processes.

Here we studied the determinant of the localization of QmcA and HflC and showed that FMM-associated cardiolipin lipid biosynthesis is required for their native localization pattern. Using Biolog phenotypic arrays, we showed that a mutant lacking all SPFH genes displayed increased sensitivity to aminoglycosides and oxidative stress that is due to the absence of HflKC. Our study therefore provides further insights into the contribution of SPFH proteins to stress tolerance in *E. coli*.

**IMPORTANCE:** Eukaryotic cells often segregate physiological processes in cholesterol-rich functional membrane micro-domains. These domains are also called lipid rafts and contain proteins of the Stomatin, Prohibitin, Flotillin, and HflK/C (SPFH) superfamily, which are also present in prokaryotes but were mostly studied in Gram-positive bacteria. Here, we showed that the cell localization of the SPFH proteins QmcA and HflKC in the Gram-negative bacteria *E. coli* is altered in absence of cardiolipin lipid synthesis. This suggests that cardiolipins contribute to *E. coli* membrane microdomain assembly. Using a broad phenotypic analysis, we also showed that HflKC contribute to *E. coli* tolerance to aminoglycosides and oxidative stress. Our study, therefore, provides new insights into the cellular processes associated with SPFH proteins in *E. coli*.

## INTRODUCTION

In addition to separating the intracellular content of cells from the environment, lipid bilayer membranes also contribute to specialized functions, including cross-membrane transport, enzymatic activity, signaling as well as anchoring of cytoskeletal and extracellular structures (1, 2). In eukaryotes, these membrane-dependent functions are often spatially and temporally regulated by functional membrane microdomains (FMMs) called lipid rafts (3–5). FMMs compartmentalize membrane cellular processes in cholesterol- and sphingolipid-enriched membrane regions formed upon lipid-lipid, lipid-protein and protein-protein interactions (5–7). A family of membrane proteins called SPFH proteins (for Stomatin/Prohibitin/Flotillin/HflK/C) has been shown to localize in eukaryotic lipid rafts and to recruit and provide a stabilizing scaffold to other raft-associated proteins (8–13).

Whereas most prokaryotes lack sphingolipids and cholesterol (14), the Gram-positive bacteria *Bacillus subtilis* and *Staphylococcus aureus* can also compartmentalize cellular processes in functional membrane microdomains (FMMs) (14–16). Although whether bacterial FMMs display a distinct lipidic composition needs to be more firmly established, they have been reported to be enriched in polyisoprenoid lipids as well as in cardiolipins at the cell poles (14–19)

FMMs also contain SPFH proteins, including flotillins, and a pool of proteins involved in diverse cellular processes (14, 16). In *B. subtilis*, flotillins FloT and FloA colocalize in membrane foci and contribute to the assembly of membrane protein complexes (20–23). Lack of flotillins impairs biofilm formation, sporulation, protease secretion, motility, and natural competence, indicating that the formation of FMMs also plays critical cellular roles in *B. subtilis* (15, 20, 24–27).

SPFH proteins are also present in Gram-negative bacteria, and *Escherichia coli* K12 even possesses four genes, *yqiK, qmcA, hflK*, and *hflC*, which encode proteins with an SPFH domain and a N-terminal transmembrane segment (28). QmcA and YqiK are predicted to face the cytoplasmic compartment, while HflK and HflC are predicted to be exposed in the periplasm, forming the HflKC complex negatively regulating the protease activity of FtsH against membrane proteins (29–32). HflC and QmcA are detected in *E. coli* membrane fractions resistant to solubilization by non-ionic detergents (detergent-resistant membranes or DRM) that are often used as - debatable - proxies for FMMs (33–36). Fluorescent microscopy also showed that *E. coli* SPFH proteins HflC and QmcA are localized in discrete polar or lateral membrane foci (37), raising the possibility that *E. coli* SPFH proteins could localize in inner-membrane FMMs and regulate specific cellular processes (38). However, apart from the functional and structural description of HflKC as a regulator of the FtsH membrane protease (29, 32, 39) and a recent study suggesting that YqiK is involved in cell motility and resistance to ampicillin (40), the functions of FMM in *E. coli* and other Gram-negative bacteria are still poorly understood.

In this study, we used fluorescent and super-resolution microscopy to perform a detailed analysis of QmcA and HflC membrane localization signals. We then showed that the integrity of QmcA and HflC protein domains is required for their inner membrane localization, and that the lack of cardiolipin and isoprenoid lipids known to associate with FMMs alters their localization. Moreover, using single and multiple SPFH gene mutants, we showed that HflKC SPFH proteins contribute to aminoglycosides and oxidative stress resistance. Our study therefore provides new insights into the determinants of cellular localization and the function associated with *E. coli* SPFH proteins.

## RESULTS

### Chromosomal *E. coli* SPFH fluorescent fusion proteins show distinct localization patterns

To investigate the determinant of cell localization of *E. coli* SPFH proteins, we first tagged YqiK and QmcA, which C-termini are predicted to be in the cytoplasm (31), with a C-terminal monomeric super folder green fluorescent protein (msfGFP). We then tagged HflC and HflK, which C-termini are predicted to be in the periplasm (31), with the C-terminal monomeric red fluorescent protein mCherry. All these fusions were expressed under their own promoter from their native chromosomal location (Sup. Fig. S1). Epifluorescence and super-resolution microscopy confirmed the previously reported polar localization of HflK and HflC (37) (Fig. 1 and Sup. Fig. S2), with 94% and 91% polar localization pattern for HflC-mCherry and HflK-mCherry, respectively (n=150). By contrast, C-terminally tagged QmcA-GFP showed punctate foci distributed throughout the cell body, with 96% of the cells harboring 5 foci or more (n=150) (Fig. 1CD). However, we could not detect YqiK-GFP, possibly due to its low native chromosomal expression level. We then used anti-GFP or mCherry antibodies to perform immunodetection on cytoplasmic as well as inner and outer membrane fractions of *E. coli* strains expressing either HflC-mCherry or QmcA-GFP. In agreement with previous results (31, 38), both fusion proteins were detected in the inner membrane fraction (Fig. 2).

**Figure 1.**
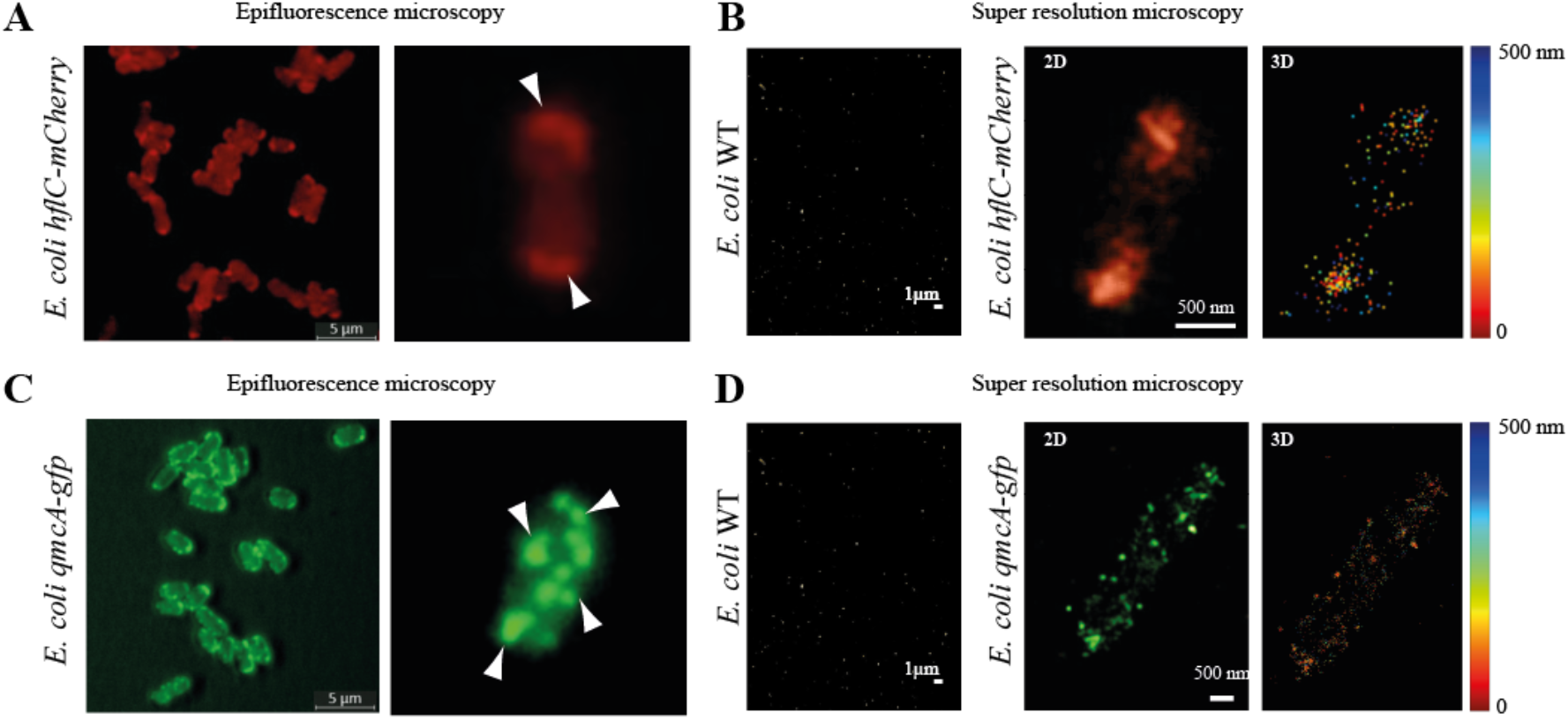
Cell localization patterns of HflC and QmcA. **A** and **C**: Epifluorescence microscopy of cells expressing HflC-mCherry (A) or QmcA-GFP (C). Arrowheads indicate polar or punctate localization foci. **B** and **D**: Super-resolution microscopy of cells expressing HflC-mCherry (B) or QmcA-GFP (D). *Left panels:* lack of subcellular localization in WT cells. *Center panels:* 3D images-artificial colors (red for HflC-mCherry, green for QmcA-gfp). *Right panels:* 3D localization of HflC-mCherry and QmcA-GFP, with colors corresponding to depth location along the Z axis, 0-500 nm, with 0 nm expressed in red, and 500 nm expressed in deep blue. Scales are indicated as white bars.

**Figure 2.**
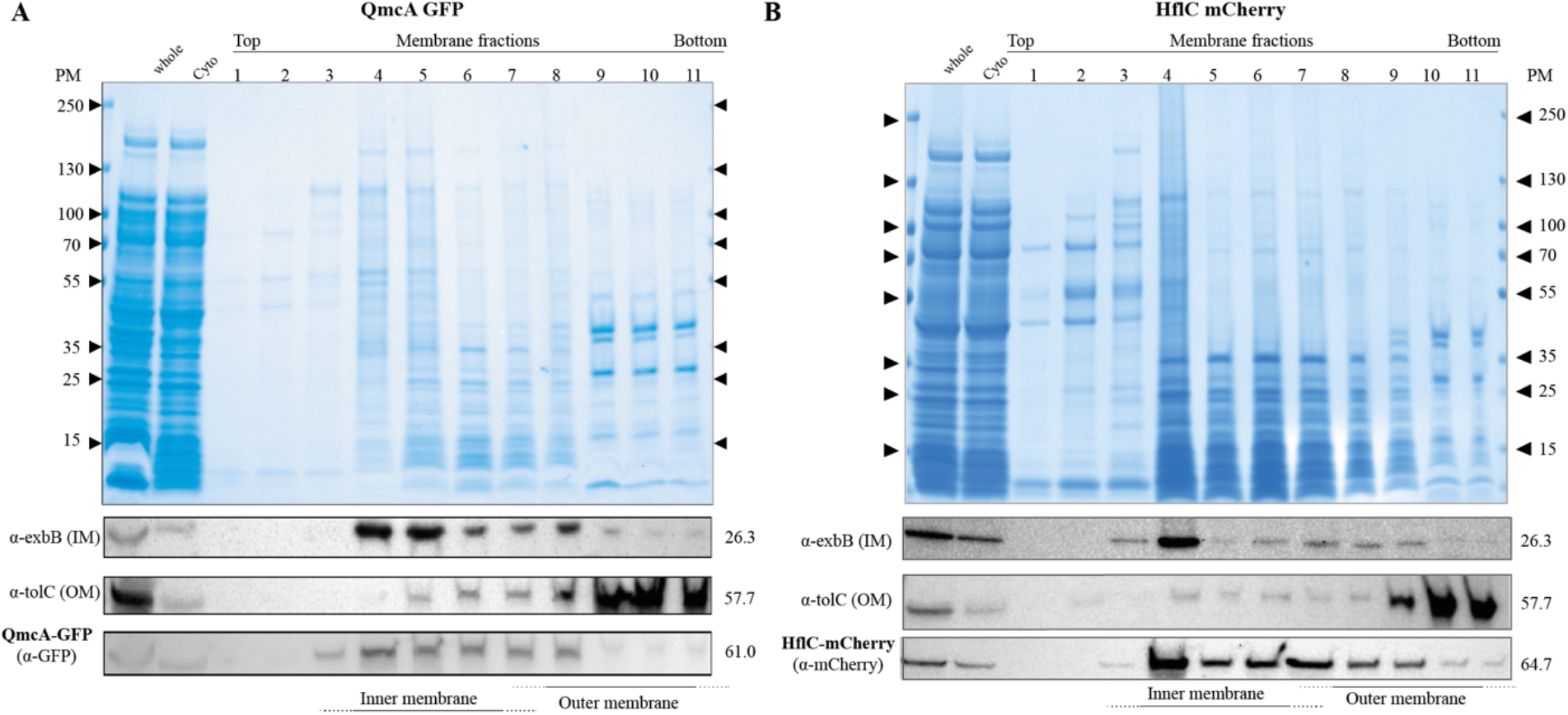
QmcA and HflC localize to the inner membrane. SDS-PAGE and immunodetection analyses of whole-cell extracts, cytosolic fractions, and inner (IM) or outer membrane (OM) fractions prepared from cells expressing QmcA-GFP (**A**) and HflC-mCherry (**B**). Anti-GFP and anti-mCherry antibodies were used to detect the presence of QmcA-GFP and HflC-mCherry, respectively. An anti-ExbB antibody were used to detect the inner membrane- (IM) marker ExbB and anti-TolC antibodies to detect the outer membrane- (OM) marker TolC.

### Domain swap analysis shows that protein integrity is essential for QmcA-GFP and HflC-mCherry localization

To identify HflC and QmcA membrane localization signals, we constructed multiple fluorescently tagged truncated versions of both proteins. We tagged with msfGFP a QmcA protein reduced to its transmembrane region and SPFH domain (TM^QmcA^-SPFH^QmcA^-GFP) and, separately, one reduced to the QmcA transmembrane region only (TM^QmcA^-GFP) (Fig. 3A). To test the role of the QmcA transmembrane region, we also swapped TM^QmcA^ in the three constructs by the single-spanning TM domain of the phage coat protein Pf3 (TM^Pf3^), which orients subsequent amino acids to the cytosol (38) (Fig. 3A). Similarly, in addition to the full length HflC-mCherry, we tagged with mCherry the HflC transmembrane region and SPFH domain (TM^HflC^-SPFH^HflC^-mCherry) and, separately, only its TM region (TM^HflC^-mCherry) (Fig. 3B). We also swapped the HflC TM region with the single-spanning TM region of colistin M immunity protein (TM^Cmi^), which orients subsequent amino acids to the periplasm (39) (Fig. 3B). Epifluorescence microscopy of HflC and QmcA derivative fusions showed that in addition to full-length constructs only full-length constructs with swapped TM (TM^Pf3^-QmcA-GFP and TM^Cmi^-HflC-mCherry) displayed significant punctate foci or polar localization, respectively (Fig. 3AB), although at reduced frequency compared to native QmcA-GFP and HflC-mCherry. Finally, we prepared inner and outer membrane fractions of *E. coli* strains expressing each QmcA and HflC derivative, and we observed that all these constructs were still mainly located in the inner membrane fraction. This indicates that, while we observed that QmcA-GFP and HflC-mChery derivatives exhibit altered cell localization, they do not exhibit significant mis-localization and remain located in the inner membrane (Sup. Fig. S3). These results therefore indicated that specific QmcA and HflC localization requires the combination of a transmembrane and full cytoplasmic (QmcA) or periplasmic (HflC) domain.

**Figure 3.**
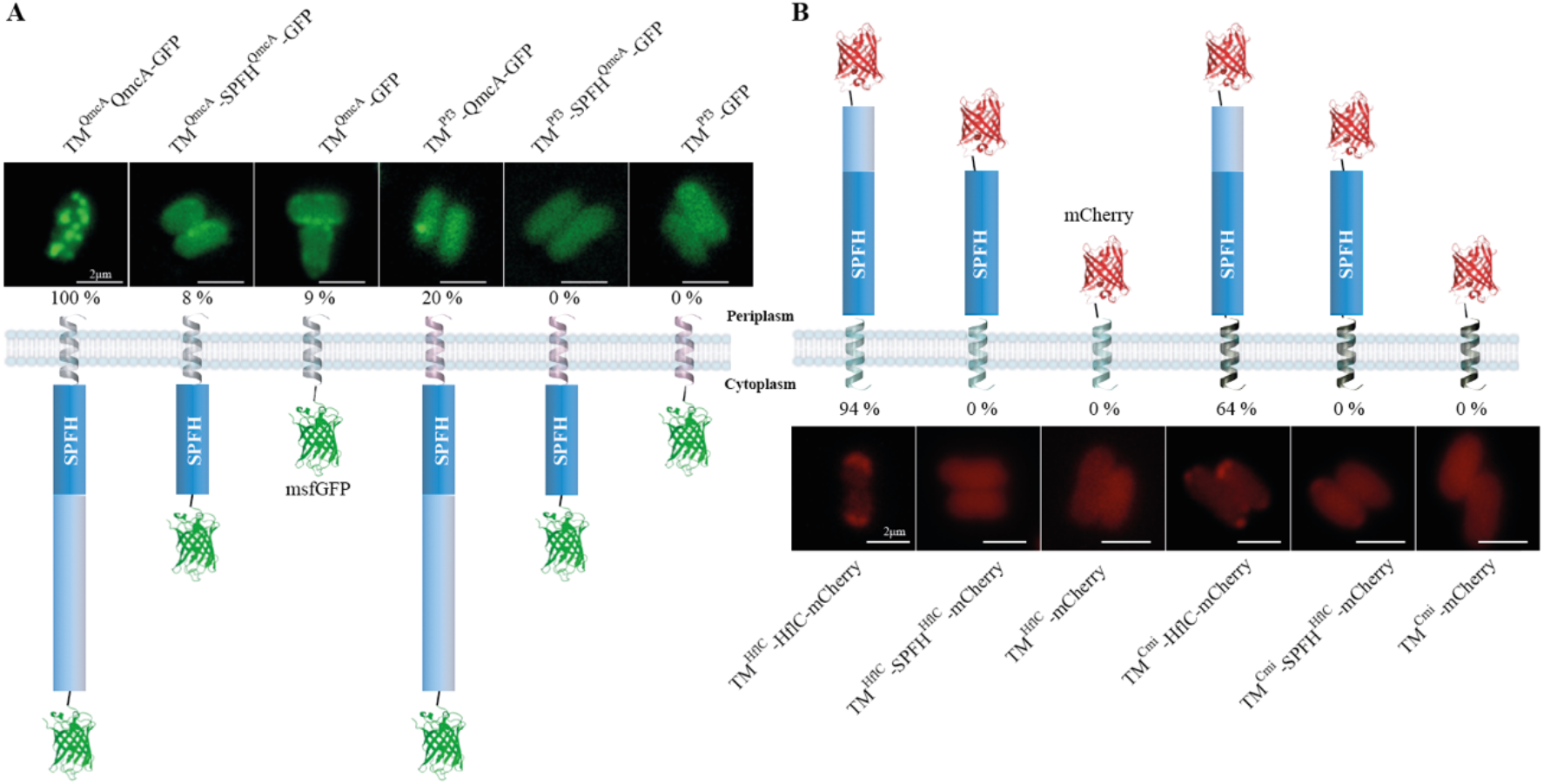
The localization pattern and membrane topology of full-length, domain swapped or truncated versions of QmcA and HflC. (**A**) GFP fusion derivatives of QmcA and (**B**) mCherry fusion derivatives of HflC. The representative images are shown in each strain with the frequencies of cells showing punctate (**A**) or polar localization (**B**). In membrane topology, helical structures represent transmembrane (TM) domains; silver, native TM domain of QmcA; pink, Pf3 domain; green, native TM domain of HflC; black, Cmi domain. Scale bars are 2 μm.

### Lack of cardiolipin and isoprenoid lipid synthesis alters the cell localization of QmcA and HflC

FMMs were proposed to be enriched with negatively charged cardiolipins and isoprenoids, which promote the localization of polar proteins and modulation of membrane lipid fluidity (15, 17, 18, 24, 41–44). We first tested whether alteration of cardiolipin synthesis could cause mis-localization of *E. coli* SPFH proteins QmcA or HflKC in a mutant lacking the major cardiolipin synthases *clsABC* (45). Whereas super-resolution microscopy analysis only showed an alteration of the number of QmcA-GFP punctates (1-5 cluster per bacterium) compared to WT (10-15 cluster per bacterium) (Fig. 4A and Sup. Fig. S4), the localization of HflC-mCherry showed a drastic loss of polar localization pattern (Fig. 4B and Supp. Fig. S4). We then used an *idi* mutant with reduced isoprenoid lipid synthesis due to the lack of isomerization of isopentenyl diphosphate (IPP) into dimethylallyl diphosphate (DMAPP) (46, 47). Whereas QmcA-GFP punctate localization was not affected, HflC-mCherry polar localization was not abolished in the Δ*idi* mutant (Fig. 4A and Sup. Fig. S4). These results demonstrated that the alteration of cardiolipin and, in a lesser extent, isoprenoid lipid synthesis pathway affects HflC fusion protein localization in *E. coli*.

**Figure 4.**
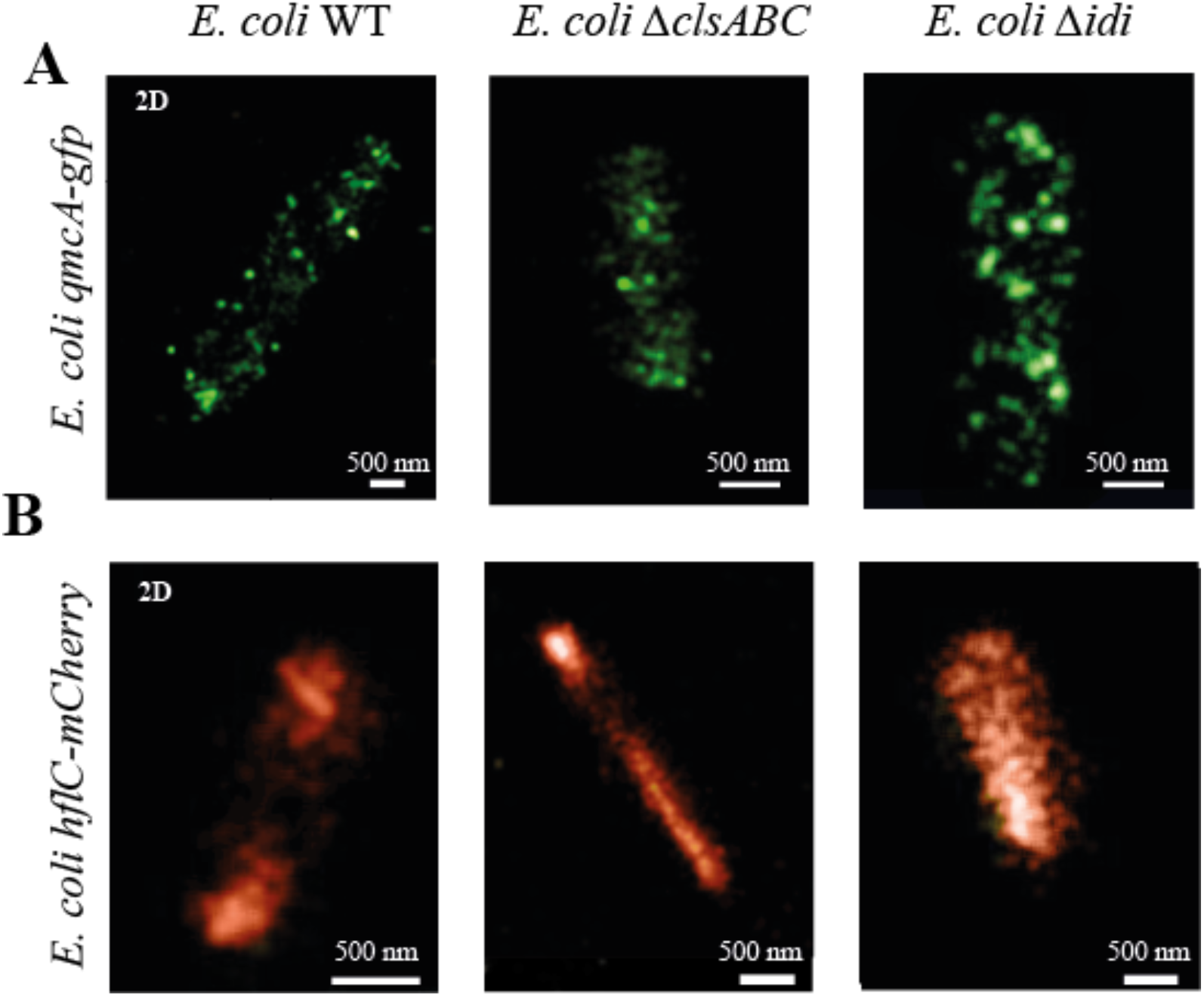
Alteration of QmcA and HflC cell localization in *E. coli* cardiolipin and isoprenoid pathway mutants. Two-dimensional super-resolution microscopy images of WT, Δ*clsABC*, and Δ*idi* strains expressing QmcA-GFP (**A**) and HflC-mCherry (**B**) in stationary phase.

### Phenotypic analysis of *E. coli* SPFH mutant shows that only the absence of HflKC increases *E. coli* sensitivity to aminoglycosides and oxidative stress

To identify potential phenotypes and functions associated with the *E. coli* SPFH proteins YqiK, QmcA, HflK and HflC, we introduced single and multiple deletions of the corresponding SPFH genes. We observed that neither single mutants nor the quadruple Δ*hflK, ΔhflC, ΔqmcA*, and Δ*yqiK* (hereafter referred to Δ*SPFH* mutant) displayed any significant growth defects in rich or minimal media (Fig. 5A and Sup. Fig. S5A). Considering the role of SPFH proteins in the activation of inner-membrane kinases involved in *B. subtilis* biofilm formation (15), we tested adhesion and biofilm capacity of WT and ΔSPFH strains but could not detect any significant differences between these two strains. We then used Biolog™ phenotypic microarrays to perform a large-scale phenotypic assay comparing an *E. coli* WT and Δ*SPFH* mutant (Sup. Table S1). This analysis revealed that the Δ*SPFH* mutant is metabolically less active when grown in the presence of various aminoglycosides (tobramycin, capreomycin, sisomicin and paromomycin) or when exposed to paraquat (Fig. 5B and Sup. Fig. S5BC). Consistently, the minimal inhibitory concentration (MIC) for tobramycin of the Δ*SPFH* mutant was 3-fold lower than that of the WT MIC (Fig.5C), and the sensitivity of the Δ*SPFH* mutant to paraquat was increased compared to the WT (Fig. 5D and Sup. Fig. S5D). Test of individual SPFH-gene mutants for their sensitivity to tobramycin and paraquat showed that the HflKC complex is the sole responsible for both phenotypes, as both single *hflK* and *hflC* or a double *hflKC* mutants displayed increased sensitivity to tobramycin and oxidative stress (Fig. 5CD and Sup. Fig. S5E). This phenotype could be complemented upon the introduction of a plasmid expressing *hflKC* genes in the double *hflKC* mutant (Sup. Fig. S6). Of note, C-terminally tagged HflC-mCherry and HflK-mCherry displayed wildtype MIC for tobramycin and paraquat, indicating that both fusions were functional and relevant proxies for the bacterial localization of the HflKC complex (Sup. Fig. S6).

**Figure 5.**
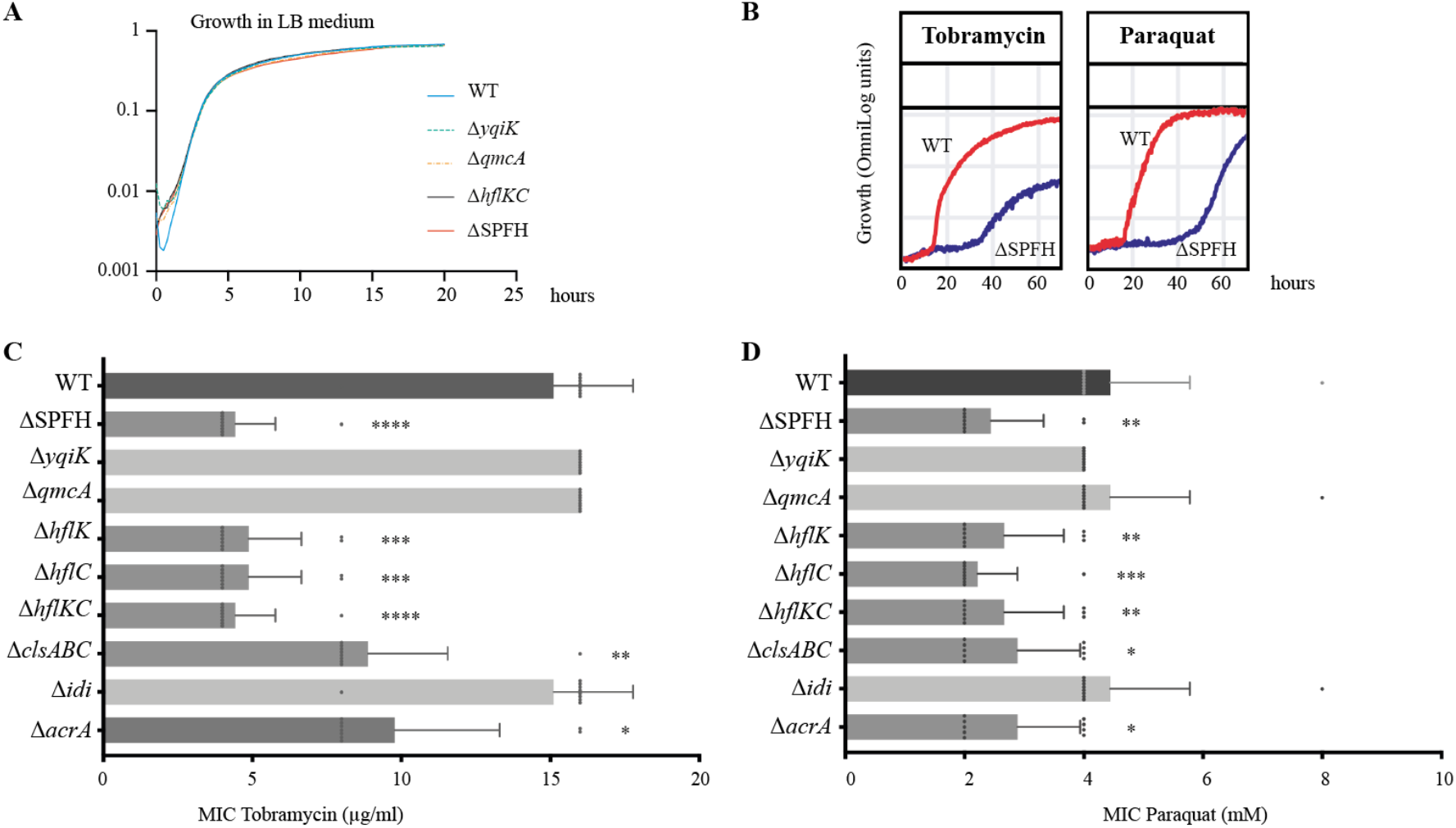
Phenotypic analysis of *E. coli* SPFH mutants. **A**: Bacterial growth curve of WT and SPFH gene deletion mutants in LB medium. **B**: Biolog bacterial growth curve of WT and ΔSPFH in the presence of tobramycin and paraquat. **C**: Minimum inhibitory concentration (MIC) for tobramycin of *E. coli* WT and indicated mutants. **D**: Minimum inhibitory concentration (MIC) for paraquat of *E. coli* WT and indicated mutants. \**p* < 0.05, \*\**p* < 0.01, \*\*\**p* < 0.001, \*\*\*\**p* < 0.0001compared with WT.

### Contribution of HflC localization to tobramycin and paraquat tolerance

Whereas an Δ*idi* mutant showed no significant difference compared to the WT, the MIC for tobramycin and paraquat of a Δ*clsABC* mutant was reduced by 2 to 3-fold, which is consistent with the respective impact of a *cls* and *idi* mutation on HflC localization (Fig. 4BD and Fig.5CD). Considering the scaffolding role of HflKC and the importance of cell localization, we tested the localization and contribution to aminoglycoside and stress tolerance of AcrA, a protein previously identified in *E. coli* inner-membrane DRM fractions often biochemically associated with FMM (37). AcrA is an efflux pump involved in the transport of a wide range of substrates including aminoglycosides (48). However, while an *acrA* deletion did not altered *E. coli* MIC profile to tobramycin and paraquat as much as a Δ*hflC* mutant (Fig.5CD), the expression of AcrA-GFP from their native chromosomal context did not lead to any distinct polar co-localization with HflC in exponential or stationary phase conditions (Sup. Fig. S7).

Taken together, these results indicate that the HflKC SPFH protein complex contributes to oxidative and antibiotic stress resistance.

## DISCUSSION

SPFH-domain proteins have been identified in most organisms (16, 49) and extensively studied in eukaryotes (3, 5, 50). By contrast, prokaryotic SPFH proteins and proteins associated with functional membrane microdomains (FMMs) are much less understood. This is particularly the case for Gram-negative bacteria, in which potential FMMs functions are mostly inferred from studies performed in *B. subtilis* and *S. aureus*. In this study, we investigated the functions and the localization determinants of *E. coli* SPFH proteins. Focusing on QmcA and HflKC SPFH proteins, we used a domain deletion and replacement approach and showed that most of the tested domain replacement variants correctly localized to the inner membrane but failed to display WT protein localization patterns. This indicates that inner-membrane localization alone is not sufficient for the correct subcellular distribution of HflC and QmcA, whose localization signals might rely on multiple addressing sequences spread throughout the entirety of each protein. Alternatively, it has been shown that oligomerization is critical for flotillin function and cluster formation (51), a process that is likely to be abolished in the domain deletion constructs, potentially explaining some of the observed localization defects.

QmcA and HflKC SPFH proteins display different localization patterns and could be part of different FMMs, potentially using different localization signals. The punctate localization pattern displayed by QmcA-GFP fusion was also observed in the case of *E. coli* YqiK expressed from plasmid and other Gram-positive bacteria flotillin homologues(15, 16, 18, 52, 53). Interestingly, *B. subtilis, B. anthracis* and *S. aureus* flotillin genes are physically associated with a gene encoding a NfeD protein, which could contribute to protein-protein interactions within flotillin assemblies (54, 55). Consistently, *E. coli qmcA* gene is located upstream of the NfeD-like *ybbJ* gene, like NfeD-like *yqiJ* gene is located upstream of *yqiK* (see Supplementary Figure S1). This further supports the notion that QmcA and YqiK could be considered as *E. coli bona fide* flotillins. By contrast, the *hflKC* transcription unit is not closely associated with a *nfeD-like* gene, suggesting that HflKC may not be a flotillin. However, while QmcA and YqiK have an opposite orientation to HflK and HflC, they are structurally similar proteins and the four *E. coli* SPFH proteins could therefore share some degrees of functionalities. The topological similarity between HflK and HflC might contribute to HflKC complex formation, and its interaction with FtsH protease, resulting in a large periplasmic FtsH-HflKC complex localized at the cell pole (29, 30, 56–58).

The strong negative impact of QmcA or HflC transmembrane (TM) domain replacement suggests that QmcA or HflC TM domains could recognize specific FMM membrane lipids facilitating the recruitment of SPFH proteins at their proper cellular membrane localization. Along with phosphatidylethanolamine and phosphatidylglycerol, cardiolipins are the primary constituent components of *E. coli* membranes that concentrate into cell poles and dividing septum (59–62). It was indeed observed that the composition of *E. coli* membrane lipids at cell poles is altered in a *clsABC* cardiolipin deficient mutant, compensated by an increased amount of phosphatidylglycerol (17, 63). Several studies also reported that cardiolipin-enriched composition in membranes at cell poles influences both the localization and activity of inner membrane proteins, such as respiratory chain protein complexes and the osmosensory transporter ProP (41, 42, 64–66). We showed here that, similarly to ProP, HflC and QmcA localization patterns are affected in a Δ*clsABC* mutant, suggesting that HflKC and QmcA complexes could act as scaffolds for cardiolipin-enriched FMM cargo proteins. Isoprenoid lipids such as farnesol, carotenoids, and hopanoids have been proposed to be constituents of bacterial FMMs or to interact with SPFH proteins and FMM-associated proteins (14). Consistently, blocking the *S. aureus* carotenoid synthetic pathway by zaragozic acid leads to flotillin mis-localization (15), and the inactivation of farnesol synthesis in a *B. subtilis yisP* mutant impairs focal localization of the FMM-associated sensor kinase KinC (15). We showed that interfering with the *E. coli idi* isoprenoid biosynthesis pathway also strongly alters the localization of HflC. This further indicates that isoprenoid lipids contribute to the formation or integrity of FMMs, possibly by altering isoprenoid-dependent membrane fluidity, as shown in *S. aureus* and *B. subtilis* FMMs (14, 67).

Our investigation of the phenotypes displayed by an *E. coli* mutant lacking all SPFH protein genes showed that the absence of HflKC leads to an increased susceptibility to oxidative stress and aminoglycosides. The HflKC complex was previously shown to modulate the quality control proteolytic activity of FtsH by regulating the access of misfolded membrane protein products to FtsH (29, 30, 68). *E. coli ΔhflK* and Δ*hflC* mutant strains were also shown to accumulate increased amounts of hydroxyl radical, suggesting that HflK and HflC could influence tolerance to aminoglycosides and oxidative stress by suppressing excessive hydroxyl radical production. Alternatively, HflK and HflC could contribute to tobramycin resistance via FtsH-dependent proteolytic activity (69) or favoring FMM formation and the assembly of membrane proteins and lipids, such as cardiolipin, involved in the transport and movement of aminoglycosides within cells and cell membranes. Consistently, several proteins associated with aminoglycosides transport were actually detected in *E. coli* DRM fractions, including transporters and several components of the AcrAB-TolC efflux pump (37), suggesting that deletion of *hflK* or *hflC* could reduce the activity of these proteins in FMMs and enhance entry of aminoglycosides. Whereas the susceptibility to aminoglycosides indeed partly relies on the AcrAB-TolC efflux pump (48, 70–72), we found that lack of the AcrA only moderately decreases the MIC to tobramycin, compared to a *hflKC* mutation. We also showed that the alteration of the cardiolipin pathway in a Δ*clsABC* mutant altered both the localization and sensitivity to tobramycin and paraquat of a HflC-GFP fusion. This suggests that cardiolipin could be required for the correct localization of HflKC to FMMs at cell poles. By contrast, although an *idi* isoprenoid pathway mutant mildly affects HflC localization, it does not show altered sensitivity. This may be due to an uncomplete inactivation of the pathway since lycopene production in a Δ*idi* mutant is reduced by 1/3 but not totally abolished, due to the fact that the idi protein is a reversible isomerase. (46).

In conclusion, the present study provides new insights into the functions of *E. coli* SPFH proteins and some of their interacting partners and further experiments will be needed to fully uncover the roles played by this intriguing family of membrane proteins in Gram-negative bacteria.

## MATERIALS & METHODS

### Bacterial strains and growth conditions

Bacterial strains and plasmids used in this study are described in Sup. Table S2, and further explained in Sup. Fig. S1 and Figure 3. Unless stated otherwise, all experiments were performed in lysogeny broth (LB) or M63B1 minimal medium supplemented with 0.4% glucose (M63B1.G) at 37 °C. Antibiotics were used as follows: kanamycin (50 μg/mL); chloramphenicol (25 μg/mL); ampicillin (100 μg/mL); and zeocin (50 μg/mL). All compounds were purchased from Sigma-Aldrich (St Louis, MO, USA) except for Zeocin (InvivoGen, Santa Cruz, CA, USA).

### Mutant construction

#### Generation of mutants in E. coli

Briefly, *E. coli* deletion or insertion mutants used in this study originated either from the *E. coli* Keio collection of mutants (73) or were generated by λ-red linear recombination using pKOBEG (Cm^R^) or pKOBEGA (Amp^R^) plasmids (74) using primers listed in Sup. Table S3. P1vir transduction was used to transfer mutations between different strains. When required, antibiotic resistance markers flanked by two FRT sites were removed using the Flp recombinase (75). Plasmids used in this study were constructed using an isothermal assembly method, Gibson assembly (New England Biolabs, Ipswich, MA, USA) using primers listed in Sup. Table S3. The integrity of all cloned fragments, mutations, and plasmids was verified by PCR with specific primers and DNA sequencing

#### Construction of deletion mutants

Δ*yqiK*, Δ*qmcA*, Δ*hflK*, Δ*hflC*, Δ*clsA*, Δ*clsB*, Δ*clsC*, Δ*idi*, Δ*acrA*, deletions were transferred into *E. coli MG1655strep* by P1vir phage transduction from the corresponding mutants in the *E. coli* BW25113 background of the Keio collection (73). The associated kanamycin marker was then removed using the Flp recombinase expressed from the plasmid pCP20 (75). (Details regarding the construction of all other strains used in this study are presented in Sup.Table S2).

#### Construction of GFP and mCherry fusions

See Supplementary Methods section in Supplementary Materials

#### Construction of complemented strains

The *hflKC* genes were amplified from MG1655*strep* using primers listed in Supplementary Table S3 and cloned at the downstream of the IPTG-inducible promoter of a pZS*12 vector using the Gibson assembly to generate plasmids pZS*12-HflKC Then, these plasmids were introduced into Δ*hflKC* mutants, respectively, to construct complemented mutants (Sup. Table S2). A pZS*12 empty vector was also introduced into wildtype and Δ*hflKC* mutants. Mutants harbouring these pZS*12 plasmids were incubated and used for the below experiments in the presence of IPTG (1 mM) and ampicillin.

### Epifluorescence microscopy

Bacteria were incubated into 5 mL of fresh LB medium and harvested at OD_600_ 0.4 for samples in exponential phase or OD_600_ 2.0 for stationary phase. After washing twice with M63B1 medium, cells corresponding to 1 mL of the bacterial culture were pelleted by centrifugation and resuspended into 0.1 mL of M63B1 medium for exponential samples, or 1 mL of the medium for stationary samples. Ten μL aliquots of the cell suspension were immobilized on glass slides previously covered with freshly made M63B1 medium 0.8% agarose pads. Cells were observed using a ZEISS Definite focus fluorescent microscope (Carl Zeiss, Oberkochen, Germany), equipped with an oil-immersion objective lens microscope (Pln-Apo 63X/1.4 oil Ph3). GFP or mCherry fluorescence was exited with a ZEISS Colibri LED illumination system and the fluorescence signal was detected with Zeiss FS38 HE (Carl Zeiss) or Semrock HcRed filters (Semrock, Rochester, NY, USA). GFP, and mCherry fluorescence images were taken at 1000, and 2000 msec. exposure, respectively. Image processing was performed using ImageJ and Adobe Photoshop. For each tested mutant, the subcellular localization patterns of 150 randomly selected bacteria were evaluated and the frequencies were expressed as percentiles.

### Super-resolution microscopy

Bacteria were imaged using single-molecule localization microscopy and stochastic optical reconstruction microscopy (SMLM-STORM), using a previously described method (76). Overnight cultures were fixed with PFA 4%, permeabilized with Triton 0,05%, and labeled with either GFP monoclonal FluoTag®-Q — Sulfo-Cyanine 5 (Cy5), or RFP monoclonal FluoTag®-Q – Cy5, which are single-domain antibodies (sdAb) conjugated to Cy5. Labeling was performed at 1:250 (concentration), and washing steps were carried out three times using Abbelight’s SMART kit buffer. For imaging, Abbelight’s imaging system was used with NEO software. Abbelight’s module was added to an Olympus IX83 with 100x TIRF objective, N.A. 1.49. We used Hamamatsu’s sCMOS Flash 4 camera and a 647nm 500mW Oxxius laser, with an astigmatic lens, to allow for 3D imaging of the sample (77).

### Inner membrane separation

*E. coli* overnight cultures were diluted into 1 L of fresh LB medium to OD_600_ of 0.02 and incubated at 37°C and 180 rpm until reaching OD_600_ 0.4. Cells were harvested and washed once with 10 mM HEPES (pH 7.4) and stored at −20°C for at least 1 h. Bacteria were then resuspended in 10 mL of 10 mM HEPES (pH 7.4) containing 100 μL of Benzonase (3.10^4^ U/mL) and were passed through a FRENCH press (Thermo) at 20,000 psi. The lysate was centrifugated at 15,000 *g* for 15 min at 4 °C to remove cell debris, and aliquots of the suspension were stored at 4 °C as the whole extract. Then, the suspension was centrifuged at 100,000 *g* for 45 min at 4 °C to separate supernatant and pellets, and aliquots of the supernatant were stored at 4 °C as the cytosolic and periplasmic fractions. The pellets were suspended into 600 μL of cold 10 mM HEPES (pH 7.4) and homogenized by using 2 mL tissue grinder (Kontes Glass, Vineland, NJ, USA). Discontinuous sucrose gradients with the following composition were placed into ultracentrifugation tube: bottom to top 0.5 mL of 2 M sucrose, 2.0 mL of 1.5 M sucrose, and 1.0 mL of 0.8 M sucrose, and 500 μL of the homogenized samples were placed on the top of sucrose gradients. The gradients were centrifuged at 100,000 *g* for 17.5 h at 4 °C. Subsequently, 400 μL of aliquots were collected into 11 microtubes from top to bottom, and the samples were proceeded to the immunodetection method, as described below.

### Immunodetection of inner membrane proteins

Aliquots of samples were suspended in 4× Laemmli buffer (BioRad) with 2-Mercaptoethanol (Sigma) and incubated for 5 min at 98 °C. The protein samples (10 μL each) were run on 4-20 % Mini-PROTEAN TGX Stain-FreeTM precast Gels (BioRad) in 1× TGX buffer and then transferred to nitrocellulose membrane using a Trans-Blot® Turbo™ Transfer System (BioRad). Subsequently, the membranes were blocked using blocking buffer consisting of 5% skim milk in PBS with 0.05% Tween 20 (PBST) for 2 h at 4 °C with agitation. The membranes were then incubated in PBST containing 1% skim milk with first antibodies, polyclonal rabbit antiserum raised against ExbB and TolC (kindly given by Dr. Philippe Delepelaire), GFP (Invitrogen, A6455, Thermo Fisher Scientific, Indianapolis, IN, USA) and mCherry (Invitrogen, PA5-34974) at 1:20,000 overnight at 4 °C with agitation. The membranes were washed in PBST and incubated in PBST containing 1% skim milk with a secondary antibody, anti-rabbit IgG conjugated with horseradish peroxidase (Cell signaling, 7074S), at 1:10,000 for 2 h at 25 °C with agitation. After washing the excess secondary antibody, specific bands were visualized using the ECL prime detection method (GE Healthcare) and imaged with an imaging system, iBright™ CL1500 (Invitrogen).

### Microbial growth phenotypic analysis

A high-throughput analysis for microbial growth phenotypes using a colorimetric reaction, Phenotype MicroArrays (Biolog Inc., Hayward, CA, USA), was performed in accordance with the manufacturer’s protocol. Briefly, several colonies of *E. coli* grown on LB agar were transferred in 10 mL of a mixture of Biolog IF-0a media (BioLog) and sterilized water into a sterile capped test tube. The suspension was mixed gently, and the turbidity was adjusted to achieve the appropriate transmittance using the Biolog turbidimeter (BioLog). The cell suspension was diluted with the IF-0a plus dye mix, as mentioned in the manufacturer’s protocol. 100 μL of the mixture suspension was inoculated into PM plates 1-3 and 9-20 and incubated for 72 h at 37 °C. The absorbance of each well was taken every 15 min. The OmniLog software (BioLog) was used to view and edit data, to compare data lists, and to generate reports.

### Monitoring of bacterial growth

An overnight culture of *E. coli* was diluted into fresh LB and M63B1 supplemented with 0.4% glucose medium to OD_600_ of 0.05, and 200 μL aliquots were cultured in the presence or absence of paraquat (Methyl viologen dichloride hydrate, Sigma-Aldrich) in 96-well microplates at 37 °C for 24 h with shaking. The absorbance of each culture at 600 nm was measured every 15 min for 24 h using a microplate reader (Tecan Infinite, Mannedorf, Switzerland).

### Susceptibility of *E. coli* against Tobramycin and Paraquat

The broth microdilution method was used to determine the MIC (minimum inhibitory concentration) values of Tobramycin (Sigma-Aldrich) and Paraquat (Sigma-Aldrich) in 96-well microtiter plates. Briefly, 100 μL of LB medium was distributed into each well of the microtiter plates. Tobramycin was 2-fold serially diluted in each well. Five μL of approximately 1 × 10^7^ CFU/mL of *E. coli* was inoculated into each well, and the plates were incubated at 37°C for 24 h. The lowest concentration that visibly inhibited bacterial growth was defined as the MIC. All strains were evaluated in biological and technical triplicates.

The spot assay was performed to evaluate the susceptibility of *E. coli* against paraquat. Briefly, an overnight culture of *E. coli* was diluted into fresh LB medium to OD_600_ of 0.05. Ten μL of the diluted culture was spotted on LB plates containing either no or 100 μM paraquat. The plates were incubated at 37°C for 24 h, and the photographs were taken. All strains were evaluated in triplicate.

## Supporting information

This pdf file includes supplementary methods, supplementary figures S1 to S7 and supplementaryTables S2 and S3.

## Statistical analysis

Data analysis was performed using GraphPad Prism 9.5 software (GraphPad, La Jolla, CA, USA). All data are expressed as mean (± standard deviation, SD) in figures. Statistical analysis was performed using unpaired non-parametric Mann-Whitney test. Differences were considered statistically significant for *P* values of <0.05.

## ACKNOWLEDGMENTS

We thank Philippe Delepelaire for insightful comments and material support. We thank Uwe Sauer and Philip Warmer for initial assessment of lipid composition of some of the strains used in this study. We are grateful to Eva Wolrab and Sven van Teeffelen for their initial interest in the project and for providing the strains for msfGFP and mCherry constructions. This work was supported by EU Horizon 2020 Rafts4Biotech grant 720776 (to JMG, DL, AKW, YY, AR, UV and MS), the French government’s Investissement d’Avenir Program, Laboratoire d’Excellence “Integrative Biology of Emerging Infectious Diseases” (grant n°ANR-10-LABX-62-IBEID) and the *Fondation pour la Recherche Médicale* (grant no. DEQ20180339185). This work benefited from the facilities and expertise of Add Photonic BioImaging platform (UTechS PBI, Institut Pasteur). A.K.W. was supported by a Pasteur-Roux-Cantarini postdoctoral and a grant from the Philippe Foundation.

## AUTHOR CONTRIBUTIONS

A.K.W., Y.Y., and J.-M.G. designed the experiments. A.K.W., Y.Y., A.R., R.B.; M.S., J.B-B. performed the experiments. A.K.W., Y.Y., A.R., U.V. M.S., R.B., J.-M.B., C.B., D.L. and J.-M.G. analyzed the data. Y.Y., A.K.W. and J.-M.G. wrote the paper with significant contribution from all authors.

## DATA AVAILABILITY

The data that support the findings of this study are presented in the paper and/or the Supplementary Materials. Strains and plasmids are available from the corresponding author, JMG, upon reasonable request.

