## Supplementary material for "*Escherichia coli* SPFH membrane microdomain proteins HflKC contribute to aminoglycoside and oxidative stress tolerance": This pdf file includes supplementary methods, supplementary figures S1 to S7 and supplementaryTables S2 and S3.

Wessel AK, Yoshii Y *et al.*

This pdf file includes supplementary methods, supplementary figures S1 to S7 and supplementaryTables S2 and S3. Supplementary Table S1 is provided separately as a dataset.

### SUPPLEMENTARY METHODS

*Construction of YqiK tagged with GFP and HflK tagged with mCherry*

***YqiK-GFP chromosomal construct***

The *yqiK* (except for the stop codon), msf*gfp,* and *aphAIII* genes encoding YqiK, msfGFP, and KmFRT cassette were amplified from MG1655 *strep*, MG1655mreB-GFP (1), and MG1655Δ*cpxP*::KmFRT (2) chromosomes, respectively. Then, the intermediate plasmid pUC18 *yqiK-gfp-aphAIII* was constructed by combining these three DNA fragments with the linearized backbone vector pUC18, using Gibson assembly (New England Biolabs, Ipswich, MA, USA) and primers described in Table S3. Next, the DNA fragment encoding YqiK-GFP-KmFRT was amplified from the plasmid pUC18*yqiK-gfp-aphAIII* using the long recombining primers described in Table S3. The resulting PCR product was introduced on its native location under the *yqiK* promoter in strain MG1655 *strep* using pKOBEG plasmid by λ-red recombination to produce YqiK-GFP.

***HflK-mCherry chromosomal construct***

The DNA gene block corresponding to the entire *hflK* gene except for the stop codon, mCherry, and KmFRT with 40 bp-long homology regions of upstream of the *hflK* start codon and downstream of the *aphAIII* last codon was designed from the DNA sequence of MG1655 *strep*, MG1655mreB-mCherry (3), and MG1655Δ*cpxP*::KmFRT (2) chromosomes, respectively, and purchased from Eurofins Genomics (France). Then, the DNA oligonucleotide was introduced on its native location under the *hflK* promoter in *MG1655 strep* using pKOBEG plasmid by λ-red recombination to produce HflK-mCherry.

*Construction of QmcA-GFP and QmcA-derivatives tagged with GFP*

The *qmcA and ybbJ*, msf*gfp,* and *aphAIII* genes encoding QmcA and YbbJ, msfGFP and KmFRT cassette, respectively, were amplified from MG1655 *strep*, MG1655mreB-GFP, and MG1655Δ*cpxP*::KmFRT (2) chromosomes. The sequence of information on the *pf3* gene was obtained from (4).

• ***TM$^{QmcA}$-QmcA-GFP chromosomal construct***

The intermediate plasmid pUC18*qmcA-gfp-aphAIII* was constructed by combining DNA fragments of the entire *qmcA* gene except for the stop codon, msfGFP, the entire *ybbJ* gene followed by 20 bp upstream regions of the start codon, and KmFRT, with the linearized backbone vector pUC18 using Gibson assembly (New England Biolabs). The DNA fragment encoding QmcA-GFP-KmFRT was amplified from the plasmid pUC18*qmcA-gfp-aphAIII* using the long recombining primers described in Table S3, and introduced on its native location under the *qmcA* promoter in strain MG1655 *strep* using pKOBEG plasmid by λ-red recombination to produce TM$^{qmcA}$-QmcA-GFP.

***TM$^{QmcA}$-SPFH$^{QmcA}$-GFP chromosomal construct***

The DNA fragment encoding msfGFP-KmFRT with 40 bp-long upstream and downstream homology regions was amplified from the TM$^{pf3}$-SPFH$^{QmcA}$-GFP chromosomal construct using the long recombining primers described in Table S3. Then, the resulting PCR product was introduced on the chromosome at the downstream position of the *qmcA* SPFH domain region in strain MG1655 *strep* by λ-red recombination to produce TM$^{QmcA}$-SPFH$^{QmcA}$-GFP.

• ***TM$^{QmcA}$ -GFP chromosomal construct***

The DNA fragment encoding to msfGFP-KmFRT with 40 bp-long upstream and downstream homology regions was amplified from the TM$^{QmcA}$-QmcA-GFP chromosomal construct using long recombining primers described in Table S3. Then, the resulting PCR product was introduced on the chromosome at the downstream position of the *qmcA* TM domain region in strain MG1655 *strep* by λ-red recombination to produce TM$^{QmcA}$-GFP.

• **Δ*qmcA::zeo* chromosomal construct**

The *zeo* gene with 40 bp-long upstream and downstream homology regions was amplified from the MG1655_ λATT:*amp_GFPmut3_ΔfimAICDFGH::zeo ΔyadyadNecpDhtrEyadMLKC::cat* (5) using long recombining primers described in Table S3. Then, the resulting PCR product was introduced on the native location under the *qmcA* promoter in strain MG1655 strep by λ-red recombination to generate *ΔqmcA*::zeo.

• ***$TM^{pf3}$-QmcA-GFP chromosomal construct***

The plasmid pZS*12*$TM^{pf3}$-gfp-aphAIII* was constructed via pZS*12*$TM^{pf3}$-gfp,* by combining DNA fragments of *pf3*, msfGFP, and KmFRT, with the linearized backbone vector pZS*12 using Gibson assembly (New England Biolabs) and primers described in Table S3. Then, the intermediate plasmid pZS*12*$TM^{pf3}$-qmcA-gfp-aphAIII* was constructed by combining the DNA fragment of *qmcA* gene except for the native TM domain region with the linearized vector pZS*12*$TM^{pf3}$-gfp* using Gibson assembly (New England Biolabs). Next, the DNA fragment encoding $TM^{pf3}$-QmcA-GFP-KmFRT with 40 bp-long upstream and downstream homology regions was amplified from the intermediate plasmid using the long recombining primers described in Table S3. The resulting PCR product was introduced on its native location under *qmcA* promoter by λ-red recombination into strain Δ*qmcA*::*zeo* using pKOBEG plasmid to produce $TM^{pf3}$-QmcA-GFP.

• ***$TM^{pf3}$-$SPFH^{QmcA}$-GFP chromosomal construct***

The intermediate plasmid pZS*12*$TM^{pf3}$-$SPFH^{QmcA}$-gfp-aphAIII* was constructed by combining the SPFH domain region of *qmcA* gene with the linearized vector PZ*12 *$TM^{pf3}$-gfp-aphAIII* using Gibson assembly (New England Biolabs). Then, the DNA fragment encoding $TM^{pf3}$-$SPFH^{QmcA}$-GFP-KmFRT with 40 bp-long upstream and downstream homology regions was amplified from the plasmid pZS*12 *$TM^{pf3}$-$SPFH^{QmcA}$-GFP-aphAIII* using the long recombining primers described in Table S3. The resulting PCR product was introduced on its native location under *qmcA* promoter by λ-red recombination into Δ*qmcA*::*zeo* strain using pKOBEG plasmid to produce $TM^{pf3}$-$SPFH^{QmcA}$-GFP.

• ***$TM^{pf3}$-GFP chromosomal construct***

The DNA fragment encoding *$TM^{pf3}$*-GFP-KmFRT with 40 bp-long upstream and downstream homology regions was amplified from pZS*12*$TM^{pf3}$-gfp-aphAIII* using the long recombining primers described in Table S3. The resulting PCR product was introduced on its native location under *qmcA* promoter by λ-red recombination into Δ*qmcA*::*zeo* strain using pKOBEG plasmid to produce $TM^{pf3}$-GFP.

*Construction of HflC derivatives tagged with mCherry*

The *hflC*, *mCherry,* and *cat* genes encoding HflC, mCherry, and CmFRT cassette, respectively, were amplified from MG1655 *strep*, MG1655mreB-mCherry, and MG1655ΔyfcV-P::CmFRT (6) chromosomes. The sequence of information on *cmi* gene was obtained from (4).

• ***TM$^{HflC}$-HflC-mCherry chromosomal construct***

The intermediate plasmid pUC18*TM$^{hflC}$-HflC-mCherry-aphAIII* was constructed by combining DNA fragments of the *hflC* gene except for the stop codon, mCherry, and KmFRT, with the linearized backbone vector pUC18 using Gibson assembly (New England Biolabs) and primers described in Table S3. The DNA fragment encoding HflC-mCherry-KmFRT with 40 bp-long upstream and downstream homology regions was amplified from the plasmid pUC18*TM$^{hflC}$-HflC-mCherry-aphAIII* using the long recombining primers described in Table S3 and introduced on its native location under the *hflC* promoter in strain *MG1655 strep* using pKOBEG plasmid by λ-red recombination to TM$^{hflC}$-HflC-mCherry.

• ***TM$^{HflC}$- SPFH$^{HflC}$ -mCherry chromosomal construct***

The DNA fragment encoding mCherry-CmFRT with 40 bp-long upstream and downstream homology regions was amplified from the TM$^{cmi}$-SPFH$^{HflC}$-mCherry chromosomal construct, using the long recombining primers described in Table S3. The resulting PCR product was introduced on the chromosome at the downstream position of the *hflC* SPFH domain region by λ-red recombination into *MG1655 strep* strain using pKOBEGA plasmid to produce TM$^{hflC}$-SPFH$^{HflC}$-mCherry.

• ***TM$^{HflC}$-mCherry chromosomal construct***

The DNA fragment encoding mCherry-KmFRT with 40 bp-long upstream and downstream homology regions was amplified form the plasmid pUC18*TM$^{hflC}$-HflC-mCherry-aphAIII* using the long recombining primers described in Table S3. The resulting PCR product was introduced on the chromosome at the downstream position of the *hflC* TM domain region by λ-red recombination into *MG1655 strep* strain using pKOBEGA plasmid to produce TM$^{hflC}$-mCherry.

• ***TM$^{cmi}$-HflC-mCherry chromosomal construct***

The plasmid pZS*12*TM^cmi^-mCherry-cat* was constructed, via pZS*12*TM^cmi^-mCherry,* by combining DNA fragments encoding $TM^{cmi}$, mCherry, and CmFRT, with the linearized backbone vector pZS*12 using Gibson assembly (New England Biolabs) and the primers described in Table S3. Then, the intermediate plasmid pZS*12$TM^{cmi}$*-hflC-mCherry-cat* was constructed by combining the DNA fragment of *hflC* gene except for the native TM domain region with the linearized vector pZS*12$TM^{cmi}$*-mCherry-cat* using Gibson assembly (New England Biolabs). Then, the DNA fragment encoding $TM^{cmi}$-HflC-mCherry-CmFRT with 40 bp-long upstream and downstream homology regions was amplified from the intermediated plasmid using the long recombining primers described in Table S3. The resulting PCR product was introduced on its native location under *hflC* promoter by λ-red recombination into Δ*hflC*::KmFRT strain using pKOBEGA plasmid to produce $TM^{cmi}$-HflC-mCherry.

**• *$TM^{cmi}$- $SPFH^{HflC}$ -mCherry chromosomal construct***

The intermediated plasmid pZS*12$TM^{cmi}$*-*$SPFH^{HflC}$*-mCherry-cat* was constructed by combining the *hflC* SPFH domain region with the linearized initial plasmid pZS*12$TM^{cmi}$*-mCherry-cat*, which was used for generating $TM^{cmi}$-HflC-mCherry chromosomal construct, using Gibson assembly (New England Biolabs). Then, the DNA fragment encoding $TM^{cmi}$-$SPFH^{HflC}$-mCherry-CmFRT with 40 bp-long upstream and downstream homology regions was amplified from the intermediated plasmid using the long recombining primers described in Table S3. The resulting PCR product was introduced on its native location under *hflC* promoter by λ-red recombination into Δ*hflC*::KmFRT strain using pKOBEGA plasmid to produce $TM^{cmi}$-$SPFH^{HflC}$-mCherry.

**• *$TM^{cmi}$- mCherry chromosomal construct***

The DNA fragment encoding $TM^{cmi}$-GFP-CmFRT with 40 bp-long upstream and downstream homology regions was amplified from the plasmid pZS*12$TM^{cmi}$*-mCherry-cat* using the long recombining primers described in Table S3. The resulting PCR product was introduced on its native location under *hflC* promoter by λ-red recombination into Δ*hflC*::KmFRT strain using pKOBEGA plasmid to produce $TM^{cmi}$-mCherry.

*Construction of QmcA-GFP and HflC-mCherry in the ΔclsABC and Δidi mutants*

**• *QmcA-GFP chromosomal construct* in the Δ*clsABC* strain**

$TM^{QmcA}$-QmcA-GFP-KmFRT mutation from $TM^{QmcA}$-QmcA-GFP chromosomal construct was introduced into strain Δ*clsABC* by P1vir phage transduction (Δ*clsABC* QmcA-GFP).

**• *TM^QmcA^-QmcA-GFP chromosomal construct* in the Δ*idi* strain**

$TM^{QmcA}$-QmcA-GFP-KmFRT mutation from $TM^{QmcA}$-QmcA-GFP chromosomal construct was introduced into strain Δ*idi* by P1vir phage transduction (Δ*idi* QmcA-GFP).

**• *TM^HflC^-HflC-mCherry chromosomal construct* in the *ΔclsABC* strain**

$TM^{HflC}$-HflC-mCherry-KmFRT mutation from $TM^{HflC}$-HflC-mCherry chromosomal construct was introduced into strain Δ*clsABC* by P1vir phage transduction (Δ*clsABC* HflC-mCherry).

**• *TM^HflC^-HflC-mCherry chromosomal construct* in the Δ*idi* strain**

$TM^{HflC}$-HflC-mCherry-KmFRT mutation from $TM^{HflC}$-HflC-mCherry chromosomal construct was introduced into strain Δ*idi* by P1vir phage transduction (Δ*idi* HflC-mCherry).

**• *AcrA-GFP***

The DNA fragment encoding GFP-KmFRT region with 40 bp-long upstream and downstream homology regions was amplified from the $TM^{pf3}$-$SPFH^{QmcA}$-GFP chromosomal construct using the long recombining primers described in Table S3. The resulting PCR product was introduced on chromosome at the downstream position of the *acrA* gene (without stop codons) in *MG1655 strep* by λ-red recombination, to produce a GFP fusion. The mutation harboring KmFRT was introduced into a MG1655 *strep* strain having clean background by P1vir phage transduction (AcrA-GFP).

### SUPPLENTARY FIGURES

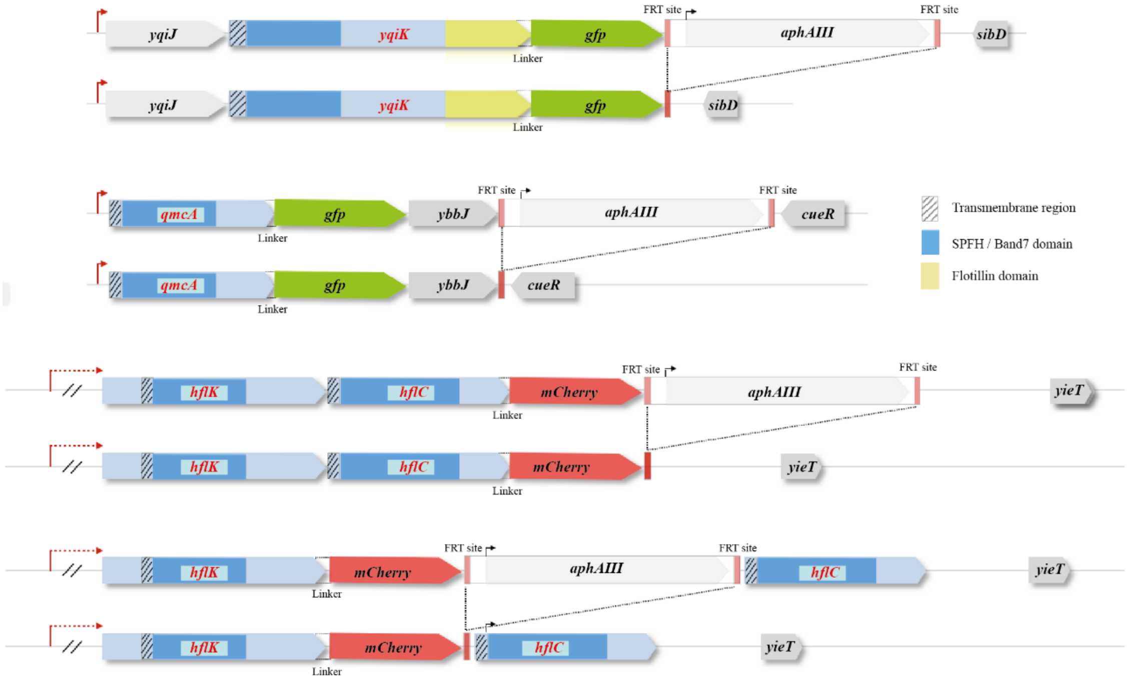

**Supplementary Figure S1. Schematic representation of chromosomal GFP or mCherry fusions to genes encoding *E. coli* SPFH proteins.** SPFH protein-encoding genes, *yqiK*, *qmcA*, are fused with monomeric super folder green fluorescent protein GFP (msfGFP) and *hflK*, and *hflC* to monomeric red fluorescent protein mCherry, respectively. Fusions were constructed as described in the Materials and Method section and the associated kanamycin antibiotic resistance marker (*aphAIII)* was inserted downstream of the fusion genes and removed using pCP20 plasmid, encoding the *flp* flippase.

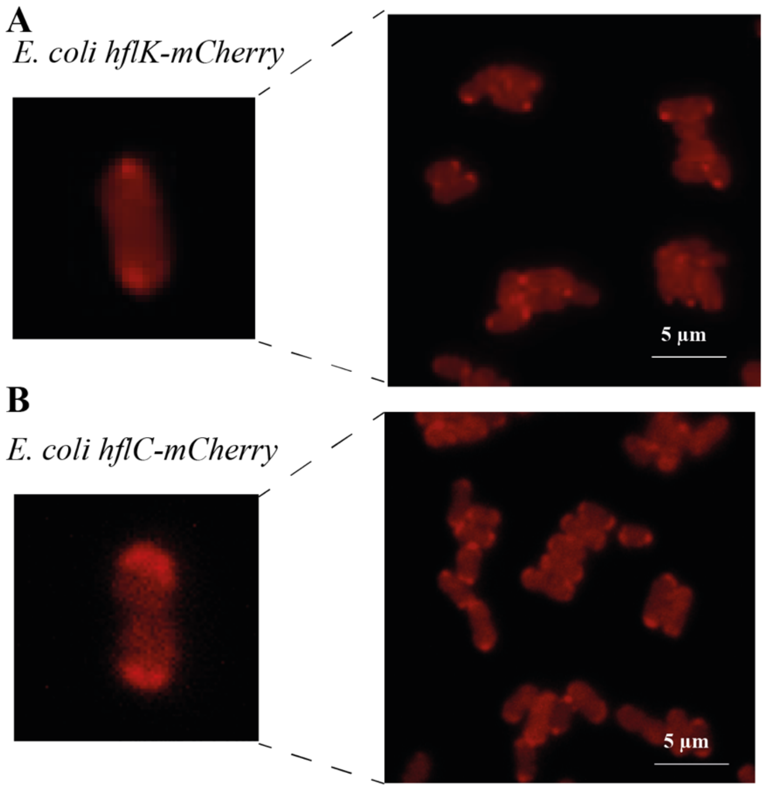

**Supplementary Figure S2. Localization pattern of HflK and HflC**. Epifluorescence microscopic images of cells expressing HflK-mCherry (A) and HflC-mCherry (B) in exponential phase with high and low magnifications.

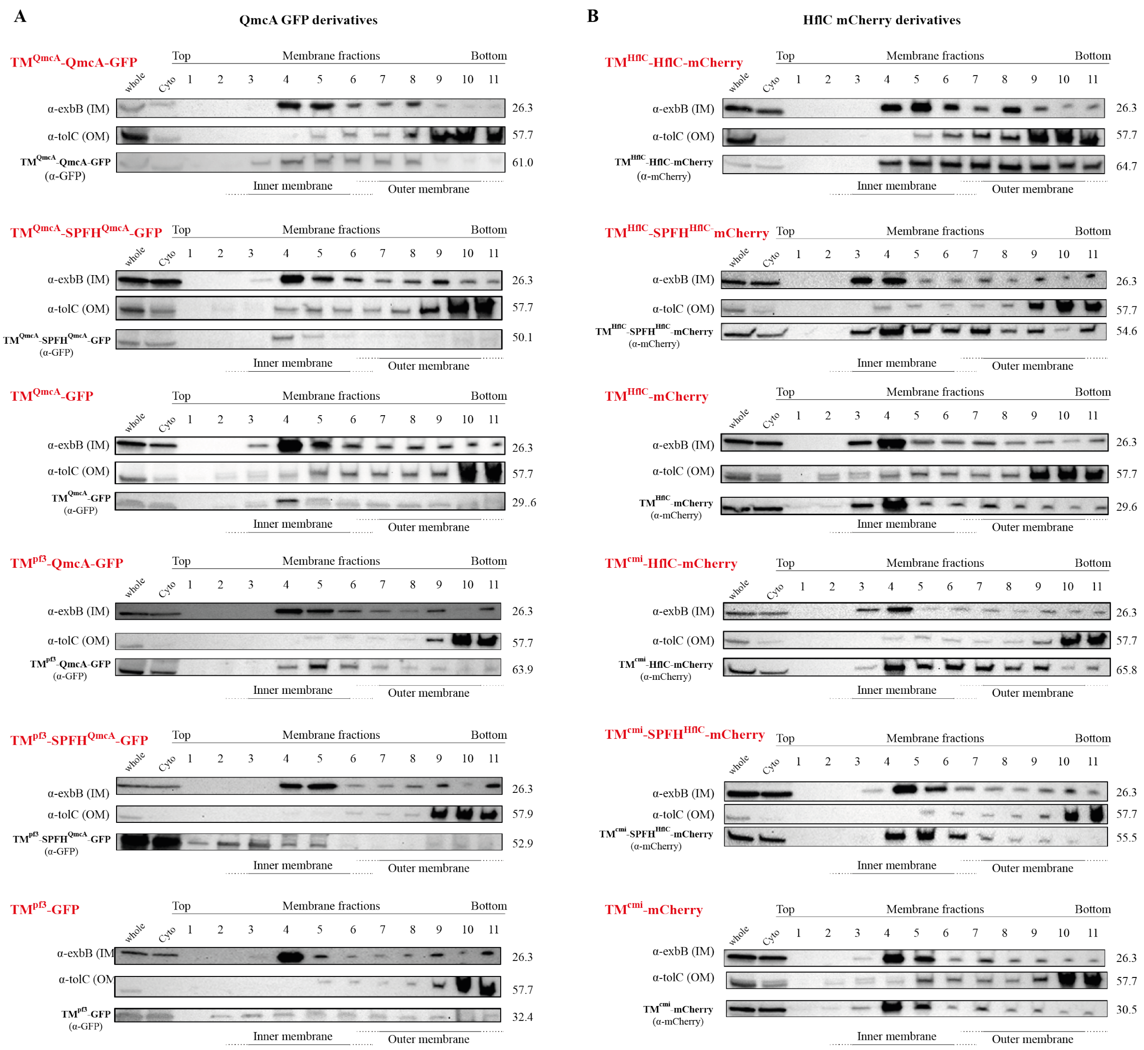

**Supplementary Figure S3. Inner membrane localization of QmcA and HflC in domain swap constructs.** SDS-PAGE and immunodetection analyses using proteins of whole-cell extract, cytosolic fraction, and membrane fractions prepared from cells expressing GFP fusion derivatives of QmcA (**A**) or mCherry fusion derivatives of HflC. (**B**). In immunodetection analysis, anti-ExbB antibodies were used to detect the inner membrane- (IM) marker ExbB and anti-TolC antibodies to detect the outer membrane- (OM) marker TolC. Anti-GFP and anti-mCherry antibodies were used to detect the expressions of GFP and mCherry fused proteins, respectively.

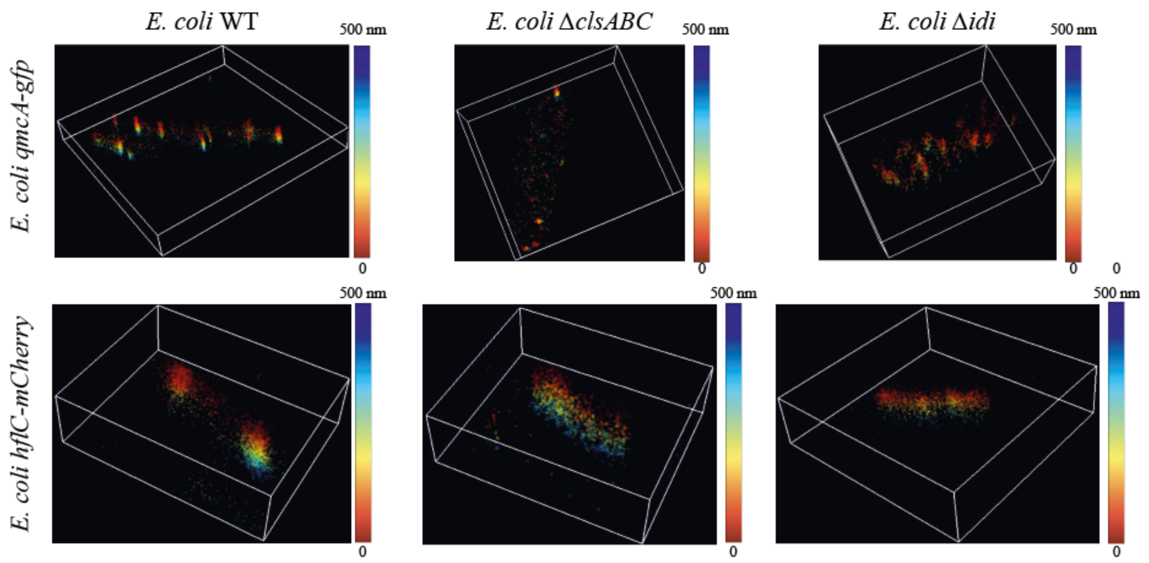

**Supplementary Figure S4. The subcellular localization of QmcA and HflC in mutants with the alteration of cardiolipin or isoprenoid synthetic pathway.** Three-dimensional super-resolution microscopy images of WT, Δ*clsABC*, and Δ*idi* strains expressing QmcA-GFP and HflC-mCherry in stationary phase, with colors corresponding to depth location along the Z axis, 0-500 nm, with 0 nm expressed in red, and 500 nm expressed in deep blue.

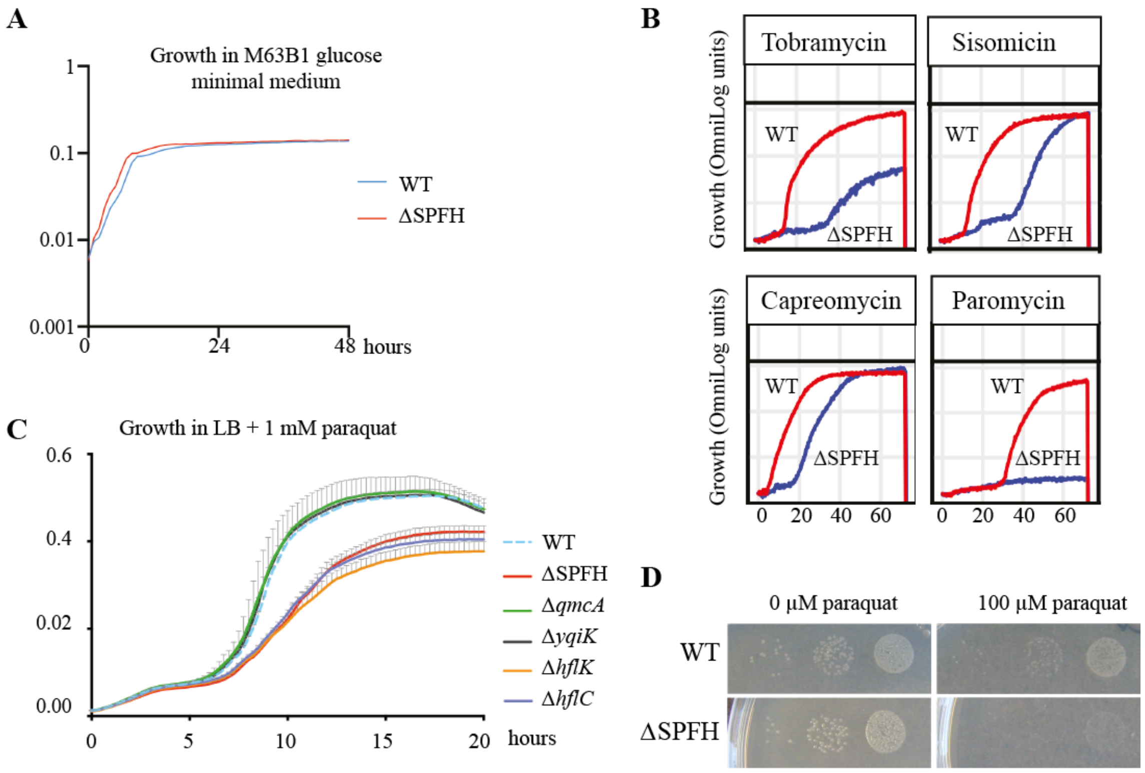

**Supplementary Figure S5. Phenotypic analysis of *E. coli* SPFH mutant**

(**A**) Growth curve of *E. coli* WT and ΔSPFH mutant in M63B1glucose minimal medium. (**B**) Biolog metabolic activity curve of WT and ΔSPFH in presence of tobramycin, capreomycin, sisomicin, and paromycin. (**C**) Bacterial growth curve of WT and ΔSPFH in the presence of 0.8 mM paraquat. (**D**) Paraquat susceptibility assay of WT and ΔSPFH spotted on M63B1glucose agar containing 100 μM paraquat. (**E**) Growth curve of WT and indicated SPFH gene mutants in the presence of 0.8 mM paraquat.

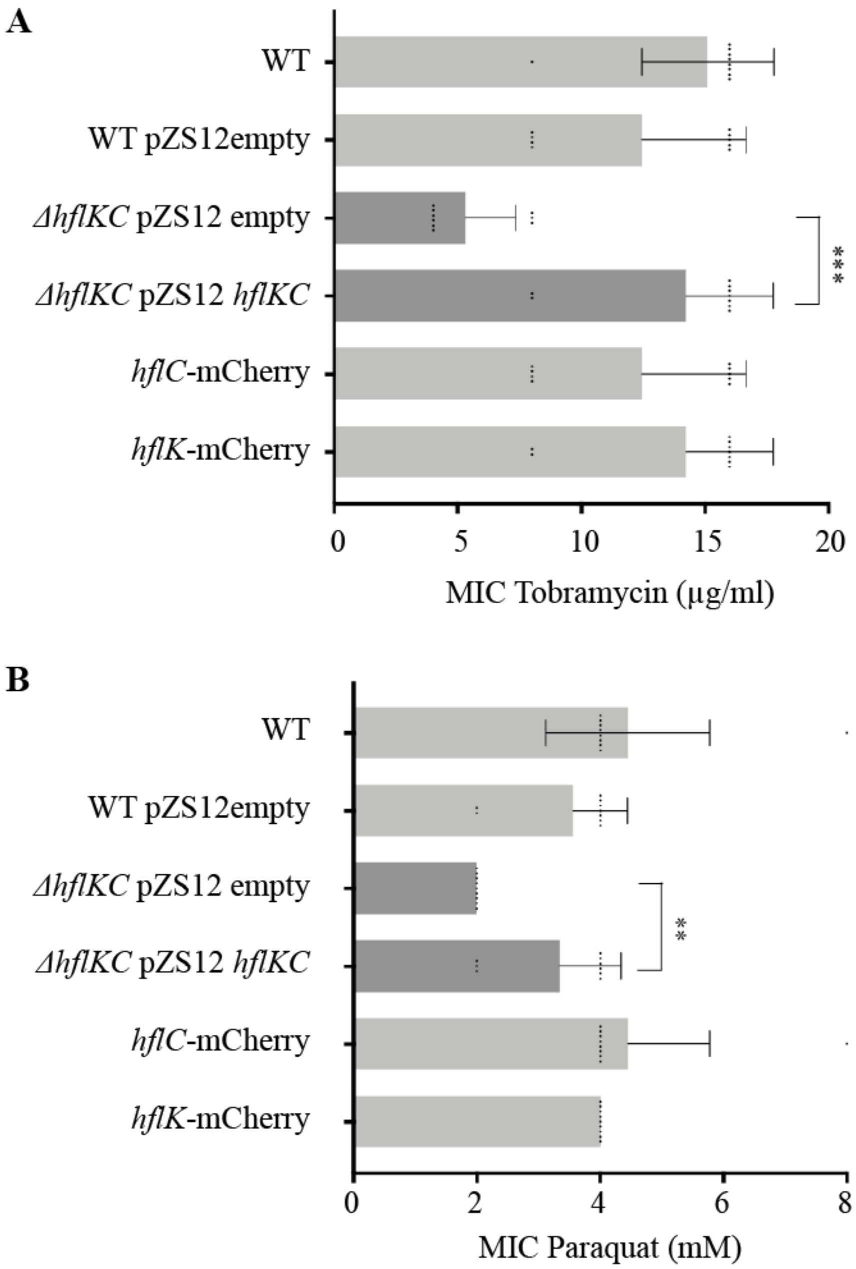

**Supplementary Figure S6. Susceptibility to tobramycin and paraquat of complemented mutants**. Minimum inhibitory concentrations (MICs) for tobramycin (**A**) and paraquat (**B**) of *E. coli* Wt, Δ*hflKC* strains harboring empty vector and *hflKC*- complemented strains or strain expressing *hflC or hflK-mCherry constructs* from their native chromosomal context. $^{**}p < 0.01$, $^{***}p < 0.001$.

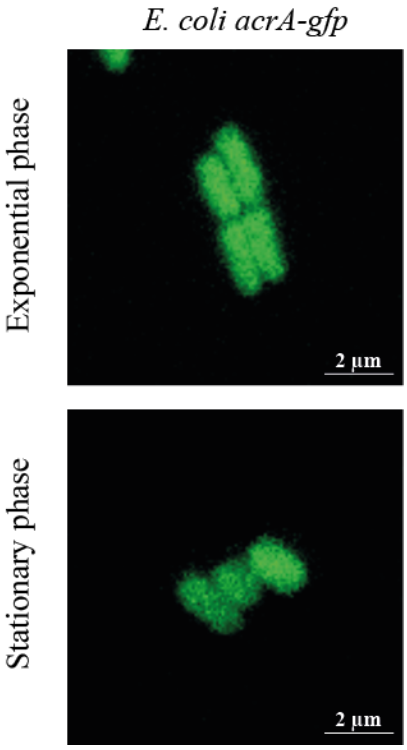

**Supplementary Figure S7. Localization of AcrA-GFP fusion protein does not show polar localization pattern when expressed from native chromosomal context.** Epifluorescence microscopic images of cells expressing AcrA-GFP in exponential and stationary phases.

### Supplementary Table S2. Bacterial strains and plasmids used in this study

| *E. coli* strains and plasmids | Genotype or description | Antibiotic resistance(s) | Reference |
|---|---|---|---|
| **Strains** | | | |
| BW25113Δ*yqiK*::Kmfrt | Keio collection | $Km^R$ | (1) |
| BW25113Δ*qmcA*::Kmfrt | Keio collection | $Km^R$ | (1) |
| BW25113Δ*hflK*::Kmfrt | Keio collection | $Km^R$ | (1) |
| BW25113Δ*hflC*::Kmfrt | Keio collection | $Km^R$ | (1) |
| BW25113Δ*clsA*::Kmfrt | Keio collection | $Km^R$ | (1) |
| BW25113Δ*clsB*::Kmfrt | Keio collection | $Km^R$ | (1) |
| BW25113Δ*clsC*::Kmfrt | Keio collection | $Km^R$ | (1) |
| BW25113Δ*idi*::Kmfrt | Keio collection | $Km^R$ | (1) |
| BW25113Δ*acrA*::Kmfrt | Keio collection | $Km^R$ | (1) |
| MG1655 *strep* | Spontaneous streptomycin-resistant mutant of K-12 wild-type strain | $Strep^R$ | (2) |
| MG1655mreB-GFP | Source of msf*gfp* gene | No resistance | Gift from Dr. Sven van Teeffelen |
| MG1655mreB-mCherry | Source of *mCherry* gene | No resistance | Gift from Dr. Sven van Teeffelen |
| MG1655Δ*cpxP*::KmFRT | Source of kanamycine resistance cassette | $Km^R$ | (3) |
| MG1655ΔyfcV-P::CmFRT | Source of chloramphenicol resistance cassette | $Cm^R$ | (4) |
| DH5alpir pSW25 ptetAccdB | Source of spectinomycin resistance cassette | $Spec^R$ | (5) |

| | | | |
|---|---|---|---|
| MG1655_λATT:*amp_GFPmut3_ΔfimAICDFGH*::zeo Δ*yadyadNecpDhtrEyadMLKC*::*cat* | Source of zeocin resistance cassette | $Zeo^R$ | (6) |
| Δ*yqiK* | P1 vir transduction from BW25113Δ*yqiK*::Kmfrt into MG1655 *strep* | $Strep^R$ | This study |
| Δ*qmcA* | P1 vir transduction from BW25113Δ*qmcA*::Kmfrt into MG1655 *strep* | $Strep^R$ | This study |
| Δ*hflK* | P1 vir transduction from BW25113Δ*hflK*::Kmfrt into MG1655 *strep* | $Strep^R$ | This study |
| Δ*hflC* | P1 vir transduction from BW25113Δ*hflC*::Kmfrt into MG1655 *strep* | $Strep^R$ | This study |
| Δ*hflKC* | Double mutant of Δ*hflK* and Δ*hflC*. MG1655*strepR*, Δ*hlfKC*::Cmfrt | $Strep^R$ | This study |
| Δ*yqiJK* | Double mutant of Δ*yqiJ* and Δ*yqiK*. MG1655*strepR*, Δ*yqiJK*::Spec | $Strep^R$, $Spec^R$ | This study |
| Δ*qmcA*, Δ*yqiJK* | Triple mutant of Δ*qmcA* and Δ*yqiJK*. MG1655*strepR*, Δ*qmcA,* Δ*yqiJK*::Spec | $Strep^R$, $Spec^R$ | This study |
| Δ*SPFH* | Quintuple mutant of Δ*yqiK*, Δ*qmcA*, Δ*hflK*, and Δ*hflC*. MG1655*strepR*, Δ*qmcA*, Δ*yqiJK*::Spec, Δ*hlfKC*::Cmfrt | $Strep^R$, $Spec^R$ | This study |
| Δ*clsA* | P1 vir transduction from BW25113Δ*clsA*::Kmfrt into MG1655 *strep* | $Strep^R$ | This study |
| Δ*clsB* | P1 vir transduction from BW25113Δ*clsB*::Kmfrt into MG1655 *strep* | $Strep^R$ | This study |
| Δ*clsC* | P1 vir transduction from BW25113Δ*clsC*::Kmfrt into MG1655 *strep* | $Strep^R$ | This study |

| | | | |
|---|---|---|---|
| Δ*clsAB* | Double mutant of Δ*clsA* and Δ*clsB*. P1 vir transduction from BW25113Δ*clsA*::Kmfrt into Δ*clsB* | $Strep^R$ | This study |
| Δ*clsABC* | Triple mutant of Δ*clsA*, Δ*clsB*, and Δ*clsC*. P1 vir transduction from BW25113Δ*clsC*::Kmfrt into Δ*clsAB* | $Strep^R$ | This study |
| Δ*idi* | P1 vir transduction from BW25113Δ*idi*::Kmfrt into MG1655 *strep* | $Strep^R$ | This study |
| Δ*acrA* | P1 vir transduction from BW25113Δ*acrA*::Kmfrt into MG1655 *strep* | $Strep^R$ | This study |
| Δ*qmcA*::*zeo* | MG1655 *strep,* Δ*qmcA*::*zeo* | $Strep^R$, $Zeo^R$ | This study |
| *yqiK-gfp* | MG1655 *strep, yqiK*::*gfp* Kmfrt | $Strep^R$, $Km^R$ | This study |
| *qmcA-gfp* | MG1655 *strep, qmcA*::*gfp* Kmfrt | $Strep^R$, $Km^R$ | This study |
| *hflK-mCherry* | MG1655 *strep, hflK*::*mCherry* Kmfrt | $Strep^R$, $Km^R$ | This study |
| *hflC-mCherry* | MG1655 *strep, hflC*::*mCherry* Kmfrt | $Strep^R$, $Km^R$ | This study |
| $TM^{qmcA}$-$SPFH^{qmcA}$-*gfp* | MG1655 *strep,* Δ*qmcA*::$TM^{qmcA}$-$SPFH^{qmcA}$-*gfp* Kmfrt | $Strep^R$, $Km^R$ | This study |
| $TM^{qmcA}$-*gfp* | MG1655 *strep,* Δ*qmcA*::$TM^{qmcA}$-*gfp* Kmfrt | $Strep^R$, $Km^R$ | This study |
| $TM^{Pf3}$-*qmcA-gfp* | MG1655 *strep,* Δ*qmcA*::$TM^{pf3}$-*qmcA-gfp* Kmfrt | $Strep^R$, $Km^R$ | This study |
| $TM^{Pf3}$-$SPFH^{qmcA}$-*gfp* | MG1655 *strep,* Δ*qmcA*::$TM^{pf3}$-$SPFH^{qmcA}$-*gfp* Kmfrt | $Strep^R$, $Km^R$ | This study |
| $TM^{Pf3}$-*gfp* | MG1655 *strep,* Δ*qmcA*::$TM^{pf3}$-*gfp* Kmfrt | $Strep^R$, $Km^R$ | This study |
| $TM^{hflC}$-$SPFH^{hflC}$-*mCherry* | MG1655 *strep,* ΔhflC::$TM^{hflC}$-$SPFH^{hflC}$-*mCherry* Cmfrt | $Strep^R$, $Cm^R$ | This study |
| $TM^{hflC}$-*mCherry* | MG1655 *strep,* ΔhflC::$TM^{hflC}$-*mCherry* Kmfrt | $Strep^R$, $Km^R$ | This study |
| $TM^{cmi}$-*hflC-mCherry* | MG1655 *strep,* Δ*hflC*::$TM^{cmi}$-*hflC- mCherry* Cmfrt | $Strep^R$, $Cm^R$ | This study |
| $TM^{cmi}$-$SPFH^{hflC}$-*mCherry* | MG1655 *strep,* Δ*hflC*::$TM^{Cmi}$ -$SPFH^{hflC-}$*mCherry* Cmfrt | $Strep^R$, $Cm^R$ | This study |
| $TM^{cmi}$-*mCherry* | MG1655 *strep,* Δ*hflC*::$TM^{Cmi}$-*mCherry* Cmfrt | $Strep^R$, $Cm^R$ | This study |

| | | | |
|---|---|---|---|
| Δ*clsABC qmcA-gfp* | P1 vir transduction from *qmcA-gfp* into Δ*clsABC* | $Strep^R$, $Km^R$ | This study |
| Δ*idi qmcA-gfp* | P1 vir transduction from *qmcA-gfp* into Δ*idi* | $Strep^R$, $Km^R$ | This study |
| Δ*clsABC hflC-mCherry* | P1 vir transduction from *hflC-mCherry* into Δ*clsABC* | $Strep^R$, $Km^R$ | This study |
| Δ*idi hflC-mCherry* | P1 vir transduction from *hflC-mCherry* into Δ*idi* | $Strep^R$, $Km^R$ | This study |
| *acrA-gfp* | MG1655 *strep, acrA*::*gfp* Kmfrt | $Strep^R$, $Km^R$ | This study |
| **Plasmids** | | | |
| pKOBEG | *ori*R101*ts araC* arabinose-inducible λred γβα operon | $Cm^R$ | (7) |
| pKOBEGA | pKOBEG derivative | $Amp^R$ | (7) |
| pCP20 | Rep(Ts) $Flp^+$ | $Cm^R$, $Amp^R$ | (8) |
| pUC18 | High-copy-number, lac promoter cloning vector, ampicillin resistant (AmpR) | $Amp^R$ | (9) |
| pZS*12 | Tightly regulated $P_{LtetO-1}$ Low copy-number vector ampicillin resistant (AmpR) | $Amp^R$ | (10) |
| pUC18y*qiK-gfp-aphAIII* | Assembly of *YqiK-mCherry-aphAIII* into pUC18 | $Amp^R$, $Km^R$ | This study |
| pUC18$TM^{QmcA}$-*QmcA-gfp-aphAIII* | Assembly of $TM^{QmcA}$-*QmcA-gfp-aphAIII* into pUC18 | $Amp^R$, $Km^R$ | This study |
| pZS*12$TM^{pf3}$-*gfp* | Assembly of $TM^{pf3}$-*gfp* into pZS*12 | $Amp^R$ | This study |
| pZS*12$TM^{pf3}$-*gfp-aphAIII* | Assembly of *aphAIII* into PZ*12$TM^{pf3}$-*gfp* | $Amp^R$, $Km^R$ | This study |
| pZS*12$TM^{pf3}$-*QmcA-gfp-aphAIII* | Assembly of *qmcA* into PZ*12$TM^{pf3}$-*gfp-aphAIII* | $Amp^R$, $Km^R$ | This study |
| pZS*12 $TM^{pf3}$-$SPFH^{QmcA}$-*gfp-aphAIII* | Assembly of $SPFH^{qmcA}$ into PZ*12$TM^{pf3}$-*gfp-aphAIII* | $Amp^R$, $Km^R$ | This study |
| pUC18$TM^{HflC}$-*HflC-mCherry-aphAIII* | Assembly of $TM^{HflC}$-*HflC-mCherry-aphAIII* into pUC18 | $Amp^R$, $Km^R$ | This study |
| pZS*12$TM^{cmi}$-*mCherry* | Assembly of *cmi-mCherry* into pZS*12 | $Amp^R$ | This study |
| pZS*12$TM^{cmi}$-*mCherry-cat* | Assembly of *Cmfrt* into PZ*12 *cmi-mCherry* | $Amp^R$, $Cm^R$ | This study |

| | | | |
|---|---|---|---|
| pZS*12*TM$^{cmi}$-hflC-mCherry-cat* | Assembly of *hflC* into PZ*12 *cmi-mCherry-Cmfrt* | Amp$^R$, Cm$^R$ | This study |
| pZS*12*TM$^{cmi}$-SPFH$^{HflC}$-mCherry-cat* | Assembly of *SPFH$^{HflC}$* into PZ*12 *cmi-mCherry-Cmfrt* | Amp$^R$, Cm$^R$ | This study |
| pZS*12-*HflKC* | Assembly of *hflKC* into pZS*12 | Amp$^R$ | This study |

### Supplementary Table S3. Primers used for generating chromosomal mutants or assembling plasmids in this study

| Purpose | Name | Sequence (5’-3’) |
|---|---|---|
| For amplifying the DNA fragment to generate Δ*hflKC* strain | Up.ΔhflKC-CmFRT-5 | gacaacagggatcaccgcataacaaatatggagcacaaacgtgtaggctggagctgcttcgaag |
| | Down.ΔhflKC-CmFRT-3 | ggatgcggtggctttattgacctgtaccgcagtcgttatacatatgaatatcctccttagttcc |
| For amplifying the DNA fragment to generate Δ*yqiJK* strain | Up.ΔyqiJK-Spec-5 | caaagtaaatattgcgtcactaaatggacattggagtgatatgcgtcaattgcgccgaataaatac |
| | Down.ΔyqiJK-Spec-3 | ccccggctttttgggccagggattattcctggaagagggttcattggctggcaccaagcagttt |
| For amplifying the DNA fragment to generate Δ*qmcA*::*zeo* strain | Up.ΔqmcA-Zeo-5 | caattttgctgattatggacggtacaacaggaggtttttccgttctggatccgagcacgtgttgac |
| | Down.ΔqmcA-Zeo-3 | gactgagccagaaaatatgtggatgaacgaccattaactccatctcagtcctgctcctcggccacg |
| For assembling the pUC18y*qiK-gfp-aphAIII* plasmid | Up.GFP_pUC18yqiK-gfp-5 | caccccttaacctcaacaactcccgtcgaagaaaaagcagagggcggcggcggcagcatgagtaaaggtgaagaac |
| | Down.GFP_pUC18yqiK-gfp-3 | caaaatctttctgtcgttatttgtagagttcatccatg |
| | Up.KmFRT_pUC18yqiK-gfp-5 | cgacagaaagattttgggagcaaatgatgtgtgtaggc |
| | Down.KmFRT_pUC18yqiK-gfp-3 | ccccggctttttgggccagggattattcctggaagagggtcagggtttactcatatgaatatc |
| | Up.linearized pUC18_pUC18yqiK-gfp-5 | gcccaaaaagccgggggaattcgtaatcatggtcatag |
| | Down.linearized pUC18_pUC18yqiK-gfp-5 | gttgaggttaagggtgggatcctctagagtcgacctg |
| For amplifying the DNA fragment to generate YqiK-GFP strain | Up.yqiK_YqiK-GFP-KmFRT-5 | Same as Up.GFP_pUC18yqiK-gfp-5 |
| | Down.KmFRT_YqiK-GFP-KmFRT-3 | Same as Down.KmFRT_pUC18yqiK-gfp-3 |
| For assembling the pUC18TM$^{QmcA}$-*QmcA-gfp-aphAIII* plasmid | Up.qmcA-gfp_pUC18qmcA-gfp-5 | ccgagctggtgaaagacagcgccaacaaacggactcagccaggcggcggcggcagcatgagtaaaggtgaagaactgttc |
| | Down.qmcA-gfp_pUC18qmcA-gfp-3 | ggctgagtccgtttgttggcttatttgtagagttcatccatgccgtgcgtgatacctgctg |
| | Up.ybbJ_pUC18qmcA-gfp-5 | cagcaggtatcacgcacggcatggatgaactctacaaataagccaacaaacggactcagcc |
| | Down.ybbJ_pUC18qmcA-gfp-3 | caaaatctttctgtcgttaagacgagacggctcggatatg |
| | Up.KmFRT_pUC18qmcA-gfp-5 | cgacagaaagattttgggagcaaatgatgtgtgtaggc |

| | | |
|---|---|---|
| | Down.KmFRT_pUC18qmcA-gfp-3 | gaaaatctctccggctgctgtcatcatcgggcagggtgatcagggtttactcatatgaatatc |
| | Up.linearized pUC18_pUC18qmcA-gfp-5 | cagccggagagattttcgaattcgtaatcatggtcatag |
| | Down.linearized pUC18_pUC18qmcA-gfp-5 | ctttcaccagctcggggatcctctagagtcgacctg |
| For amplifying the DNA fragment to generate TM$^{qmcA}$-QmcA-GFP strain | Up.qmcA_QmcA-GFP-KmFRT-5 | Same as Up.qmcA-gfp_pUC18qmcA-gfp-5 |
| | Down.KmFRT_QmcA-GFP-KmFRT-3 | Down.KmFRT_pUC18qmcA-gfp-3 |
| For amplifying the DNA fragment to generate TM$^{qmcA}$-SPFH$^{qmcA}$-GFP strain | UP.SPFHqmcA_SPFHqmcA-GFP-KmFRT-5 | gcggaagggatccgtcaggcggaaatcctcaaagccgaaggtgaatttaacaacaacatgag taaagg |
| | Down.KmFRT_SPFHqmcA-GFP-KmFRT-3 | gaaaatctctccggctgctgtcatcatcgggcagggtgatcagggtttactcatatgaatatc |
| For amplifying the DNA fragment to generate TM$^{qmcA}$-GFP strain | Up.TMqmcA_TMqmcA-GFP-KmFRT-5 | ctcatttttgtcgcgctggtcattgtcggcgcgggtgtcaaaatcgtagaatttaacaacaacatg agtaaagg |
| | Down.KmFRT_TMqmcA-GFP-KmFRT-3 | Same sequence as Down.KmFRT_SPFHqmcA-GFP-KmFRT-3 |
| For assembling the pZS*12*TM$^{pf3}$-gfp* plasmid | Up.pf3-GFP_pZS*12pf3-gfp-5 | gtgagcggataacaagatacatgcaatccgtgattactg |
| | Down.pf3-GFP_pZS*12pf3-gfp-3 | ctttactcatgttgttgttaaattcaaagaattg |
| | Up.GFP _pZS*12pf3-gfp-5 | taacaacaacatgagtaaaggtgaagaac |
| | Down.GFP_pZS*12pf3-gfp-3 | tctagactcagctaattaagtcatttgtagagttcatcc |
| | Up.linearized pZS*12 pZS*12pf3-gfp-5 | ctacaaatgacttaattagctgagtctagag |
| | Down.linearized pZS*12pZS*12pf3-gfp-3 | cggattgcatgtatcttgttatccgctc |
| For assembling the pZS*12*TM$^{pf3}$-gfp-aphAIII* plasmid | Up.KmFRT _pZS*12pf3-gfp-KmFRT-5 | cgaaaggctcagtcgaacgacagaaagattttgggag |
| | Down.KmFRT_pZS*12pf3-gfp-KmFRT-3 | cgaaaggcccagtcttcagggtttactcatatgaatatcc |
| | Up. linearized pZS*12_pZS*12pf3-gfp-KmFRT-5 | ggatattcatatgagtaaaccctgaagactgggcctttcg |
| | Down. linearized pZS*12_pZS*12pf3-gfp-KmFRT-3 | ctttctgtcgttcgactgagcctttcgttttatttg |
| For assembling the pZS*12*TM$^{pf3}$-qmcA-gfp-aphAIII* plasmid | Up. qmcA_pZS*12pf3-qmcA-gfp-KmFRT-5 | tggatcaaagcgcaattctttatcgtaccgcagggctatc |
| | Down. qmcA_pZS*12pf3-qmcA-gfp-KmFRT-3 | actcatgttgttgttaaattctggctgagtccgtttgttggc |

| | | |
|---|---|---|
| | Up. linearized pZS*12_pZS*12pf3-gfp-Kmfrt-5 | ggatattcatatgagtaaaccctgaagactgggcctttcg |
| | Down. linearized pZS*12_pZS*12pf3-gfp-Kmfrt-3 | ggtacgataaagaattgcgctttgatcc |
| For assembling the pZS*12*TM^pf3^-SPFH^QmcA^ -gfp-aphAIII* plasmid | Up. SPFHqmcA_pZS*12pf3-SPFHqmcA-gfp-KmFRT-5 | Same as Up. qmcA_pZS*12pf3-qmcA-gfp-KmFRT-5 |
| | Down. SPFHqmcA_pZS*12pf3-SPFHqmcA-gfp-KmFRT-3 | actcatgttgttgttaaattcaccttcggctttgaggatttcc |
| | Up. Linearised pZS*12_pZS*12pf3-SPFHqmcA-gfp-KmFRT-5 | caaagccgaaggtgaatttaacaacaacatgagtaaagg |
| | Down. Linearised pZS*12_pZS*12pf3-SPFHqmcA-gfp-KmFRT-3 | Same as Down. linearized pZS*12_pZS*12pf3-gfp-Kmfrt-3 |
| For amplifying the DNA fragments to generate $TM^{Pf3}$-QmcA-GFP, $TM^{Pf3}$-$SPFH^{QmcA}$-GFP, and $TM^{Pf3}$-GFP strains | Up.Pf3_pf3-(QmcA)-GFP-KmFRT-5 | caattttgctgattatggacggtacaacaggaggtttttccgatgcaatccgtgattactgatg |
| | Down.KmFRT_pf3-(QmcA)-GFP-KmFRT-3 | Same sequence as Down.KmFRT_SPFHqmcA-GFP-KmFRT-3 |
| For assembling the pUC18*$TM^{HflC}$-HflC-mCherry-aphAIII* plasmid | Up.hflC-mCherry_pUC18hflC-mCherry-KmFRT-5 | gatttcttccgctacatgaagacgccgacttccgcaacgcgtggcggcggcggcagcatggtttccaagggcgaggag |
| | Down. hflC-mCherry_pUC18hflC-mCherry-KmFRT-3 | caaaatctttctgtcgttatttgtacagctcatccatg |
| | Up.KmFRT_pUC18hflC-mCherry-KmFRT-5 | cgacagaaagattttgggagcaaatgatgtgtgtaggc |
| | Down.KmFRT_pUC18hflC-mCherry-KmFRT-3 | gaggatgcggtggctttattgacctgtaccgcagtcgttatatcagggtttactcatatgaatatc |
| | Up.linearized pUC18_ pUC18hflC-mCherry-KmFRT-5 | caataaagccaccgcatcctcgaattcgtaatcatggtcatag |
| | Down.linearized pUC18_ pUC18hflC-mCherry-KmFRT-3 | catgtagcggaagaaatcggatcctctagagtcgacctg |
| For amplifying the DNA fragment to generate $TM^{HflC}$-HflC-mCherry strain | Up.hflC_HflC-mCherry-KmFRT-5 | Same as Up.hflC-mCherry_pUC18hflC-mCherry-5 |
| | Down.KmFRT_HflC-mCherry-KmFRT-3 | gaggatgcggtggctttattgacctgtaccgcagtcgttatatcagggtttactcatatgaatatc |
| For amplifying the DNA fragment to generate $TM^{hflC}$-$SPFH^{hflC}$-mCherry strain | Up.SPFHhflC_SPFHhflC-mCherry-CmFRT-5 | gagcgtgaagcggtagcgcgtcgtcaccgttcacaaggtcaggaatttaacaacaacatggtttccaagg |
| | Down.CmFRT_SPFHhflC-mCherry-CmFRT-3 | ggatgcggtggctttattgacctgtaccgcagtcgttatacatatgaatatcctccttagttcctattcc |
| For amplifying the DNA fragment to generate $TM^{hflC}$-mCherry strain | Up.TMhflC_TMhflC-mCherry-KmFRT-5 | catcgtgctggtagtgctttacatgtctgtctttgtcgtcggcggcggcggcagcatg |
| | Down.KmFRT_TMhflC-mCherry-KmFRT-3 | gaggatgcggtggctttattgacctgtaccgcagtcgttatatcagggtttactcatatgaatatc |
| | Up.cmi-pZS*12cmi-mcherry-5 | gaaaggtaccatgaaagtgattagcatgaaatttatttttattc |

| | | |
|---|---|---|
| For assembling the pZS*12$TM^{cmi}$-*mCherry* plasmid | Down.cmi-pZS*12cmi-mCherry-3 | tggaaaccatgttgttgttaaattcgcc |
| | Up.mCherry_pZS*12cmi-mCherry-5 | taacaacaacatggtttccaagggcgag |
| | Down.mCherry_pZS*12cmi-mCherry-3 | ctaattaagcttatttgtacagctcatccatgc |
| | Up.linearized pZS*12_pZS12cmi-mCherry-5 | gtacaaataagcttaattagctgagtctagagg |
| | Down.linearized pZS*12_pZS12cmi-mCherry-3 | tcactttcatggtacctttctcctctttaatg |
| For assembling the pZS*12$TM^{cmi}$-*mCherry-cat* plasmid | Up.CmFRT_pZS*12cmi-mCherry-CmFRT-5 | gctcagtcgagtgtaggctggagctgcttc |
| | Down. CmFRT_pZS*12cmi-mCherry-CmFRT-5 | gcccagtcttcatatgaatatcctccttagttcctattcc |
| | Up.linearized pZS*12_ pZS*12cmi-mCherry-CmFRT-5 | tattcatatgaagactgggcctttcgttttatc |
| | Down.linearized pZS*12_ pZS*12cmi-mCherry-CmFRT-3 | cagcctacactcgactgagcctttcgttttatttg |
| For assembling the pZS*12$TM^{cmi}$-*hflC-mCherry-cat* plasmid | Up.hflC_ pZS*12cmi-mCherry-CmFRT-5 | agataaaggcgtcgtcaaagaaggtgagcgc |
| | Down.hflC_ pZS*12cmi-mCherry-CmFRT-3 | tgttaaattcacgcgttgcggaagtcgg |
| | Up.linearized pZS*12_ pZS*12cmi-hflC-mCherry-CmFRT-5 | cgcaacgcgtgaatttaacaacaacatggtttcc |
| | Down.linearized pZS*12_ pZS*12cmi-hflC-mCherry-CmFRT-3 | ctttgacgacgcctttatcttcgctccaaaaaaac |
| For assembling the pZS*12$TM^{cmi}$-$SPFH^{HflC}$-*mCherry-cat* plasmid | Up. SPFHhflC_ pZS*12cmi-SPFHhflC-mCherry-CmFRT-5 | Same as Up.hflC_ pZS*12cmi-mCherry-CmFRT-5 |
| | Down. SPFHhflC_ pZS*12cmi-SPFHhflC-mCherry-CmFRT-3 | tgttaaattcctgaccttgtgaacggtg |
| | Up.linearized pZS*12_ pZS*12cmi-SPFHhflC-mCherry-CmFRT-5 | acaaggtcaggaatttaacaacaacatggtttcc |
| | Down.linearized pZS*12_ pZS*12cmi-SPFHhflC-mCherry-CmFRT-5 | Same as Down.linearized pZS*12_ pZS*12cmi-hflC-mCherry-CmFRT-3 |
| For amplifying the DNA fragments to generate $TM^{cmi}$-HflC-mCherry, $TM^{cmi}$-$SPFH^{HflC}$-mCherry, $TM^{cmi}$-mCherry | Up.cmi_cmi-(HflC)-mCherry-CmFRT-5 | ccaacgcgcagcgtaacgactaccagcgtcagggggaataacgatgaaagtgattagcatgaa |
| | Down.CmFRT_cmi-(HflC)-mCherry-CmFRT-3 | ggatgcggtggctttattgacctgtaccgcagtcgttatacatatgaatatcctccttag |
| For amplifying the DNA fragment to generate AcrA-GFP strain | Up.acrA_AcrA-GFP-KmFRT-5 | ccagcaagccgcaagcggtgctcagcctgaacagtccaagtctgaatttaacaacaacatgagtaaaggtgaagaac |
| | Down.KmFRT_AcrA-GFP-KmFRT-3 | cgggcgatcgataaagaaattaggcatgtcttaacggctcctgtttaagtcagggtttactcatatgaatatcctcc |

| | | |
|---|---|---|
| For assembling the pZS12**hflKC* plasmid | Up.hflKC_ pZS*12-HflKC-5 | cattaaagaggagaaaggtaccatggcgtggaatcagcccggtaataacgg |
| | Down.hflKC_ pZS*12-HflKC -3 | gatgcctctagactcagctaattaagcttttaacgcgttgcggaagtcggcg |
| | Up.linearized pZS*12_ pZS*12 -5 | aagcttaattagctgagtctag |
| | Down.linearized pZS*12_ pZS*12-3 | ggtacctttctcctctttaatgaattcgg |
