## Supplementary Table S1 for "*Escherichia coli* SPFH membrane microdomain proteins HflKC contribute to aminoglycoside and oxidative stress tolerance"

BIOLOG

### Phenotype MicroArrays™

#### PM1 MicroPlate™ Carbon Sources

| | | | | | | | | | | | |
|---|---|---|---|---|---|---|---|---|---|---|---|
| A1 Negative Control | A2 L-Arabinose | A3 N-Acetyl-D-Glucosamine | A4 D-Saccharic Acid | A5 Succinic Acid | A6 D-Galactose | A7 L-Aspartic Acid | A8 L-Proline | A9 D-Alanine | A10 D-Trehalose | A11 D-Mannose | A12 Dulcitol |
| B1 D-Serine | B2 D-Sorbitol | B3 Glycerol | B4 L-Fucose | B5 D-Glucuronic Acid | B6 D-Gluconic Acid | B7 D,L-α-Glycerol-Phosphate | B8 D-Xylose | B9 L-Lactic Acid | B10 Formic Acid | B11 D-Mannitol | B12 L-Glutamic Acid |
| C1 D-Glucose-6-Phosphate | C2 D-Galactonic Acid-γ-Lactone | C3 D,L-Malic Acid | C4 D-Ribose | C5 Tween 20 | C6 L-Rhamnose | C7 D-Fructose | C8 Acetic Acid | C9 α-D-Glucose | C10 Maltose | C11 D-Melibiose | C12 Thymidine |
| D-1 L-Asparagine | D2 D-Aspartic Acid | D3 D-Glucosaminic Acid | D4 1,2-Propanediol | D5 Tween 40 | D6 α-Keto-Glutaric Acid | D7 α-Keto-Butyric Acid | D8 α-Methyl-D-Galactoside | D9 α-D-Lactose | D10 Lactulose | D11 Sucrose | D12 Uridine |
| E1 L-Glutamine | E2 M-Tartaric Acid | E3 D-Glucose-1-Phosphate | E4 D-Fructose-6-Phosphate | E5 Tween 80 | E6 α-Hydroxy Glutaric Acid-γ-Lactone | E7 α-Hydroxy Butyric Acid | E8 β-Methyl-D-Glucoside | E9 Adonitol | E10 Maltotriose | E11 2-Deoxy Adenosine | E12 Adenosine |
| F1 Glycyl-L-Aspartic Acid | F2 Citric Acid | F3 M-Inositol | F4 D-Threonine | F5 Fumaric Acid | F6 Bromo Succinic Acid | F7 Propionic Acid | F8 Mucic Acid | F9 Glycolic Acid | F10 Glyoxylic Acid | F11 D-Cellobiose | F12 Inosine |
| G1 Glycyl-L-Glutamic Acid | G2 Tricarballylic Acid | G3 L-Serine | G4 L-Threonine | G5 L-Alanine | G6 L-Alanyl-Glycine | G7 Acetoacetic Acid | G8 N-Acetyl-β-D-Mannosamine | G9 Mono Methyl Succinate | G10 Methyl Pyruvate | G11 D-Malic Acid | G12 L-Malic Acid |
| H1 Glycyl-L-Proline | H2 p-Hydroxy Phenyl Acetic Acid | H3 m-Hydroxy Phenyl Acetic Acid | H4 Tyramine | H5 D-Psicose | H6 L-Lyxose | H7 Glucuronamide | H8 Pyruvic Acid | H9 L-Galactonic Acid-γ-Lactone | H10 D-Galacturonic Acid | H11 Phenylethyl-amine | H12 2-Aminoethanol |

#### PM2A MicroPlate™ Carbon Sources

| | | | | | | | | | | | |
|---|---|---|---|---|---|---|---|---|---|---|---|
| A1 Negative Control | A2 Chondroitin Sulfate C | A3 α-Cyclodextrin | A4 β-Cyclodextrin | A5 γ-Cyclodextrin | A6 Dextrin | A7 Gelatin | A8 Glycogen | A9 Inulin | A10 Laminarin | A11 Mannan | A12 Pectin |
| B1 N-Acetyl-D-Galactosamine | B2 N-Acetyl-Neuraminic Acid | B3 β-D-Allose | B4 Amygdalin | B5 D-Arabinose | B6 D-Arabitol | B7 L-Arabitol | B8 Arbutin | B9 2-Deoxy-D-Ribose | B10 I-Erythritol | B11 D-Fucose | B12 3-0-β-D-Galacto-pyranosyl-D-Arabinose |
| C1 Gentiobiose | C2 L-Glucose | C3 Lactitol | C4 D-Melezitose | C5 Maltitol | C6 α-Methyl-D-Glucoside | C7 β-Methyl-D-Galactoside | C8 3-Methyl Glucose | C9 β-Methyl-D-Glucuronic Acid | C10 α-Methyl-D-Mannoside | C11 β-Methyl-D-Xyloside | C12 Palatinose |
| D1 D-Raffinose | D2 Salicin | D3 Sedoheptulosan | D4 L-Sorbose | D5 Stachyose | D6 D-Tagatose | D7 Turanose | D8 Xylitol | D9 N-Acetyl-D-Glucosaminitol | D10 γ-Amino Butyric Acid | D11 δ-Amino Valeric Acid | D12 Butyric Acid |
| E1 Capric Acid | E2 Caproic Acid | E3 Citraconic Acid | E4 Citramalic Acid | E5 D-Glucosamine | E6 2-Hydroxy Benzoic Acid | E7 4-Hydroxy Benzoic Acid | E8 β-Hydroxy Butyric Acid | E9 γ-Hydroxy Butyric Acid | E10 α-Keto Valeric Acid | E11 Itaconic Acid | E12 5-Keto-D-Gluconic Acid |
| F1 D-Lactic Acid Methyl Ester | F2 Malonic Acid | F3 Melibionic Acid | F4 Oxalic Acid | F5 Oxalomalic Acid | F6 Quinic Acid | F7 D-Ribono-1,4-Lactone | F8 Sebacic Acid | F9 Sorbic Acid | F10 Succinamic Acid | F11 D-Tartaric Acid | F12 L-Tartaric Acid |
| G1 Acetamide | G2 L-Alaninamide | G3 N-Acetyl-L-Glutamic Acid | G4 L-Arginine | G5 Glycine | G6 L-Histidine | G7 L-Homoserine | G8 Hydroxy-L-Proline | G9 L-Isoleucine | G10 L-Leucine | G11 L-Lysine | G12 L-Methionine |
| H1 L-Ornithine | H2 L-Phenylalanine | H3 L-Pyroglutamic Acid | H4 L-Valine | H5 D,L-Carnitine | H6 Sec-Butylamine | H7 D.L-Octopamine | H8 Putrescine | H9 Dihydroxy Acetone | H10 2,3-Butanediol | H11 2,3-Butanone | H12 3-Hydroxy 2-Butanone |

#### PM01 (Carbon Sources)

ΔSPFH-1
WT−1

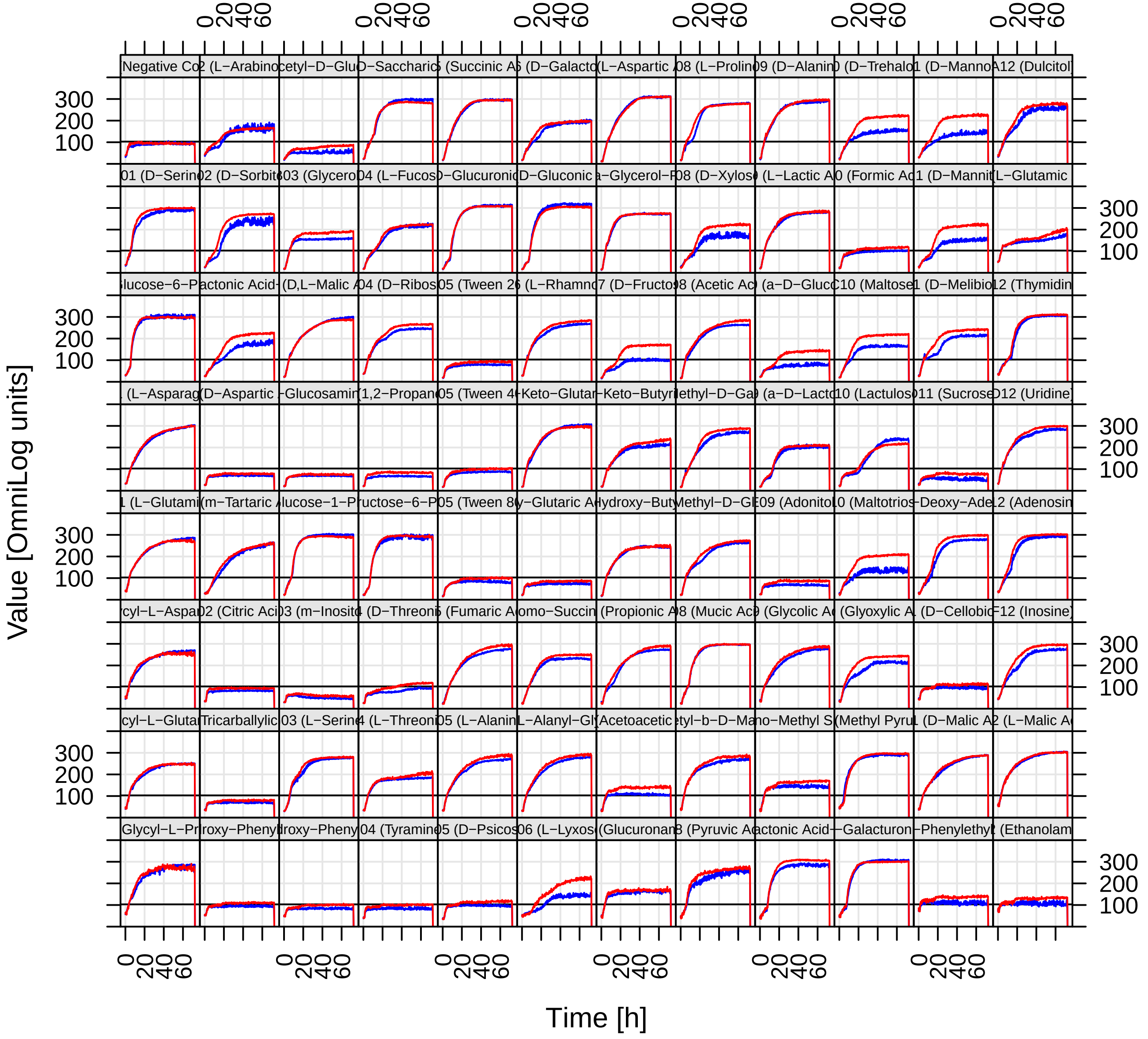

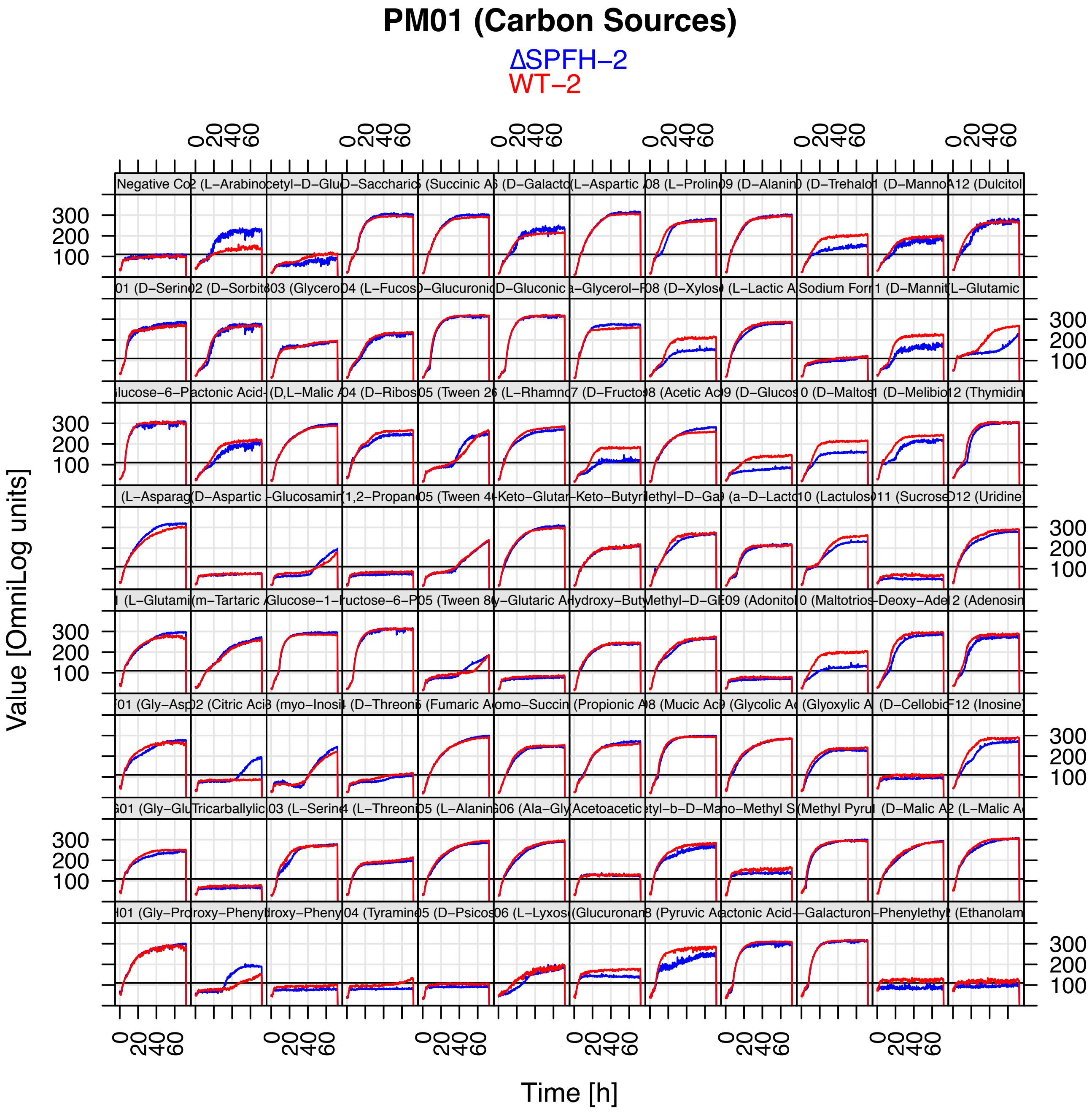

PM01 (Carbon Sources)
ΔSPFH−2
WT−2
Value [OmniLog units]
Time [h]

### PM02 (Carbon Sources)

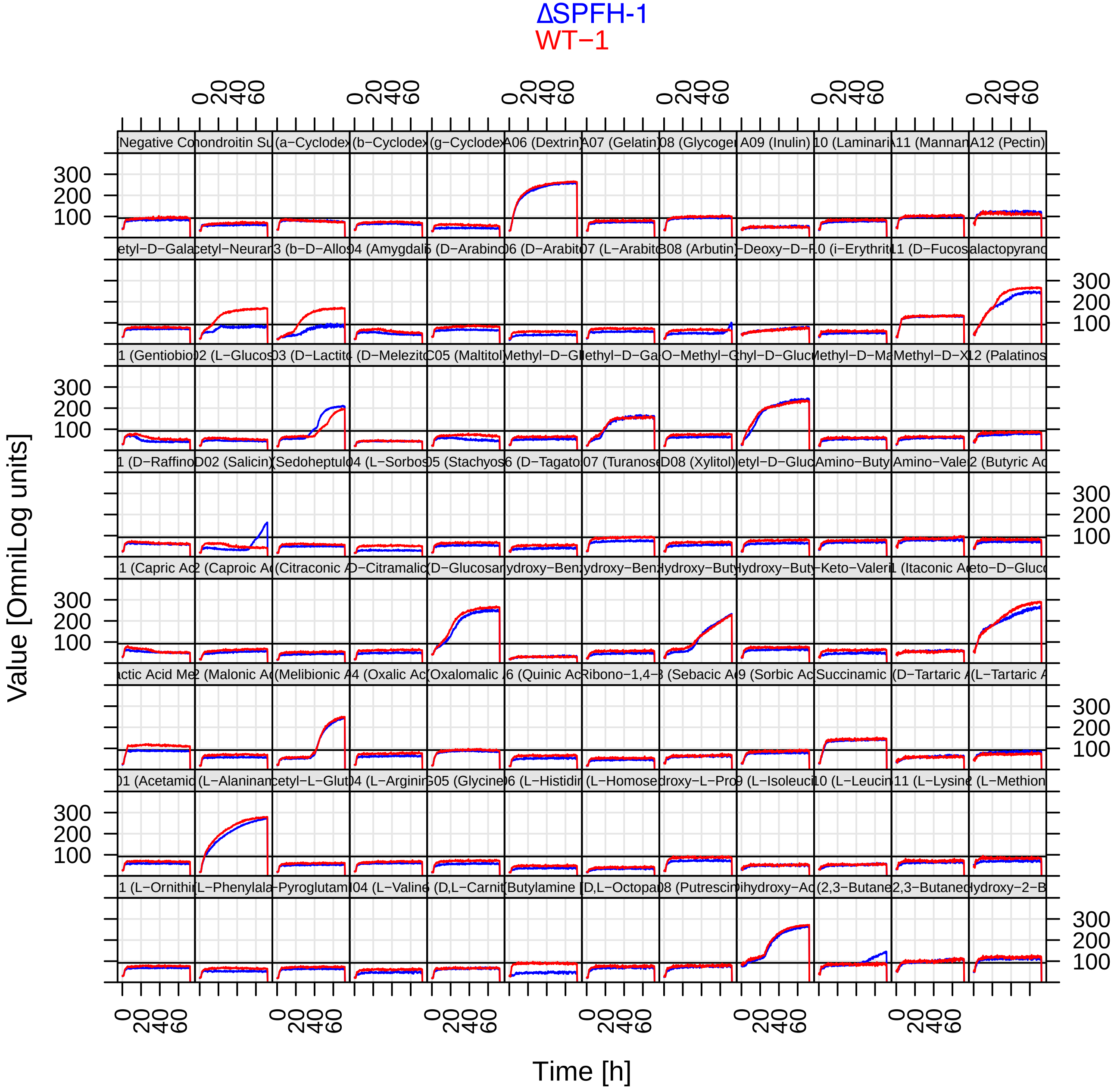

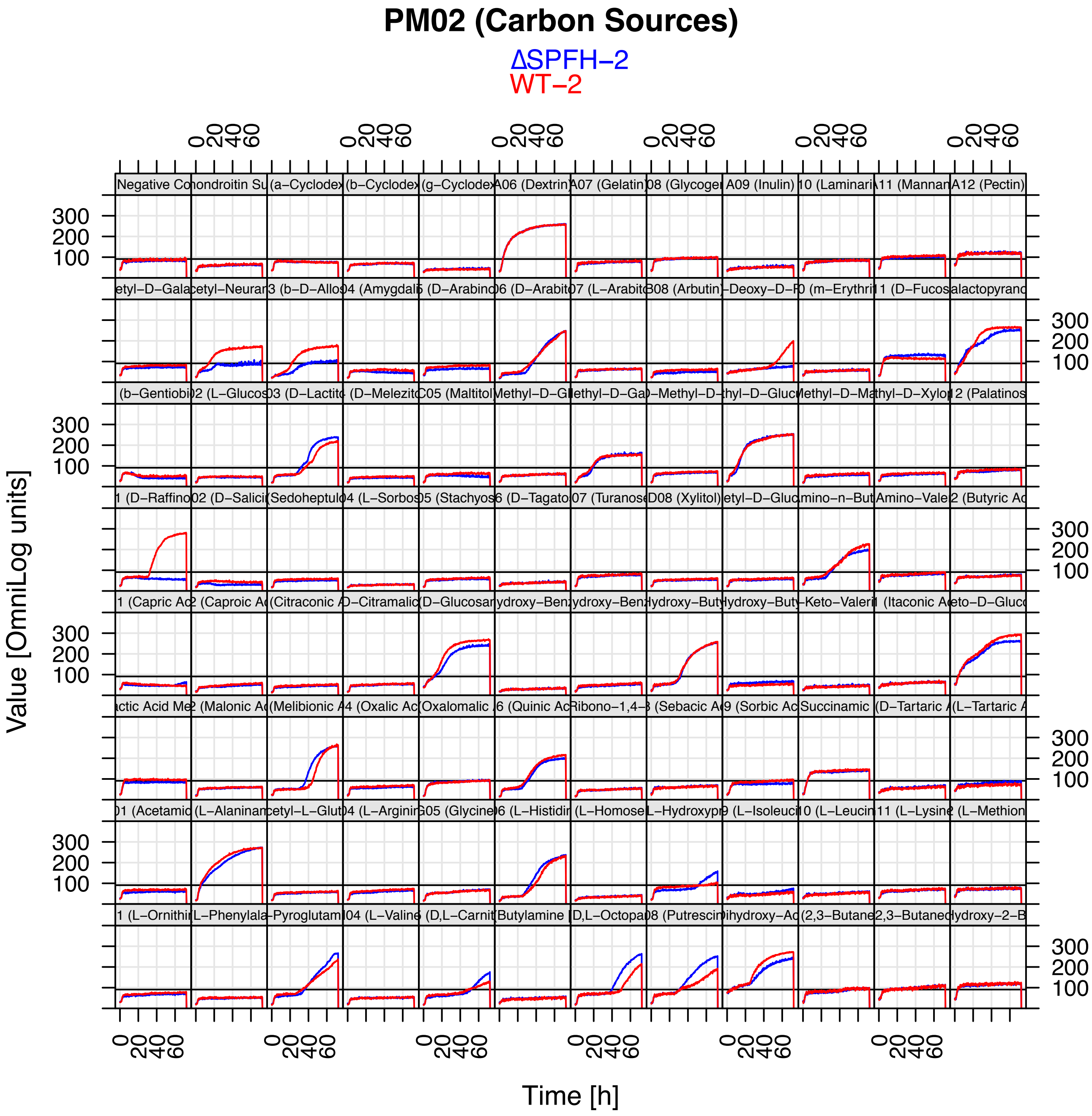
PM02 (Carbon Sources)
ΔSPFH−2
WT−2
Value [OmniLog units]
Time [h]

### Phenotype MicroArrays™

#### PM3B MicroPlate™ Nitrogen Sources

| | | | | | | | | | | | |
|---|---|---|---|---|---|---|---|---|---|---|---|
| A1 Negative Control | A2 Ammonia | A3 Nitrite | A4 Nitrate | A5 Urea | A6 Biuret | A7 L-Alanine | A8 L-Arginine | A9 L-Asparagine | A10 L-Aspartic Acid | A11 L-Cysteine | A12 L-Glutamic Acid |
| B1 L-Glutamine | B2 Glycine | B3 L-Histidine | B4 L-Isoleucine | B5 L-Leucine | B6 L-Lysine | B7 L-Methionine | B8 L-Phenylalanine | B9 L-Proline | B10 L-Serine | B11 L-Threonine | B12 L-Tryptophan |
| C1 L-Tyrosine | C2 L-Valine | C3 D-Alanine | C4 D-Asparagine | C5 D-Aspartic Acid | C6 D-Glutamic Acid | C7 D-Lysine | C8 D-Serine | C9 D-Valine | C10 L-Citrulline | C11 L-Homoserine | C12 L-Ornithine |
| D-1 N-Acetyl-D,L-Glutamic Acid | D2 N-Phthaloyl-L-Glutamic Acid | D3 L-Pyroglutamic Acid | D4 Hydroxylamine | D5 Methylamine | D6 N-Amylamine | D7 N-Butylamine | D8 Ethylamine | D9 Ethanolamine | D10 Ethylenediamine | D11 Putrescine | D12 Agmatine |
| E1 Histamine | E2 β-Phenylethyl-amine | E3 Tyramine | E4 Acetamide | E5 Formamide | E6 Glucuronamide | E7 D,L-Lactamide | E8 D-Glucosamine | E9 D-Galactosamine | E10 D-Mannosamine | E11 N-Acetyl-D-Glucosamine | E12 N-Acetyl-D-Galactosamine |
| F1 N-Acetyl-D-Mannosamine | F2 Adenine | F3 Adenosine | F4 Cytidine | F5 Cytosine | F6 Guanine | F7 Guanosine | F8 Thymine | F9 Thymidine | F10 Uracil | F11 Uridine | F12 Inosine |
| G1 Xanthine | G2 Xanthosine | G3 Uric Acid | G4 Alloxan | G5 Allantoin | G6 Parabanic Acid | G7 D,L-α-Amino-N-Butyric Acid | G8 γ-Amino-N-Butyric Acid | G9 ε-Amino-N-Caproic Acid | G10 D,L-α-Amino-Caprylic Acid | G11 δ-Amino-N-Valeric Acid | G12 α-Amino-N-Valeric Acid |
| H1 Ala-Asp | H2 Ala-Gln | H3 Ala-Glu | H4 Ala-Gly | H5 Ala-His | H6 Ala-Leu | H7 Ala-Thr | H8 Gly-Asn | H9 Gly-Gln | H10 Gly-Glu | H11 Gly-Met | H12 Met-Ala |

#### PM4A MicroPlate™ Phosphorus and Sulfur Sources

| | | | | | | | | | | | |
|---|---|---|---|---|---|---|---|---|---|---|---|
| A1 Negative Control | A2 Phosphate | A3 Pyrophosphate | A4 Trimeta-phosphate | A5 Tripoly-phosphate | A6 Triethyl Phosphate | A7 Hypophosphite | A8 Adenosine- 2'-monophosphate | A9 Adenosine- 3'-monophosphate | A10 Adenosine- 5'-monophosphate | A11 Adenosine-2',3'-cyclic monophosphate | A12 Adenosine-3',5'-cyclic monophosphate |
| B1 Thiophosphate | B2 Dithiophosphate | B3 D,L-α-Glycerol Phosphate | B4 β-Glycerol Phosphate | B5 Carbamyl Phosphate | B6 D-2-Phospho-Glyceric Acid | B7 D-3-Phospho-Glyceric Acid | B8 Guanosine- 2'-monophosphate | B9 Guanosine- 3'-monophosphate | B10 Guanosine- 5'-monophosphate | B11 Guanosine-2',3'-cyclic monophosphate | B12 Guanosine-3',5'-cyclic monophosphate |
| C1 Phosphoenol Pyruvate | C2 Phospho-Glycolic Acid | C3 D-Glucose-1-Phosphate | C4 D-Glucose-6-Phosphate | C5 2-Deoxy-D-Glucose 6-Phosphate | C6 D-Glucosamine-6-Phosphate | C7 6-Phospho-Gluconic Acid | C8 Cytidine- 2'-monophosphate | C9 Cytidine- 3'-monophosphate | C10 Cytidine- 5'-monophosphate | C11 Cytidine- 2',3'-cyclic monophosphate | C12 Cytidine- 3',5'-cyclic monophosphate |
| D1 D-Mannose-1-Phosphate | D2 D-Mannose-6-Phosphate | D3 Cysteamine-S-Phosphate | D4 Phospho-L-Arginine | D5 O-Phospho-D-Serine | D6 O-Phospho-L-Serine | D7 O-Phospho-L-Threonine | D8 Uridine- 2'-monophosphate | D9 Uridine- 3'-monophosphate | D10 Uridine- 5'-monophosphate | D11 Uridine- 2',3'-cyclic monophosphate | D12 Uridine- 3',5'-cyclic monophosphate |
| E1 O-Phospho-D-Tyrosine | E2 O-Phospho-L-Tyrosine | E3 Phosphocreatine | E4 Phosphoryl Choline | E5 O-Phosphoryl-Ethanolamine | E6 Phosphono Acetic Acid | E7 2-Aminoethyl Phosphonic Acid | E8 Methylene Diphosphonic Acid | E9 Thymidine- 3'-monophosphate | E10 Thymidine- 5'-monophosphate | E11 Inositol Hexaphosphate | E12 Thymidine 3',5'- cyclic monophosphate |
| F1 Negative Control | F2 Sulfate | F3 Thiosulfate | F4 Tetrathionate | F5 Thiophosphate | F6 Dithiophosphate | F7 L-Cysteine | F8 D-Cysteine | F9 L-Cysteinyl-Glycine | F10 L-Cysteic Acid | F11 Cysteamine | F12 L-Cysteine Sulfinic Acid |
| G1 N-Acetyl-L-Cysteine | G2 S-Methyl-L-Cysteine | G3 Cystathionine | G4 Lanthionine | G5 Glutathione | G6 D,L-Ethionine | G7 L-Methionine | G8 D-Methionine | G9 Glycyl-L-Methionine | G10 N-Acetyl-D,L-Methionine | G11 L- Methionine Sulfoxide | G12 L-Methionine Sulfone |
| H1 L-Djenkolic Acid | H2 Thiourea | H3 1-Thio-β-D-Glucose | H4 D,L-Lipoamide | H5 Taurocholic Acid | H6 Taurine | H7 Hypotaurine | H8 p-Amino Benzene Sulfonic Acid | H9 Butane Sulfonic Acid | H10 2-Hydroxyethane Sulfonic Acid | H11 Methane Sulfonic Acid | H12 Tetramethylene Sulfone |

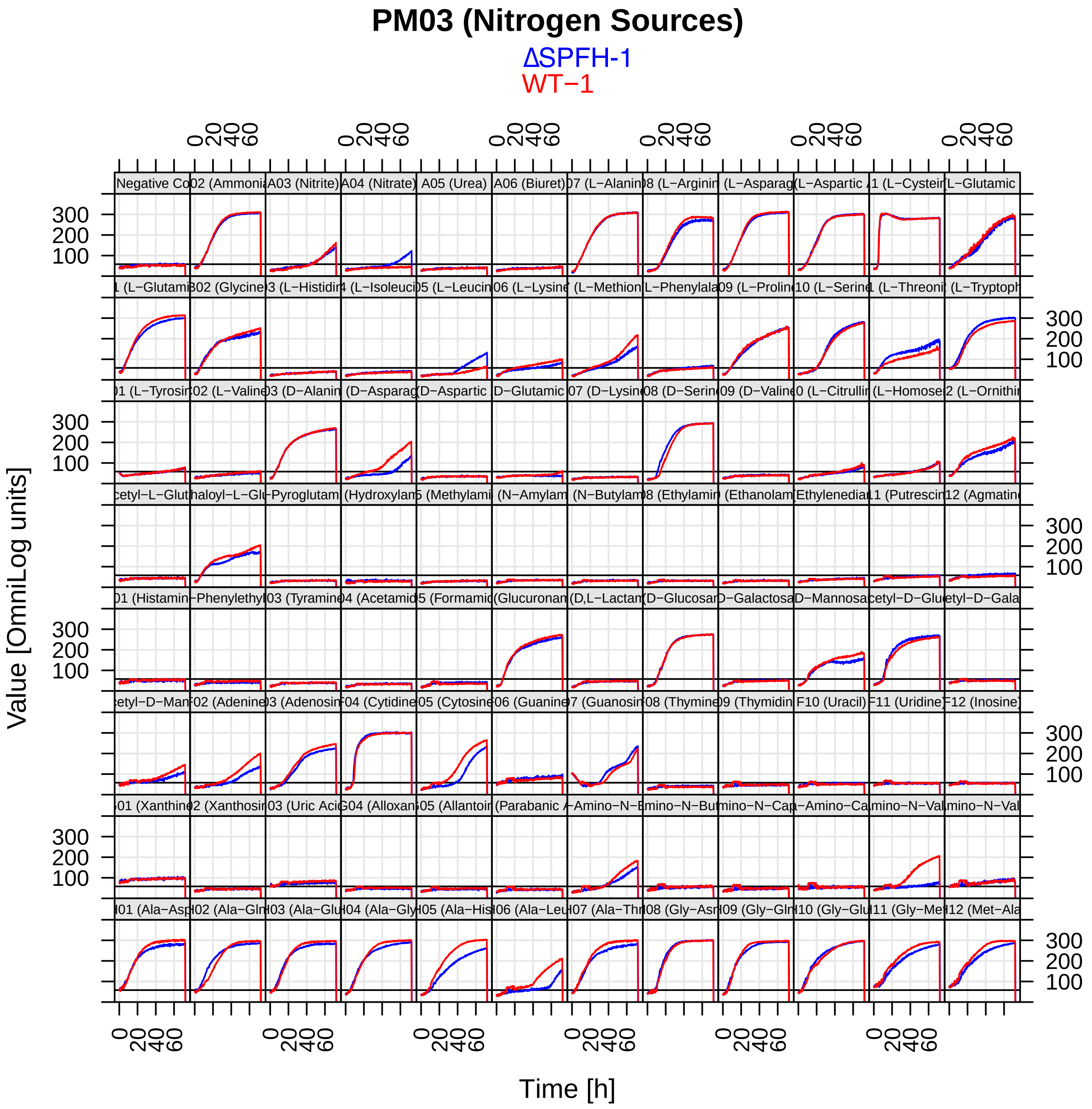
PM03 (Nitrogen Sources)
ΔSPFH-1
WT−1
Value [OmniLog units]
Time [h]

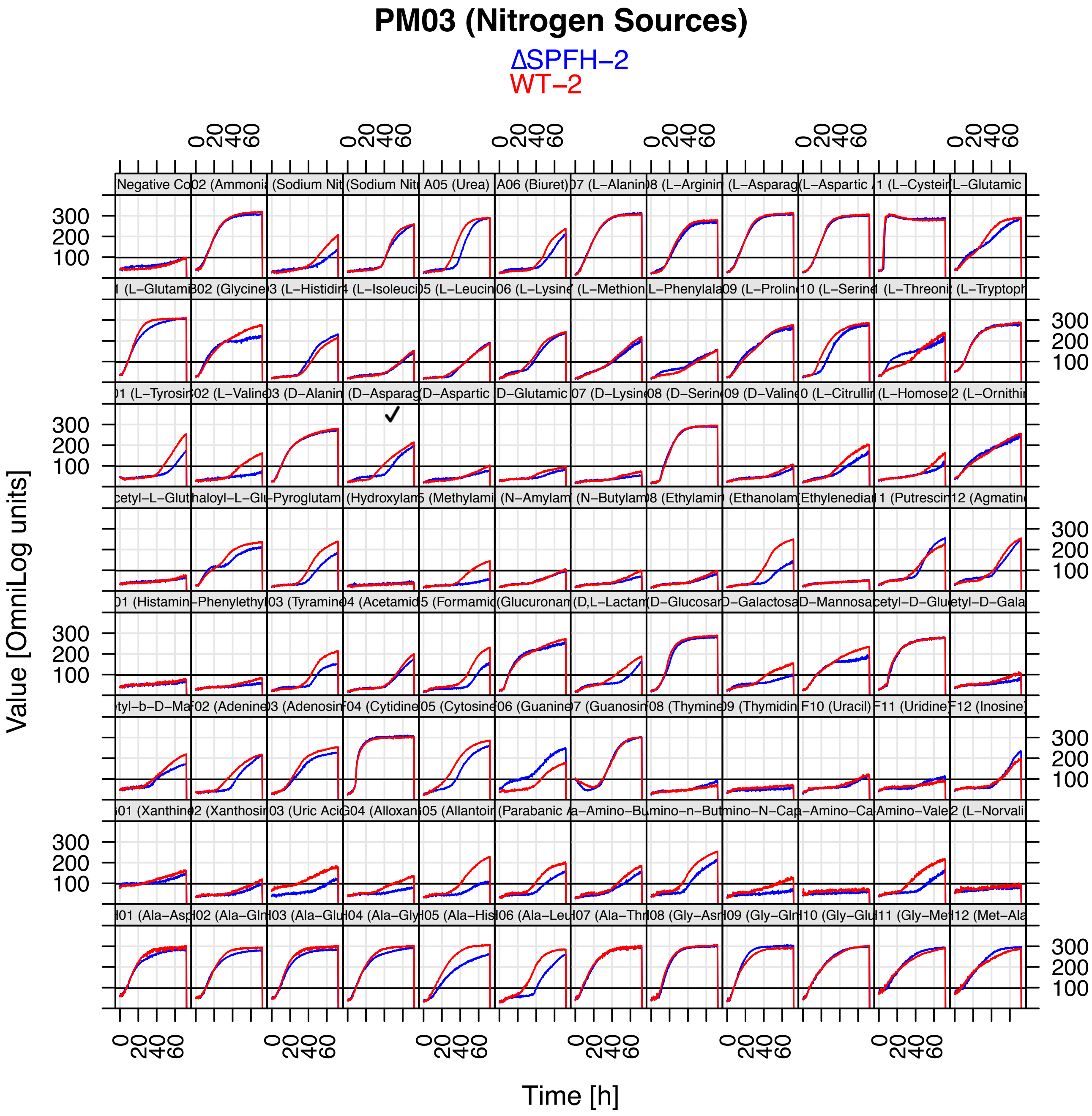
PM03 (Nitrogen Sources)
ΔSPFH−2
WT−2
Value [OmniLog units]
Time [h]

### Phenotype MicroArrays™

#### PM9 MicroPlate™ Osmolytes

| | | | | | | | | | | | |
|---|---|---|---|---|---|---|---|---|---|---|---|
| A1<br>NaCl 1% | A2<br>NaCl 2% | A3<br>NaCl 3% | A4<br>NaCl 4% | A5<br>NaCl 5% | A6<br>NaCl 5.5% | A7<br>NaCl 6% | A8<br>NaCl 6.5% | A9<br>NaCl 7% | A10<br>NaCl 8% | A11<br>NaCl 9% | A12<br>NaCl 10% |
| B1<br>NaCl 6% | B2<br>NaCl 6% +<br>Betaine | B3<br>NaCl 6% +<br>N-N Dimethyl glycine | B4<br>NaCl 6% +<br>Sarcosine | B5<br>NaCl 6% +<br>Dimethyl sulphonyl propionate | B6<br>NaCl 6% +<br>MOPS | B7<br>NaCl 6% +<br>Ectoine | B8<br>NaCl 6% +<br>Choline | B9<br>NaCl 6% +<br>Phosphoryl choline | B10<br>NaCl 6% +<br>Creatine | B11<br>NaCl 6% +<br>Creatinine | B12<br>NaCl 6% +<br>L- Carnitine |
| C1<br>NaCl 6% +<br>KCl | C2<br>NaCl 6% +<br>L-proline | C3<br>NaCl 6% +<br>N-Acethyl L-glutamine | C4<br>NaC1 6% +<br>β-Glutamic acid | C5<br>NaC1 6% +<br>γ–Amino -n-butyric acid | C6<br>NaC1 6% +<br>Glutathione | C7<br>NaCl 6% +<br>Glycerol | C8<br>NaC1 6% +<br>Trehalose | C9<br>NaC1 6% +<br>Trimethylamine -N-oxide | C10<br>NaCl 6% +<br>Trimethylamine | C11<br>NaCl 6% +<br>Octopine | C12<br>NaC1 6% +<br>Trigonelline |
| D-1<br>Potassium chloride 3% | D2<br>Potassium chloride 4% | D3<br>Potassium chloride 5% | D4<br>Potassium chloride 6% | D5<br>Sodium sulfate 2% | D6<br>Sodium sulfate 3% | D7<br>Sodium sulfate 4% | D8<br>Sodium sulfate 5% | D9<br>Ethylene glycol 5% | D10<br>Ethylene glycol 10% | D11<br>Ethylene glycol 15% | D12<br>Ethylene glycol 20% |
| E1<br>Sodium formate 1% | E2<br>Sodium formate 2% | E3<br>Sodium formate 3% | E4<br>Sodium formate 4% | E5<br>Sodium formate 5% | E6<br>Sodium formate 6% | E7<br>Urea 2% | E8<br>Urea 3% | E9<br>Urea 4% | E10<br>Urea 5% | E11<br>Urea 6% | E12<br>Urea 7% |
| F1<br>Sodium Lactate 1% | F2<br>Sodium Lactate 2% | F3<br>Sodium Lactate 3% | F4<br>Sodium Lactate 4% | F5<br>Sodium Lactate 5% | F6<br>Sodium Lactate 6% | F7<br>Sodium Lactate 7% | F8<br>Sodium Lactate 8% | F9<br>Sodium Lactate 9% | F10<br>Sodium Lactate 10% | F11<br>Sodium Lactate 11% | F12<br>Sodium Lactate 12% |
| G1<br>Sodium Phosphate pH 7 20mM | G2<br>Sodium Phosphate pH 7 50mM | G3<br>Sodium Phosphate pH 7 100mM | G4<br>Sodium Phosphate pH 7 200mM | G5<br>Sodium Benzoate pH 5.2 20mM | G6<br>Sodium Benzoate pH 5.2 50mM | G7<br>Sodium Benzoate pH5.2 100mM | G8<br>Sodium Benzoate pH 5.2 200mM | G9<br>Ammonium sulfate pH8 10mM | G10<br>Ammonium sulfate pH 8 20mM | G11<br>Ammonium sulfate pH 8 50mM | G12<br>Ammonium sulfate pH8 100mM |
| H1<br>Sodium Nitrate 10mM | H2<br>Sodium Nitrate 20mM | H3<br>Sodium Nitrate 40mM | H4<br>Sodium Nitrate 60mM | H5<br>Sodium Nitrate 80mM | H6<br>Sodium Nitrate 100mM | H7<br>Sodium Nitrite 10mM | H8<br>Sodium Nitrite 20mM | H9<br>Sodium Nitrite 40mM | H10<br>Sodium Nitrite 60mM | H11<br>Sodium Nitrite 80mM | H12<br>Sodium Nitrite 100mM |

#### PM10 MicroPlate™ pH

| | | | | | | | | | | | |
|---|---|---|---|---|---|---|---|---|---|---|---|
| A1<br>pH 3.5 | A2<br>pH 4 | A3<br>pH 4.5 | A4<br>pH 5 | A5<br>pH 5.5 | A6<br>pH 6 | A7<br>pH 7 | A8<br>pH 8 | A9<br>pH 8.5 | A10<br>pH 9 | A11<br>pH 9.5 | A12<br>pH 10 |
| B1<br>pH 4.5 | B2<br>pH 4.5 +<br>L-Alanine | B3<br>pH 4.5 +<br>L-Arginine | B4<br>pH 4.5 +<br>L-Asparagine | B5<br>pH 4.5 +<br>L-Aspartic Acid | B6<br>pH 4.5 +<br>L-Glutamic Acid | B7<br>pH 4.5 +<br>L-Glutamine | B8<br>pH 4.5 +<br>Glycine | B9<br>pH 4.5 +<br>L-Histidine | B10<br>pH 4.5 +<br>L-Isoleucine | B11<br>pH 4.5 +<br>L-Leucine | B12<br>pH 4.5 +<br>L-Lysine |
| C1<br>pH 4.5 +<br>L-Methionine | C2<br>pH 4.5 +<br>L-Phenylalanine | C3<br>pH 4.5 +<br>L-Proline | C4<br>pH 4.5 +<br>L-Serine | C5<br>pH 4.5 +<br>L-Threonine | C6<br>pH 4.5 +<br>L-Tryptophan | C7<br>pH 4.5 +<br>L-Tyrosine | C8<br>pH 4.5 +<br>L-Valine | C9<br>pH 4.5 +<br>Hydroxy-L-Proline | C10<br>pH 4.5 +<br>L-Ornithine | C11<br>pH 4.5 +<br>L-Homoarginine | C12<br>pH 4.5 +<br>L-Homoserine |
| D-1<br>pH 4.5 +<br>Anthranilic acid | D2<br>pH 4.5 +<br>L-Norleucine | D3<br>pH 4.5 +<br>L-Norvaline | D4<br>pH 4.5 +<br>α- Amino-N-butyric acid | D5<br>pH 4.5 +<br>p-Aminobenzoate | D6<br>pH 4.5 +<br>L-Cysteic acid | D7<br>pH 4.5 +<br>D-Lysine | D8<br>pH 4.5 +<br>5-Hydroxy Lysine | D9<br>pH 4.5 +<br>5-Hydroxy Tryptophan | D10<br>pH 4.5 +<br>D,L-Diamino pimelic acid | D11<br>pH 4.5 +<br>Trimethyl amine-N-oxide | D12<br>pH 4.5 +<br>Urea |
| E1<br>pH 9.5 | E2<br>pH 9.5 +<br>L-Alanine | E3<br>pH 9.5 +<br>L-Arginine | E4<br>pH 9.5 +<br>L-Asparagine | E5<br>pH 9.5 +<br>L-Aspartic Acid | E6<br>pH 9.5 +<br>L-Glutamic Acid | E7<br>pH 9.5 +<br>L-Glutamine | E8<br>pH 9.5 +<br>Glycine | E9<br>pH 9.5 +<br>L-Histidine | E10<br>pH 9.5 +<br>L-Isoleucine | E11<br>pH 9.5 +<br>L-Leucine | E12<br>pH 9.5 +<br>L-Lysine |
| F1<br>pH 9.5 +<br>L-Methionine | F2<br>pH 9.5 +<br>L-Phenylalanine | F3<br>pH 9.5 +<br>L-Proline | F4<br>pH 9.5 +<br>L-Serine | F5<br>pH 9.5 +<br>L-Threonine | F6<br>pH 9.5 +<br>L-Tryptophan | F7<br>pH 9.5 +<br>L-Tyrosine | F8<br>pH 9.5 +<br>L-Valine | F9<br>pH 9.5 +<br>Hydroxy-L-Proline | F10<br>pH 9.5 +<br>L-Ornithine | F11<br>pH 9.5 +<br>L-Homoarginine | F12<br>pH 9.5 +<br>L-Homoserine |
| G1<br>pH 9.5 +<br>Anthranilic acid | G2<br>pH 9.5 +<br>L-Norleucine | G3<br>pH 9.5 +<br>L-Norvaline | G4<br>pH 9.5 +<br>Agmatine | G5<br>pH 9.5 +<br>Cadaverine | G6<br>pH 9.5 +<br>Putrescine | G7<br>pH 9.5 +<br>Histamine | G8<br>pH 9.5 +<br>Phenylethylamine | G9<br>pH 9.5 +<br>Tyramine | G10<br>pH 9.5 +<br>Creatine | G11<br>pH 9.5 +<br>Trimethyl amine-N-oxide | G12<br>pH 9.5 +<br>Urea |
| H1<br>X-Caprylate | H2<br>X–α-D-Glucoside | H3<br>X-β-D-Glucoside | H4<br>X-α-D-Galactoside | H5<br>X-β-D-Galactoside | H6<br>X-α- D-Glucuronide | H7<br>X-β- D-Glucuronide | H8<br>X-β-D-Glucosaminide | H9<br>X-β-D-Galactosaminide | H10<br>X-α-D-Mannoside | H11<br>X-PO4 | H12<br>X-SO4 |

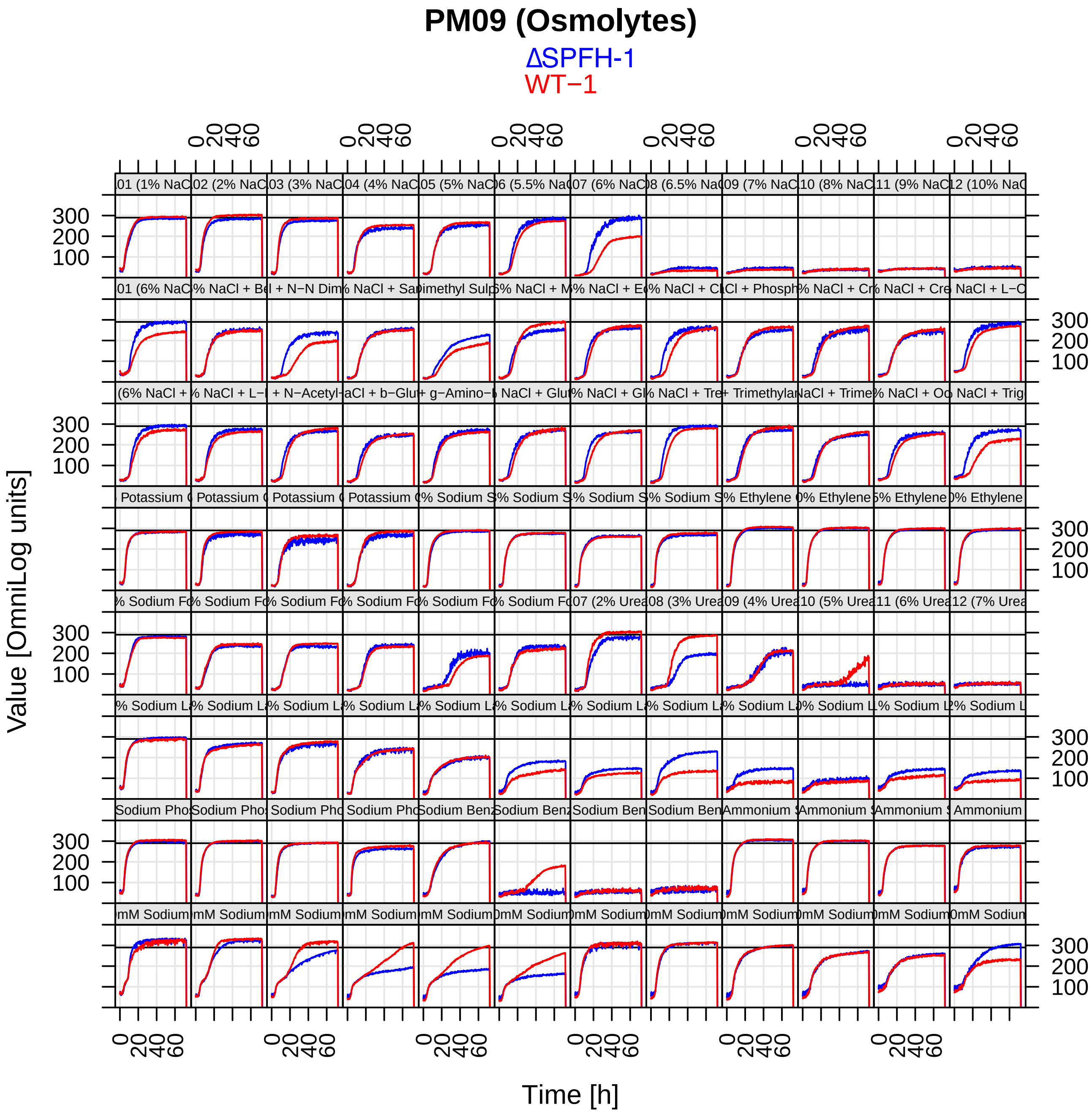
PM09 (Osmolytes)
ΔSPFH-1
WT−1
Value [OmniLog units]
Time [h]

### PM09 (Osmolytes)

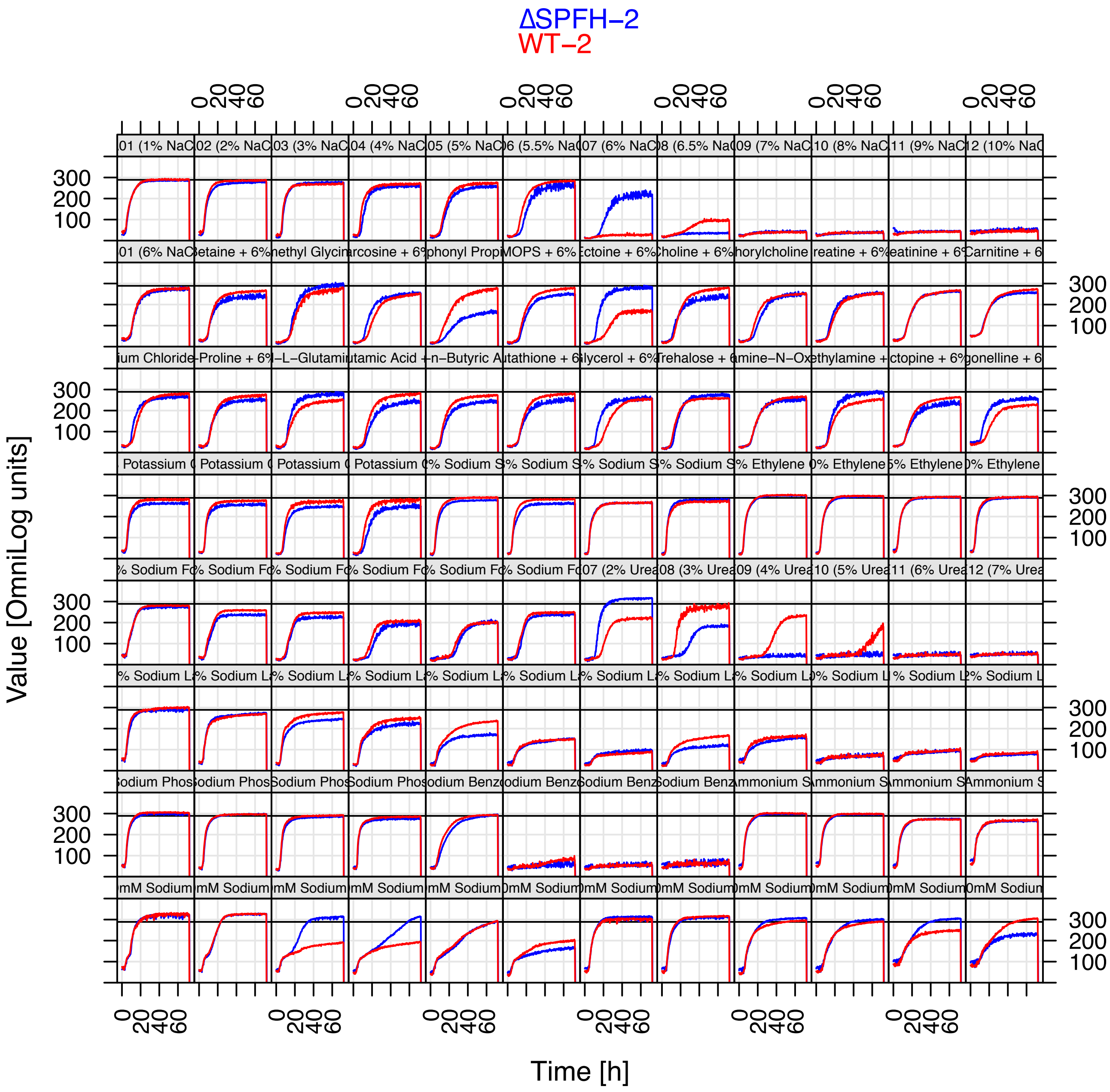

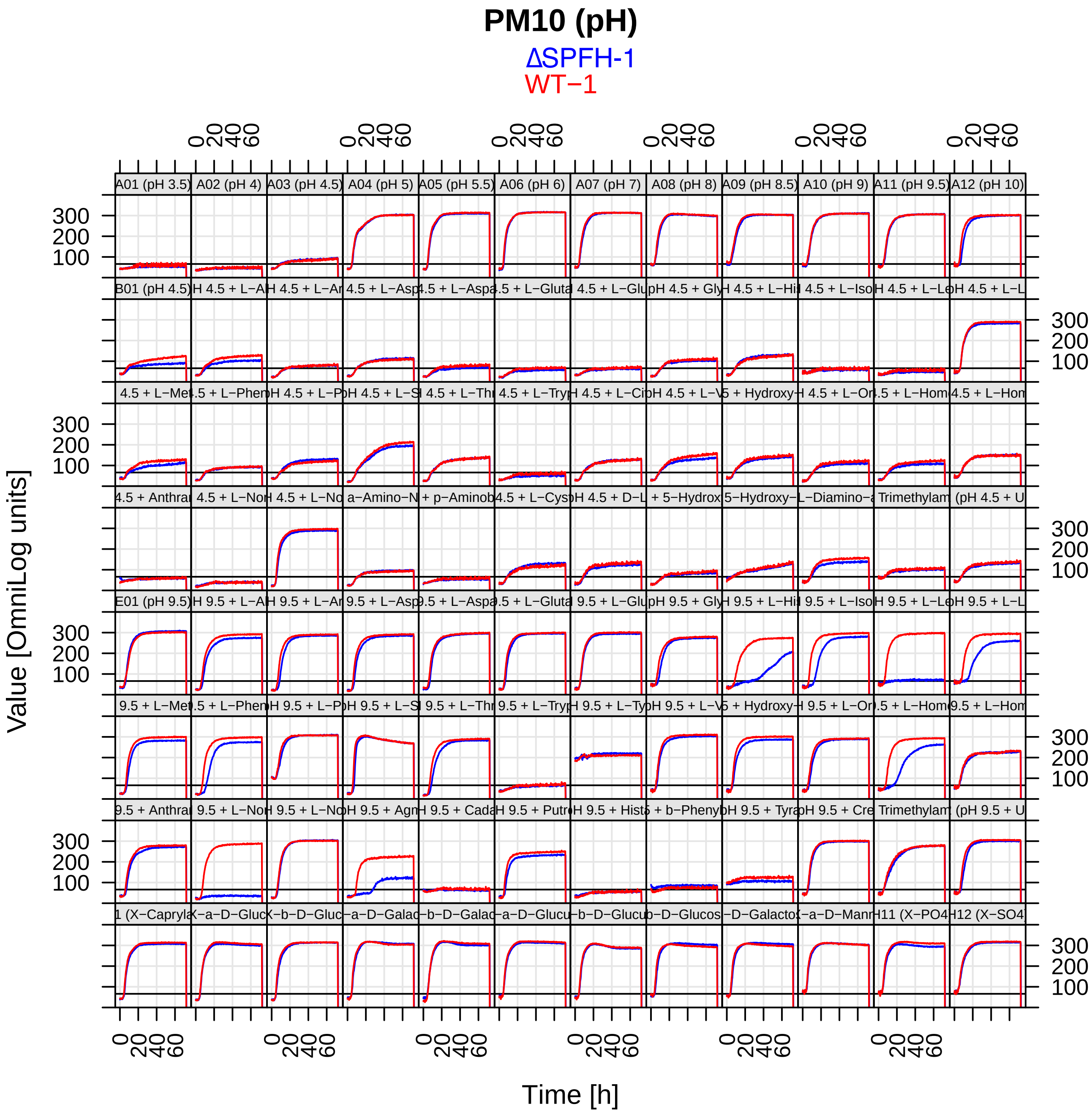
PM10 (pH)
ΔSPFH-1
WT−1
Value [OmniLog units]
Time [h]

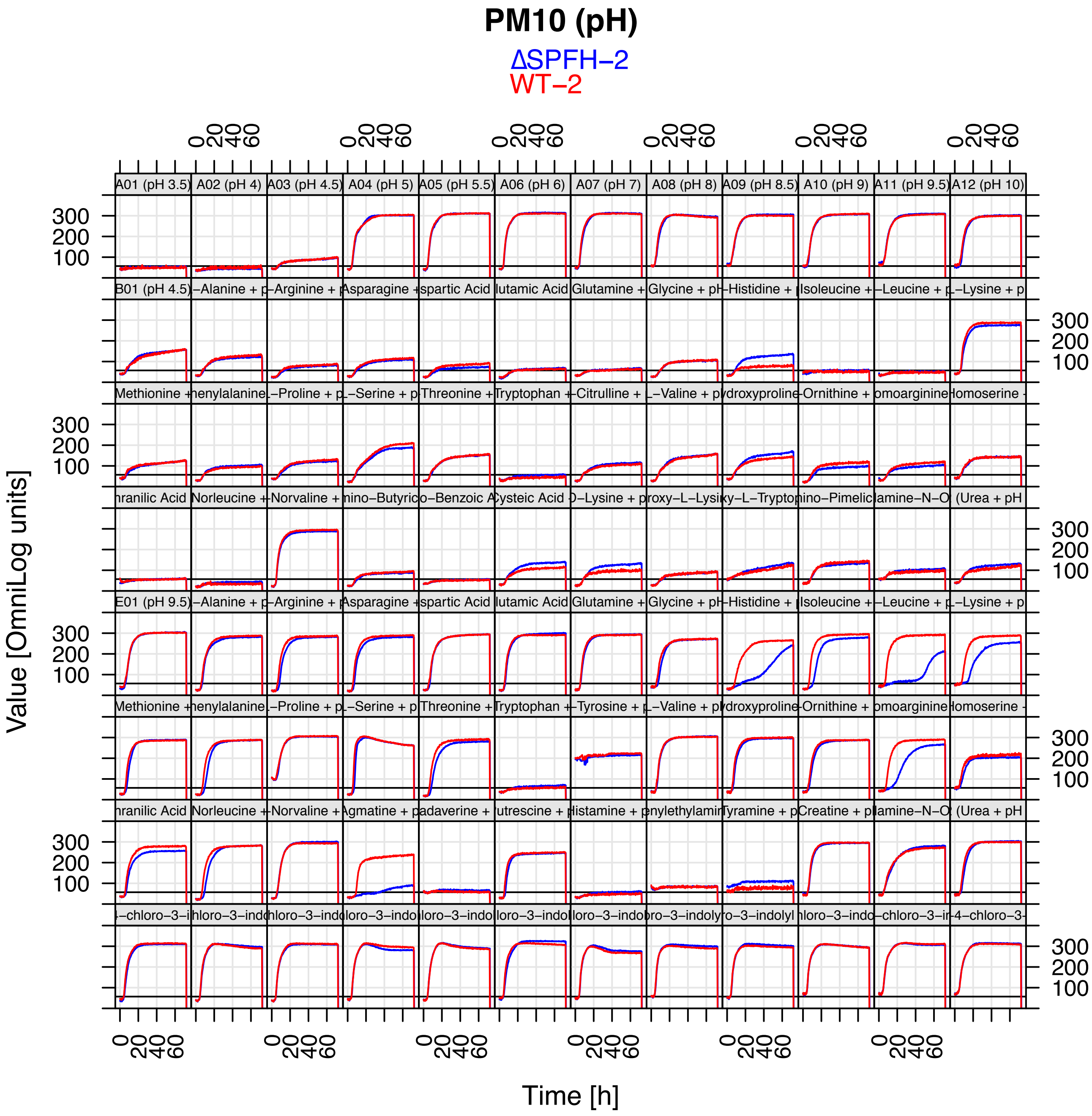
PM10 (pH)
ΔSPFH−2
WT−2
Value [OmniLog units]
Time [h]
A01 (pH 3.5)
A02 (pH 4)
A03 (pH 4.5)
A04 (pH 5)
A05 (pH 5.5)
A06 (pH 6)
A07 (pH 7)
A08 (pH 8)
A09 (pH 8.5)
A10 (pH 9)
A11 (pH 9.5)
A12 (pH 10)
100
200
300
0
20
40
60

#### PM11C MicroPlate™

| | | | | | | | | | | | |
|---|---|---|---|---|---|---|---|---|---|---|---|
| A1<br>Amikacin<br>1 | A2<br>Amikacin<br>2 | A3<br>Amikacin<br>3 | A4<br>Amikacin<br>4 | A5<br>Chlortetracycline<br>1 | A6<br>Chlortetracycline<br>2 | A7<br>Chlortetracycline<br>3 | A8<br>Chlortetracycline<br>4 | A9<br>Lincomycin<br>1 | A10<br>Lincomycin<br>2 | A11<br>Lincomycin<br>3 | A12<br>Lincomycin<br>4 |
| B1<br>Amoxicillin<br>1 | B2<br>Amoxicillin<br>2 | B3<br>Amoxicillin<br>3 | B4<br>Amoxicillin<br>4 | B5<br>Cloxacillin<br>1 | B6<br>Cloxacillin<br>2 | B7<br>Cloxacillin<br>3 | B8<br>Cloxacillin<br>4 | B9<br>Lomefloxacin<br>1 | B10<br>Lomefloxacin<br>2 | B11<br>Lomefloxacin<br>3 | B12<br>Lomefloxacin<br>4 |
| C1<br>Bleomycin<br>1 | C2<br>Bleomycin<br>2 | C3<br>Bleomycin<br>3 | C4<br>Bleomycin<br>4 | C5<br>Colistin<br>1 | C6<br>Colistin<br>2 | C7<br>Colistin<br>3 | C8<br>Colistin<br>4 | C9<br>Minocycline<br>1 | C10<br>Minocycline<br>2 | C11<br>Minocycline<br>3 | C12<br>Minocycline<br>4 |
| D1<br>Capreomycin<br>1 | D2<br>Capreomycin<br>2 | D3<br>Capreomycin<br>3 | D4<br>Capreomycin<br>4 | D5<br>Demeclocycline<br>1 | D6<br>Demeclocycline<br>2 | D7<br>Demeclocycline<br>3 | D8<br>Demeclocycline<br>4 | D9<br>Nafcillin<br>1 | D10<br>Nafcillin<br>2 | D11<br>Nafcillin<br>3 | D12<br>Nafcillin<br>4 |
| E1<br>Cefazolin<br>1 | E2<br>Cefazolin<br>2 | E3<br>Cefazolin<br>3 | E4<br>Cefazolin<br>4 | E5<br>Enoxacin<br>1 | E6<br>Enoxacin<br>2 | E7<br>Enoxacin<br>3 | E8<br>Enoxacin<br>4 | E9<br>Nalidixic acid<br>1 | E10<br>Nalidixic acid<br>2 | E11<br>Nalidixic acid<br>3 | E12<br>Nalidixic acid<br>4 |
| F1<br>Chloramphenicol<br>1 | F2<br>Chloramphenicol<br>2 | F3<br>Chloramphenicol<br>3 | F4<br>Chloramphenicol<br>4 | F5<br>Erythromycin<br>1 | F6<br>Erythromycin<br>2 | F7<br>Erythromycin<br>3 | F8<br>Erythromycin<br>4 | F9<br>Neomycin<br>1 | F10<br>Neomycin<br>2 | F11<br>Neomycin<br>3 | F12<br>Neomycin<br>4 |
| G1<br>Ceftriaxone<br>1 | G2<br>Ceftriaxone<br>2 | G3<br>Ceftriaxone<br>3 | G4<br>Ceftriaxone<br>4 | G5<br>Gentamicin<br>1 | G6<br>Gentamicin<br>2 | G7<br>Gentamicin<br>3 | G8<br>Gentamicin<br>4 | G9<br>Potassium tellurite<br>1 | G10<br>Potassium tellurite<br>2 | G11<br>Potassium tellurite<br>3 | G12<br>Potassium tellurite<br>4 |
| H1<br>Cephalothin<br>1 | H2<br>Cephalothin<br>2 | H3<br>Cephalothin<br>3 | H4<br>Cephalothin<br>4 | H5<br>Kanamycin<br>1 | H6<br>Kanamycin<br>2 | H7<br>Kanamycin<br>3 | H8<br>Kanamycin<br>4 | H9<br>Ofloxacin<br>1 | H10<br>Ofloxacin<br>2 | H11<br>Ofloxacin<br>3 | H12<br>Ofloxacin<br>4 |

#### PM12B MicroPlate™

| | | | | | | | | | | | |
|---|---|---|---|---|---|---|---|---|---|---|---|
| A1<br>Penicillin G<br>1 | A2<br>Penicillin G<br>2 | A3<br>Penicillin G<br>3 | A4<br>Penicillin G<br>4 | A5<br>Tetracycline<br>1 | A6<br>Tetracycline<br>2 | A7<br>Tetracycline<br>3 | A8<br>Tetracycline<br>4 | A9<br>Carbenicillin<br>1 | A10<br>Carbenicillin<br>2 | A11<br>Carbenicillin<br>3 | A12<br>Carbenicillin<br>4 |
| B1<br>Oxacillin<br>1 | B2<br>Oxacillin<br>2 | B3<br>Oxacillin<br>3 | B4<br>Oxacillin<br>4 | B5<br>Penimepicycline<br>1 | B6<br>Penimepicycline<br>2 | B7<br>Penimepicycline<br>3 | B8<br>Penimepicycline<br>4 | B9<br>Polymyxin B<br>1 | B10<br>Polymyxin B<br>2 | B11<br>Polymyxin B<br>3 | B12<br>Polymyxin B<br>4 |
| C1<br>Paromomycin<br>1 | C2<br>Paromomycin<br>2 | C3<br>Paromomycin<br>3 | C4<br>Paromomycin<br>4 | C5<br>Vancomycin<br>1 | C6<br>Vancomycin<br>2 | C7<br>Vancomycin<br>3 | C8<br>Vancomycin<br>4 | C9<br>D,L-Serine hydroxamate<br>1 | C10<br>D,L-Serine hydroxamate<br>2 | C11<br>D,L-Serine hydroxamate<br>3 | C12<br>D,L-Serine hydroxamate<br>4 |
| D1<br>Sisomicin<br>1 | D2<br>Sisomicin<br>2 | D3<br>Sisomicin<br>3 | D4<br>Sisomicin<br>4 | D5<br>Sulfamethazine<br>1 | D6<br>Sulfamethazine<br>2 | D7<br>Sulfamethazine<br>3 | D8<br>Sulfamethazine<br>4 | D9<br>Novobiocin<br>1 | D10<br>Novobiocin<br>2 | D11<br>Novobiocin<br>3 | D12<br>Novobiocin<br>4 |
| E1<br>2,4-Diamino-6,7-diisopropyl-pteridine<br>1 | E2<br>2,4-Diamino-6,7-diisopropyl-pteridine<br>2 | E3<br>2,4-Diamino-6,7-diisopropyl-pteridine<br>3 | E4<br>2,4-Diamino-6,7-diisopropyl-pteridine<br>4 | E5<br>Sulfadiazine<br>1 | E6<br>Sulfadiazine<br>2 | E7<br>Sulfadiazine<br>3 | E8<br>Sulfadiazine<br>4 | E9<br>Benzethonium chloride<br>1 | E10<br>Benzethonium chloride<br>2 | E11<br>Benzethonium chloride<br>3 | E12<br>Benzethonium chloride<br>4 |
| F1<br>Tobramycin<br>1 | F2<br>Tobramycin<br>2 | F3<br>Tobramycin<br>3 | F4<br>Tobramycin<br>4 | F5<br>Sulfathiazole<br>1 | F6<br>Sulfathiazole<br>2 | F7<br>Sulfathiazole<br>3 | F8<br>Sulfathiazole<br>4 | F9<br>5-Fluoroorotic acid<br>1 | F10<br>5-Fluoroorotic acid<br>2 | F11<br>5-Fluoroorotic acid<br>3 | F12<br>5-Fluoroorotic acid<br>4 |
| G1<br>Spectinomycin<br>1 | G2<br>Spectinomycin<br>2 | G3<br>Spectinomycin<br>3 | G4<br>Spectinomycin<br>4 | G5<br>Sulfa-methoxazole<br>1 | G6<br>Sulfa-methoxazole<br>2 | G7<br>Sulfa-methoxazole<br>3 | G8<br>Sulfa-methoxazole<br>4 | G9<br>L-Aspartic-β-hydroxamate<br>1 | G10<br>L-Aspartic-β-hydroxamate<br>2 | G11<br>L-Aspartic-β-hydroxamate<br>3 | G12<br>L-Aspartic-β-hydroxamate<br>4 |
| H1<br>Spiramycin<br>1 | H2<br>Spiramycin<br>2 | H3<br>Spiramycin<br>3 | H4<br>Spiramycin<br>4 | H5<br>Rifampicin<br>1 | H6<br>Rifampicin<br>2 | H7<br>Rifampicin<br>3 | H8<br>Rifampicin<br>4 | H9<br>Dodecyltrimethyl ammonium bromide<br>1 | H10<br>Dodecyltrimethyl ammonium bromide<br>2 | H11<br>Dodecyltrimethyl ammonium bromide<br>3 | H12<br>Dodecyltrimethyl ammonium bromide<br>4 |

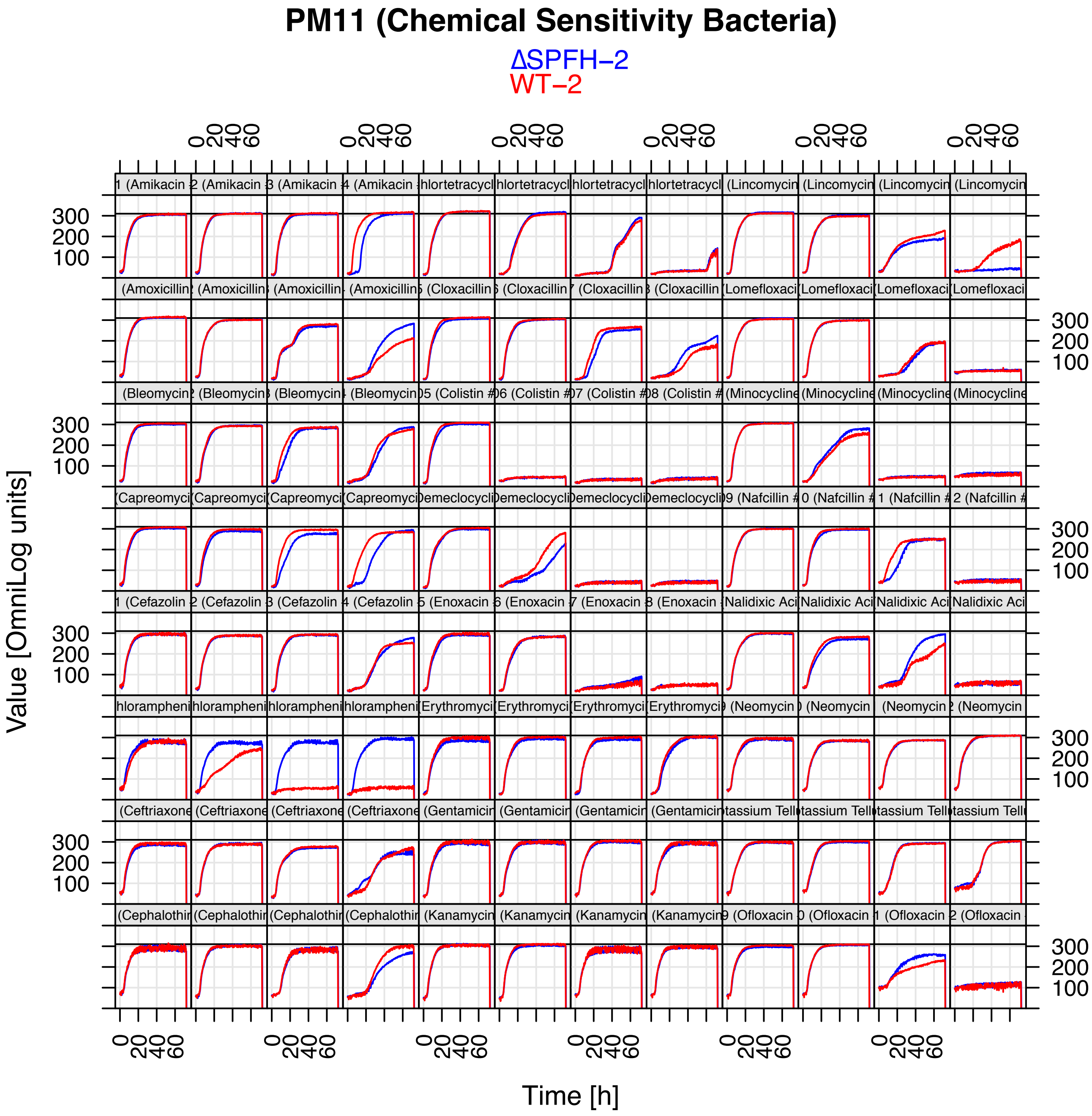
PM11 (Chemical Sensitivity Bacteria)
ΔSPFH−2
WT−2
Value [OmniLog units]
Time [h]

### PM11 (Chemicals)

ΔSPFH-1
WT−1

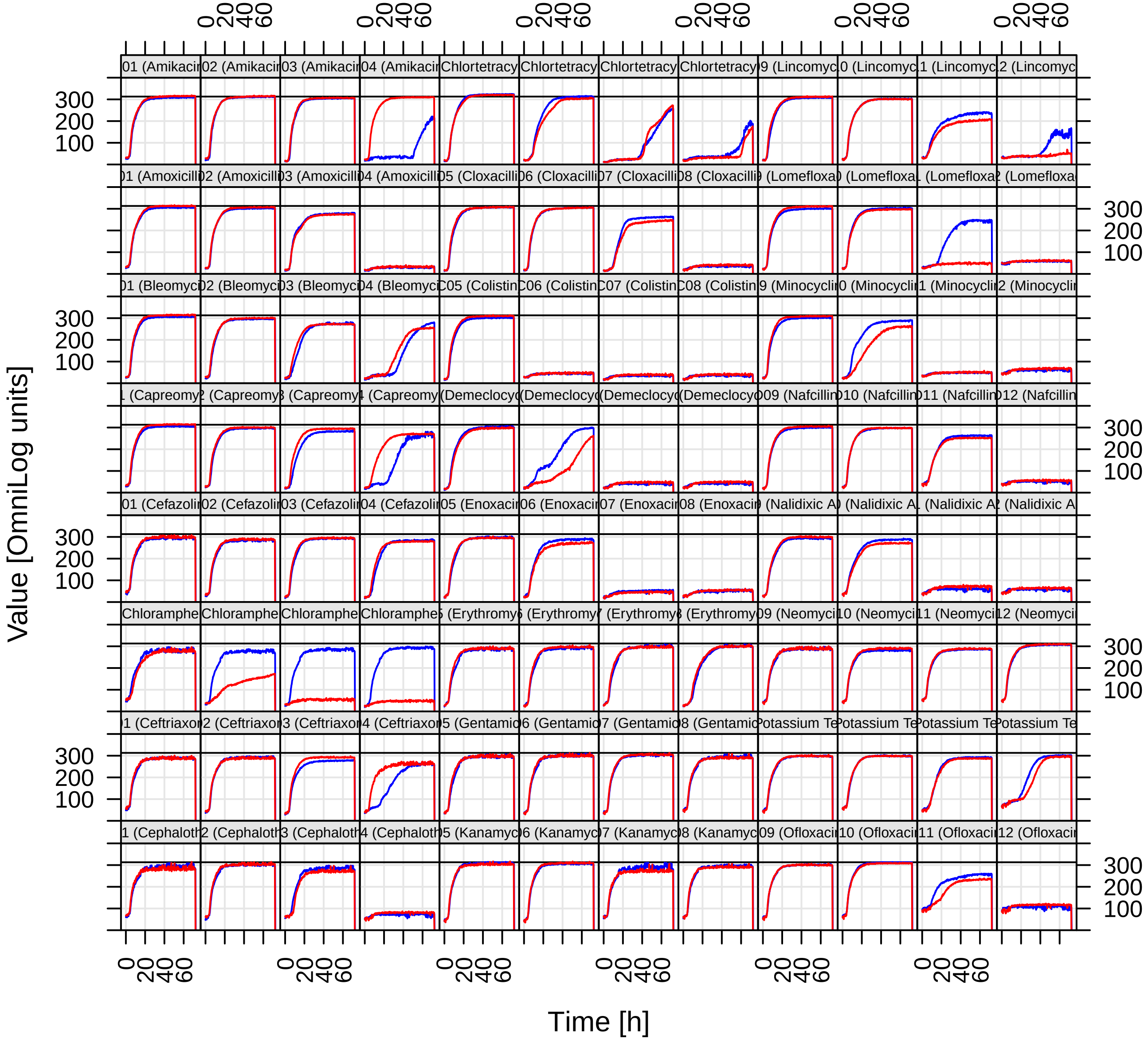

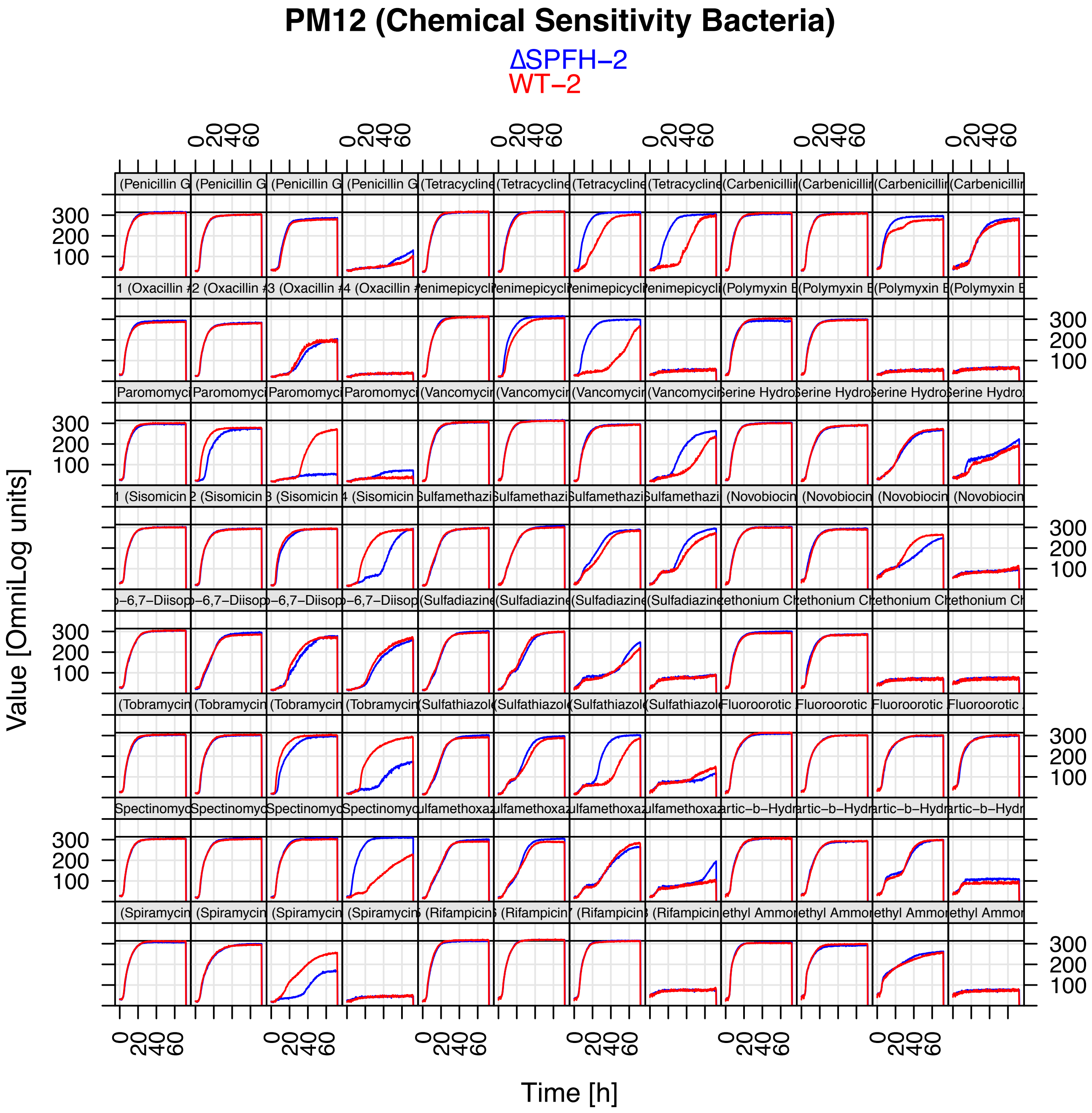
PM12 (Chemical Sensitivity Bacteria)
ΔSPFH−2
WT−2
Value [OmniLog units]
Time [h]

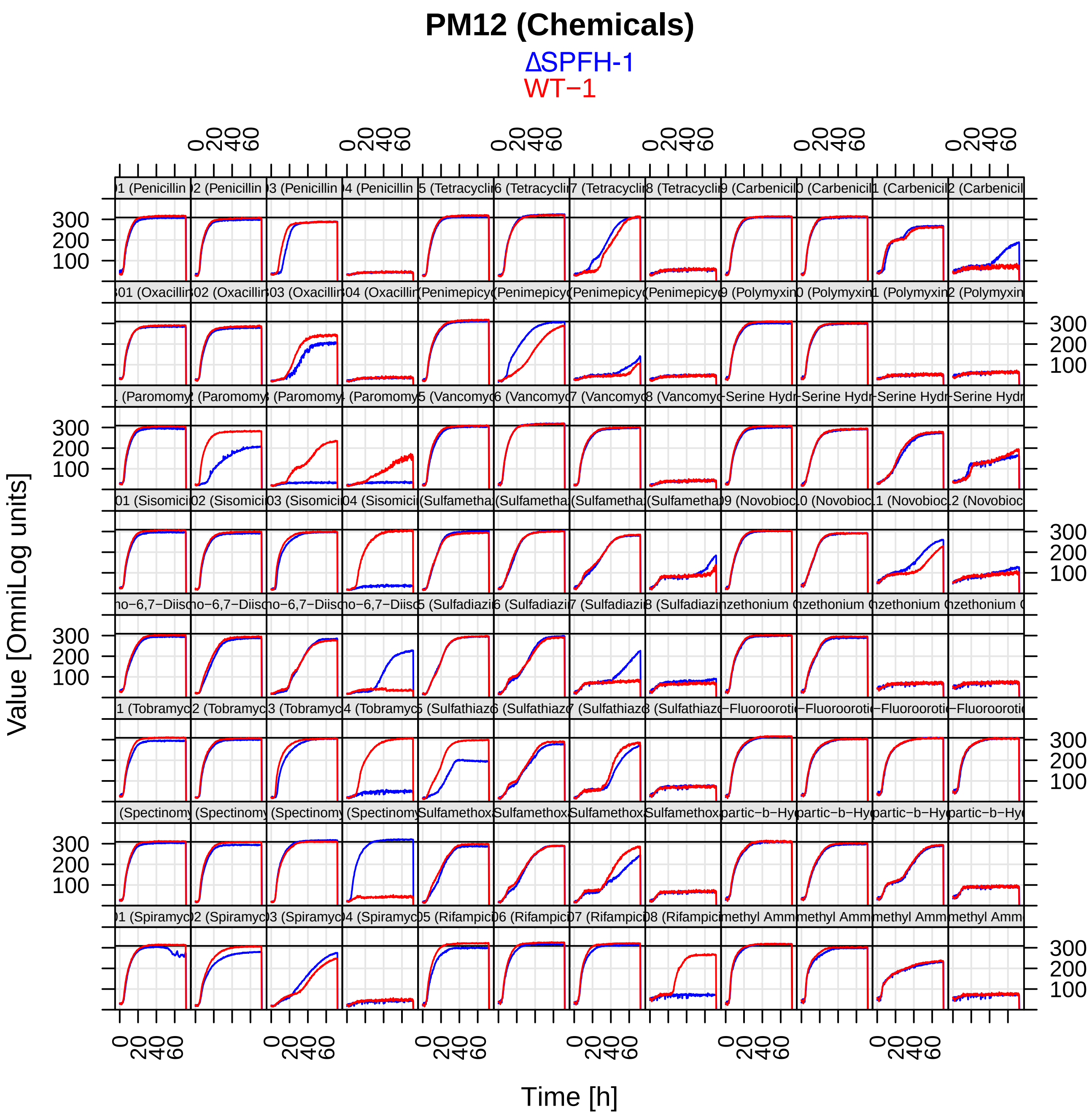
PM12 (Chemicals)
ΔSPFH-1
WT−1
Value [OmniLog units]
Time [h]
0
20
40
60
100
200
300
)1 (Penicillin
)2 (Penicillin
)3 (Penicillin
)4 (Penicillin
)5 (Tetracyclin
)6 (Tetracyclin
)7 (Tetracyclin
)8 (Tetracyclin
)9 (Carbenicil
)0 (Carbenicil
)1 (Carbenicil
)2 (Carbenicil
)01 (Oxacillin
)02 (Oxacillin
)03 (Oxacillin
)04 (Oxacillin
(Penimepicy
(Penimepicy
(Penimepicy
(Penimepicy
)9 (Polymyxin
)0 (Polymyxin
)1 (Polymyxin
)2 (Polymyxin
. (Paromomy
(Paromomy
(Paromomy
(Paromomy
)5 (Vancomyc
)6 (Vancomyc
)7 (Vancomyc
)8 (Vancomyc
-Serine Hydr
-Serine Hydr
-Serine Hydr
-Serine Hydr
)01 (Sisomici
)02 (Sisomici
)03 (Sisomici
)04 (Sisomici
(Sulfametha
(Sulfametha
(Sulfametha
(Sulfametha
)9 (Novobioc
)0 (Novobioc
)1 (Novobioc
)2 (Novobioc
no−6,7−Diis
no−6,7−Diis
no−6,7−Diis
no−6,7−Diis
)5 (Sulfadiazi
)6 (Sulfadiazi
)7 (Sulfadiazi
)8 (Sulfadiazi
nzethonium
nzethonium
nzethonium
nzethonium
)1 (Tobramyc
)2 (Tobramyc
)3 (Tobramyc
)4 (Tobramyc
)5 (Sulfathiaz
)6 (Sulfathiaz
)7 (Sulfathiaz
)8 (Sulfathiaz
−Fluoroorotic
−Fluoroorotic
−Fluoroorotic
−Fluoroorotic
(Spectinomy
(Spectinomy
(Spectinomy
(Spectinomy
Sulfamethox
Sulfamethox
Sulfamethox
Sulfamethox
partic−b−Hy
partic−b−Hy
partic−b−Hy
partic−b−Hy
)1 (Spiramyc
)2 (Spiramyc
)3 (Spiramyc
)4 (Spiramyc
)5 (Rifampic
)6 (Rifampic
)7 (Rifampic
)8 (Rifampic
methyl Amm
methyl Amm
methyl Amm
methyl Amm

#### PM13B MicroPlate™

| | | | | | | | | | | | |
|---|---|---|---|---|---|---|---|---|---|---|---|
| A1 Ampicillin 1 | A2 Ampicillin 2 | A3 Ampicillin 3 | A4 Ampicillin 4 | A5 Dequalinium chloride 1 | A6 Dequalinium chloride 2 | A7 Dequalinium chloride 3 | A8 Dequalinium chloride 4 | A9 Nickel chloride 1 | A10 Nickel chloride 2 | A11 Nickel chloride 3 | A12 Nickel chloride 4 |
| B1 Azlocillin 1 | B2 Azlocillin 2 | B3 Azlocillin 3 | B4 Azlocillin 4 | B5 2, 2'-Dipyridyl 1 | B6 2, 2'-Dipyridyl 2 | B7 2, 2'-Dipyridyl 3 | B8 2, 2'-Dipyridyl 4 | B9 Oxolinic acid 1 | B10 Oxolinic acid 2 | B11 Oxolinic acid 3 | B12 Oxolinic acid 4 |
| C1 6-Mercapto-purine 1 | C2 6-Mercapto-purine 2 | C3 6-Mercapto-purine 3 | C4 6-Mercapto-purine 4 | C5 Doxycycline 1 | C6 Doxycycline 2 | C7 Doxycycline 3 | C8 Doxycycline 4 | C9 Potassium chromate 1 | C10 Potassium chromate 2 | C11 Potassium chromate 3 | C12 Potassium chromate 4 |
| D1 Cefuroxime 1 | D2 Cefuroxime 2 | D3 Cefuroxime 3 | D4 Cefuroxime 4 | D5 5-Fluorouracil 1 | D6 5-Fluorouracil 2 | D7 5-Fluorouracil 3 | D8 5-Fluorouracil 4 | D9 Rolitetracycline 1 | D10 Rolitetracycline 2 | D11 Rolitetracycline 3 | D12 Rolitetracycline 4 |
| E1 Cytosine-1-beta-D-arabino-furanoside 1 | E2 Cytosine-1-beta-D-arabino-furanoside 2 | E3 Cytosine-1-beta-D-arabino-furanoside 3 | E4 Cytosine-1-beta-D-arabino-furanoside 4 | E5 Geneticin (G418) 1 | E6 Geneticin (G418) 2 | E7 Geneticin (G418) 3 | E8 Geneticin (G418) 4 | E9 Ruthenium red 1 | E10 Ruthenium red 2 | E11 Ruthenium red 3 | E12 Ruthenium red 4 |
| F1 Cesium chloride 1 | F2 Cesium chloride 2 | F3 Cesium chloride 3 | F4 Cesium chloride 4 | F5 Glycine 1 | F6 Glycine 2 | F7 Glycine 3 | F8 Glycine 4 | F9 Thallium (I) acetate 1 | F10 Thallium (I) acetate 2 | F11 Thallium (I) acetate 3 | F12 Thallium (I) acetate 4 |
| G1 Cobalt chloride 1 | G2 Cobalt chloride 2 | G3 Cobalt chloride 3 | G4 Cobalt chloride 4 | G5 Manganese chloride 1 | G6 Manganese chloride 2 | G7 Manganese chloride 3 | G8 Manganese chloride 4 | G9 Trifluoperazine 1 | G10 Trifluoperazine 2 | G11 Trifluoperazine 3 | G12 Trifluoperazine 4 |
| H1 Cupric chloride 1 | H2 Cupric chloride 2 | H3 Cupric chloride 3 | H4 Cupric chloride 4 | H5 Moxalactam 1 | H6 Moxalactam 2 | H7 Moxalactam 3 | H8 Moxalactam 4 | H9 Tylosin 1 | H10 Tylosin 2 | H11 Tylosin 3 | H12 Tylosin 4 |

#### PM14A MicroPlate™

| | | | | | | | | | | | |
|---|---|---|---|---|---|---|---|---|---|---|---|
| A1 Acriflavine 1 | A2 Acriflavine 2 | A3 Acriflavine 3 | A4 Acriflavine 4 | A5 Furaltadone 1 | A6 Furaltadone 2 | A7 Furaltadone 3 | A8 Furaltadone 4 | A9 Sanguinarine 1 | A10 Sanguinarine 2 | A11 Sanguinarine 3 | A12 Sanguinarine 4 |
| B1 9-Aminoacridine 1 | B2 9-Aminoacridine 2 | B3 9-Aminoacridine 3 | B4 9-Aminoacridine 4 | B5 Fusaric acid 1 | B6 Fusaric acid 2 | B7 Fusaric acid 3 | B8 Fusaric acid 4 | B9 Sodium arsenate 1 | B10 Sodium arsenate 2 | B11 Sodium arsenate 3 | B12 Sodium arsenate 4 |
| C1 Boric Acid 1 | C2 Boric Acid 2 | C3 Boric Acid 3 | C4 Boric Acid 4 | C5 1-Hydroxy-pyridine -2-thione 1 | C6 1-Hydroxy-pyridine -2-thione 2 | C7 1-Hydroxy-pyridine -2-thione 3 | C8 1-Hydroxy-pyridine -2-thione 4 | C9 Sodium cyanate 1 | C10 Sodium cyanate 2 | C11 Sodium cyanate 3 | C12 Sodium cyanate 4 |
| D1 Cadmium chloride 1 | D2 Cadmium chloride 2 | D3 Cadmium chloride 3 | D4 Cadmium chloride 4 | D5 Iodoacetate 1 | D6 Iodoacetate 2 | D7 Iodoacetate 3 | D8 Iodoacetate 4 | D9 Sodium dichromate 1 | D10 Sodium dichromate 2 | D11 Sodium dichromate 3 | D12 Sodium dichromate 4 |
| E1 Cefoxitin 1 | E2 Cefoxitin 2 | E3 Cefoxitin 3 | E4 Cefoxitin 4 | E5 Nitrofurantoin 1 | E6 Nitrofurantoin 2 | E7 Nitrofurantoin 3 | E8 Nitrofurantoin 4 | E9 Sodium metaborate 1 | E10 Sodium metaborate 2 | E11 Sodium metaborate 3 | E12 Sodium metaborate 4 |
| F1 Chloramphenicol 1 | F2 Chloramphenicol 2 | F3 Chloramphenicol 3 | F4 Chloramphenicol 4 | F5 Piperacillin 1 | F6 Piperacillin 2 | F7 Piperacillin 3 | F8 Piperacillin 4 | F9 Sodium metavanadate 1 | F10 Sodium metavanadate 2 | F11 Sodium metavanadate 3 | F12 Sodium metavanadate 4 |
| G1 Chelerythrine 1 | G2 Chelerythrine 2 | G3 Chelerythrine 3 | G4 Chelerythrine 4 | G5 Carbenicillin 1 | G6 Carbenicillin 2 | G7 Carbenicillin 3 | G8 Carbenicillin 4 | G9 Sodium nitrite 1 | G10 Sodium nitrite 2 | G11 Sodium nitrite 3 | G12 Sodium nitrite 4 |
| H1 EGTA 1 | H2 EGTA 2 | H3 EGTA 3 | H4 EGTA 4 | H5 Promethazine 1 | H6 Promethazine 2 | H7 Promethazine 3 | H8 Promethazine 4 | H9 Sodium orthovanadate 1 | H10 Sodium orthovanadate 2 | H11 Sodium orthovanadate 3 | H12 Sodium orthovanadate 4 |

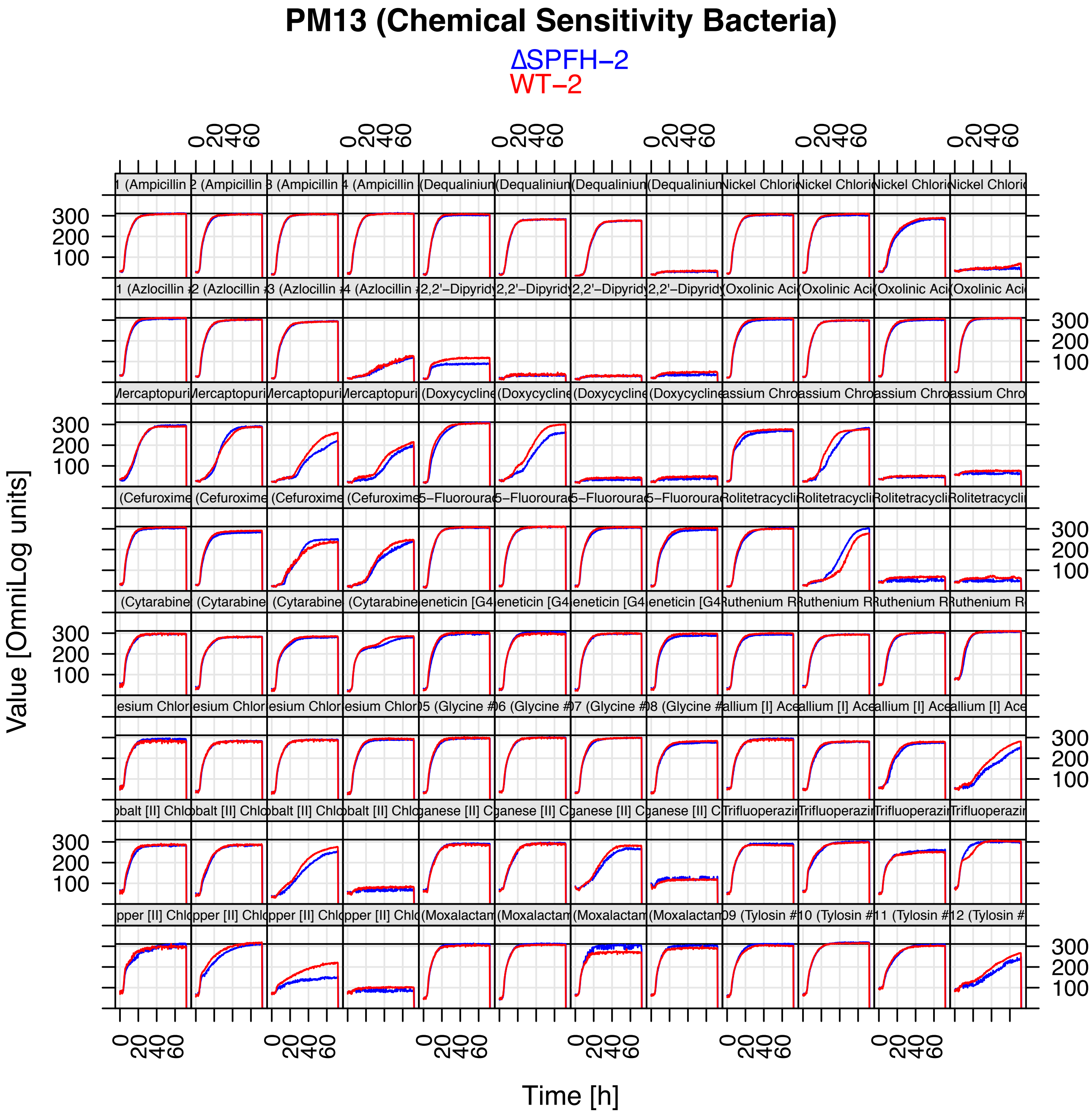
PM13 (Chemical Sensitivity Bacteria)
ΔSPFH−2
WT−2
1 (Ampicillin
2 (Ampicillin
3 (Ampicillin
4 (Ampicillin
(Dequaliniun
(Dequaliniun
(Dequaliniun
(Dequaliniun
Nickel Chlori
Nickel Chlori
Nickel Chlori
Nickel Chlori
1 (Azlocillin
2 (Azlocillin
3 (Azlocillin
4 (Azlocillin
2,2'−Dipyrid
2,2'−Dipyrid
2,2'−Dipyrid
2,2'−Dipyrid
Oxolinic Aci
Oxolinic Aci
Oxolinic Aci
Oxolinic Aci
Mercaptopuri
Mercaptopuri
Mercaptopuri
Mercaptopuri
(Doxycycline
(Doxycycline
(Doxycycline
(Doxycycline
assium Chro
assium Chro
assium Chro
assium Chro
(Cefuroxime
(Cefuroxime
(Cefuroxime
(Cefuroxime
5−Fluoroura
5−Fluoroura
5−Fluoroura
5−Fluoroura
Rolitetracycli
Rolitetracycli
Rolitetracycli
Rolitetracycli
(Cytarabine
(Cytarabine
(Cytarabine
(Cytarabine
eneticin [G4
eneticin [G4
eneticin [G4
eneticin [G4
Ruthenium R
Ruthenium R
Ruthenium R
Ruthenium R
esium Chlor
esium Chlor
esium Chlor
esium Chlor
05 (Glycine #
06 (Glycine #
07 (Glycine #
08 (Glycine #
allium [I] Ace
allium [I] Ace
allium [I] Ace
allium [I] Ace
obalt [II] Chl
obalt [II] Chl
obalt [II] Chl
obalt [II] Chl
ganese [II] C
ganese [II] C
ganese [II] C
ganese [II] C
Trifluoperazi
Trifluoperazi
Trifluoperazi
Trifluoperazi
pper [II] Chl
pper [II] Chl
pper [II] Chl
pper [II] Chl
(Moxalactam
(Moxalactam
(Moxalactam
(Moxalactam
09 (Tylosin #
10 (Tylosin #
11 (Tylosin #
12 (Tylosin #
100
200
300
0
20
40
60
Value [OmniLog units]
Time [h]

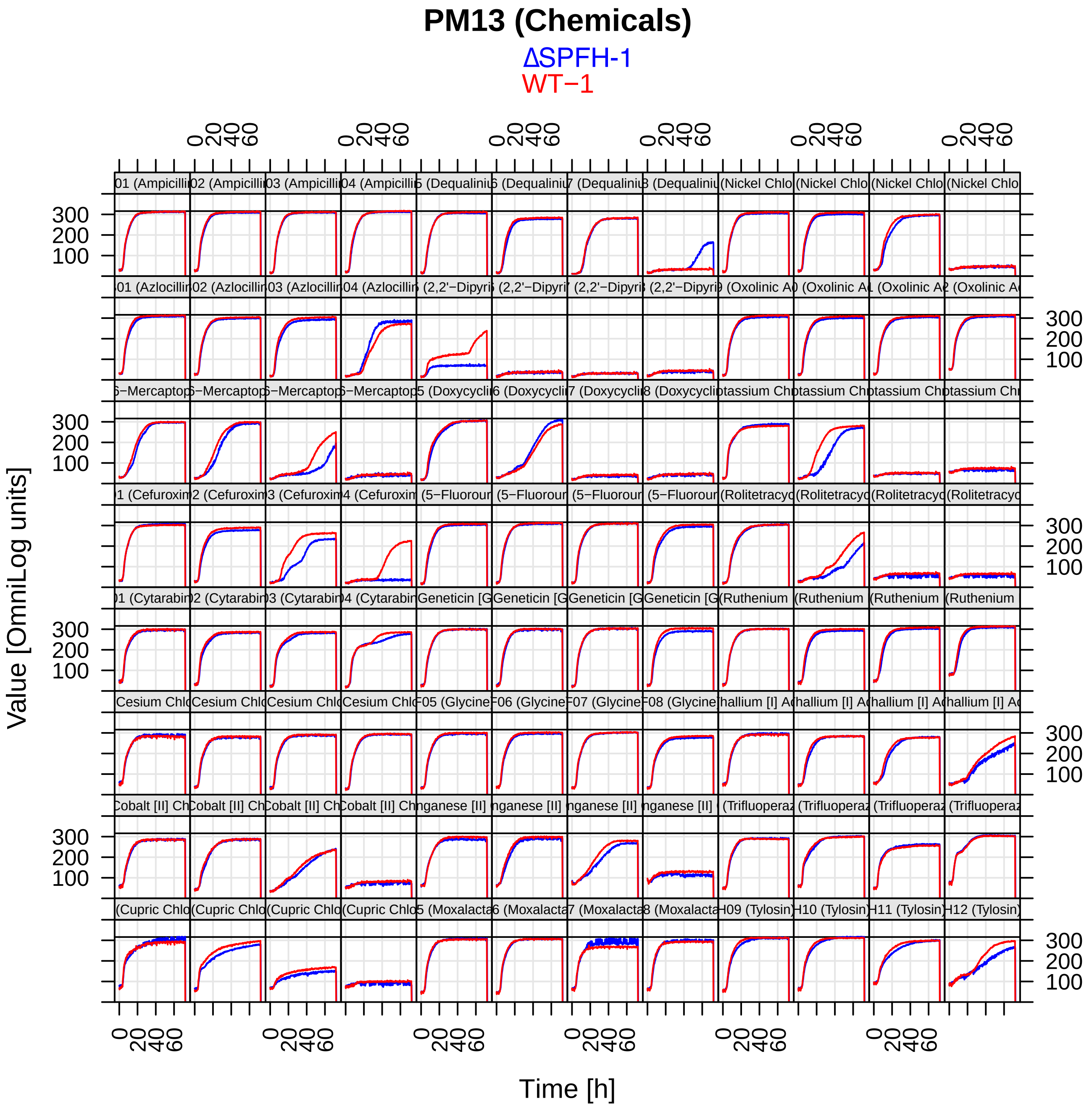
PM13 (Chemicals)
ΔSPFH-1
WT−1
Value [OmniLog units]
Time [h]

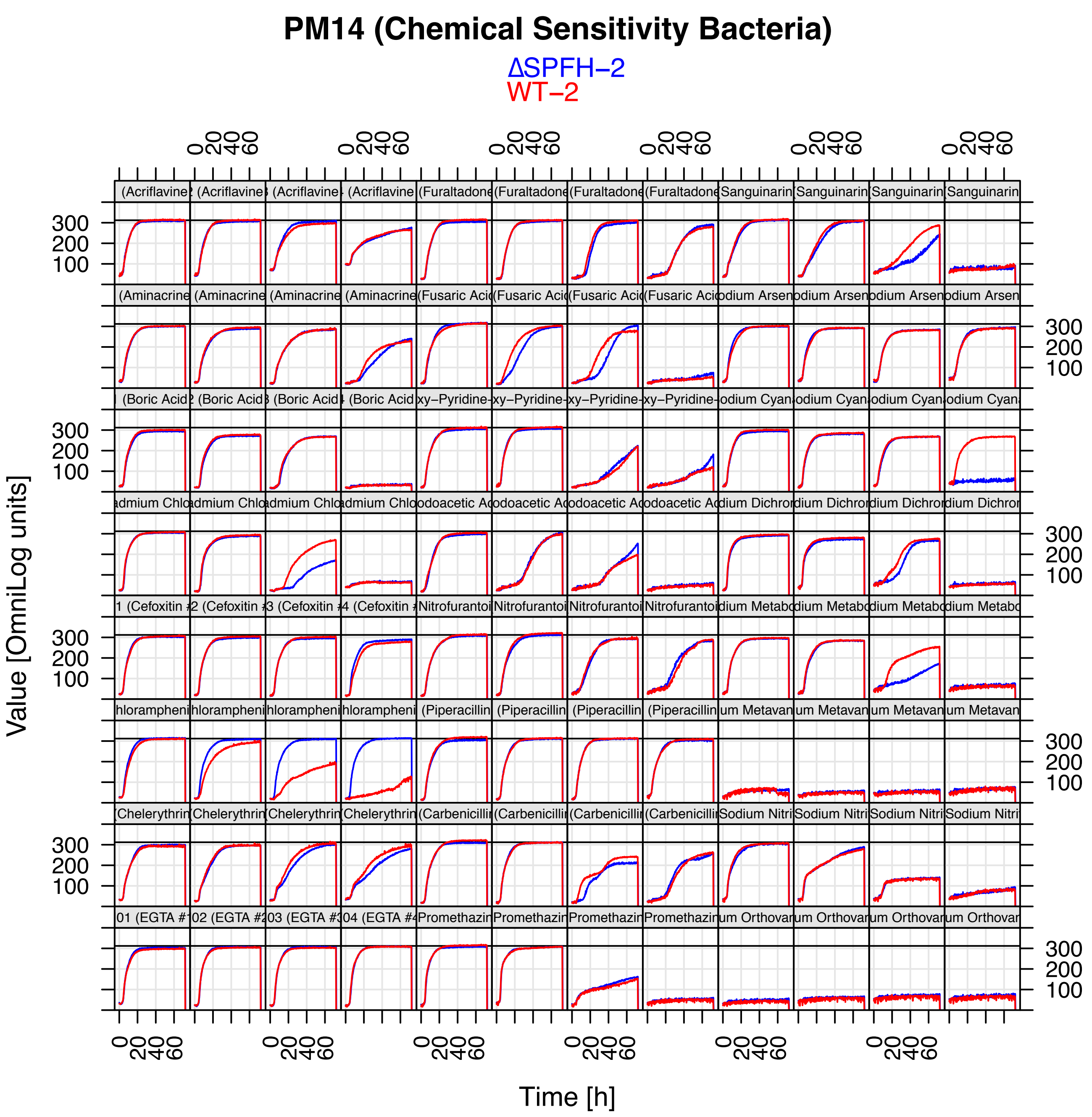
PM14 (Chemical Sensitivity Bacteria)
ΔSPFH−2
WT−2
Value [OmniLog units]
Time [h]
(Acriflavine
(Aminacrine
(Boric Acid
Cadmium Chlo
Cefoxitin
Chloramphenicol
Chelerythrine
(EGTA
(Furaltadone
(Fusaric Acid
Pyridine
Iodoacetic Acid
Nitrofurantoin
(Piperacillin
(Carbenicillin
Promethazine
Sanguinarine
Sodium Arsenite
Sodium Cyanide
Sodium Dichromate
Sodium Metaborate
Sodium Metavanadate
Sodium Nitrite
Sodium Orthovanadate

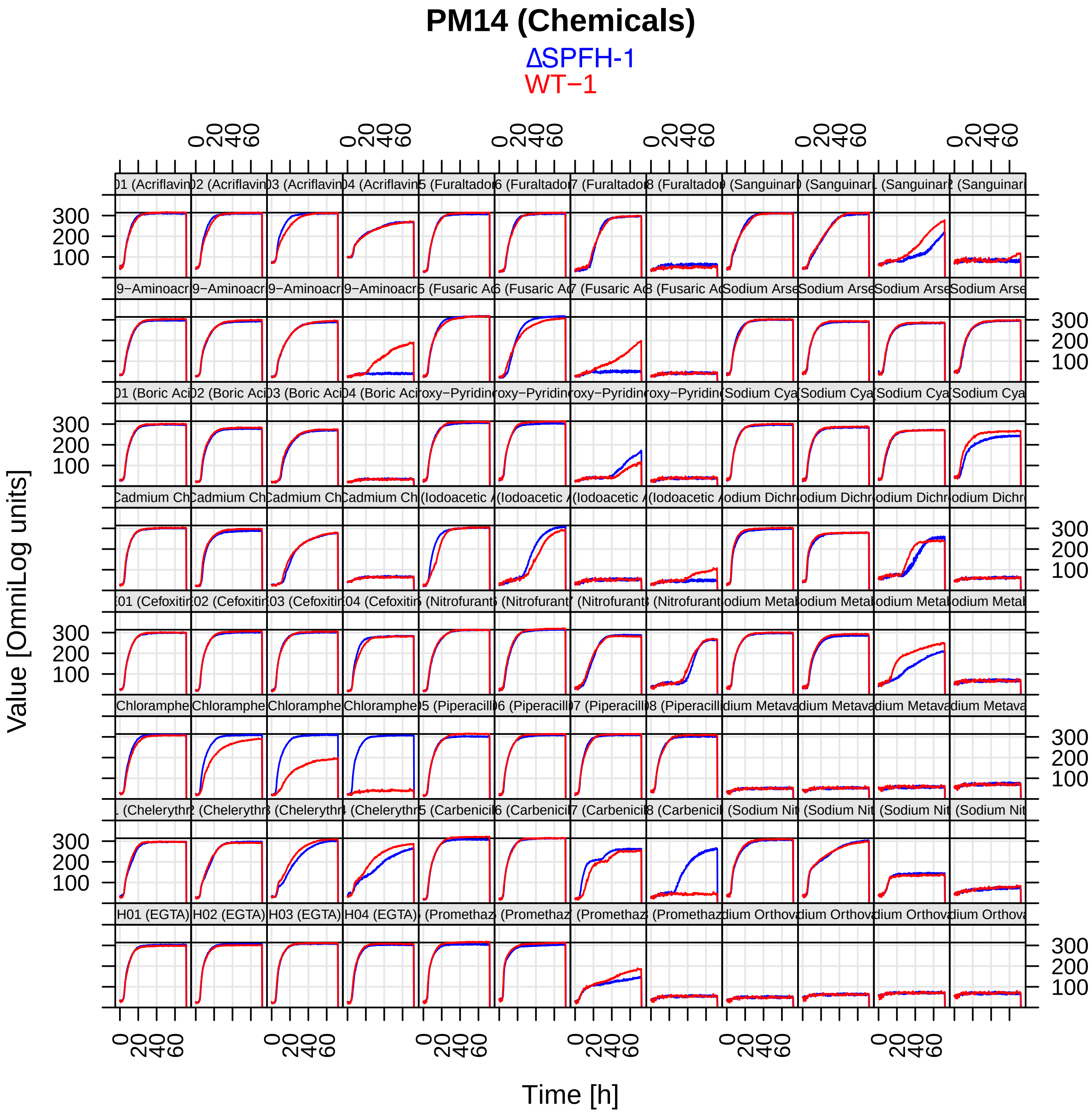

PM14 (Chemicals)
ΔSPFH-1
WT−1
Value [OmniLog units]
Time [h]

#### PM15B MicroPlate™

| | | | | | | | | | | | |
|---|---|---|---|---|---|---|---|---|---|---|---|
| A1<br>Procaine<br>1 | A2<br>Procaine<br>2 | A3<br>Procaine<br>3 | A4<br>Procaine<br>4 | A5<br>Guanidine hydrochloride<br>1 | A6<br>Guanidine hydrochloride<br>2 | A7<br>Guanidine hydrochloride<br>3 | A8<br>Guanidine hydrochloride<br>4 | A9<br>Cefmetazole<br>1 | A10<br>Cefmetazole<br>2 | A11<br>Cefmetazole<br>3 | A12<br>Cefmetazole<br>4 |
| B1<br>D-Cycloserine<br>1 | B2<br>D-Cycloserine<br>2 | B3<br>D-Cycloserine<br>3 | B4<br>D-Cycloserine<br>4 | B5<br>EDTA<br>1 | B6<br>EDTA<br>2 | B7<br>EDTA<br>3 | B8<br>EDTA<br>4 | B9<br>5,7-Dichloro- 8-hydroxy-quinaldine<br>1 | B10<br>5,7-Dichloro- 8-hydroxy-quinaldine<br>2 | B11<br>5,7-Dichloro- 8-hydroxy-quinaldine<br>3 | B12<br>5,7-Dichloro- 8-hydroxy-quinaldine<br>4 |
| C1<br>5,7-Dichloro-8-hydroxyquinoline<br>1 | C2<br>5,7-Dichloro-8-hydroxyquinoline<br>2 | C3<br>5,7-Dichloro-8-hydroxyquinoline<br>3 | C4<br>5,7-Dichloro-8-hydroxyquinoline<br>4 | C5<br>Fusidic acid<br>1 | C6<br>Fusidic acid<br>2 | C7<br>Fusidic acid<br>3 | C8<br>Fusidic acid<br>4 | C9<br>1,10-Phenanthroline<br>1 | C10<br>1,10-Phenanthroline<br>2 | C11<br>1,10-Phenanthroline<br>3 | C12<br>1,10-Phenanthroline<br>4 |
| D1<br>Phleomycin<br>1 | D2<br>Phleomycin<br>2 | D3<br>Phleomycin<br>3 | D4<br>Phleomycin<br>4 | D5<br>Domiphen bromide<br>1 | D6<br>Domiphen bromide<br>2 | D7<br>Domiphen bromide<br>3 | D8<br>Domiphen bromide<br>4 | D9<br>Nordihydroguaiaretic acid<br>1 | D10<br>Nordihydroguaiaretic acid<br>2 | D11<br>Nordihydroguaiaretic acid<br>3 | D12<br>Nordihydroguaiaretic acid<br>4 |
| E1<br>Alexidine<br>1 | E2<br>Alexidine<br>2 | E3<br>Alexidine<br>3 | E4<br>Alexidine<br>4 | E5<br>5-Nitro-2-furaldehyde semicarbazone<br>1 | E6<br>5-Nitro-2-furaldehyde semicarbazone<br>2 | E7<br>5-Nitro-2-furaldehyde semicarbazone<br>3 | E8<br>5-Nitro-2-furaldehyde semicarbazone<br>4 | E9<br>Methyl viologen<br>1 | E10<br>Methyl viologen<br>2 | E11<br>Methyl viologen<br>3 | E12<br>Methyl viologen<br>4 |
| F1<br>3, 4-Dimethoxy-benzyl alcohol<br>1 | F2<br>3, 4-Dimethoxy-benzyl alcohol<br>2 | F3<br>3, 4-Dimethoxy-benzyl alcohol<br>3 | F4<br>3, 4-Dimethoxy-benzyl alcohol<br>4 | F5<br>Oleandomycin<br>1 | F6<br>Oleandomycin<br>2 | F7<br>Oleandomycin<br>3 | F8<br>Oleandomycin<br>4 | F9<br>Puromycin<br>1 | F10<br>Puromycin<br>2 | F11<br>Puromycin<br>3 | F12<br>Puromycin<br>4 |
| G1<br>CCCP<br>1 | G2<br>CCCP<br>2 | G3<br>CCCP<br>3 | G4<br>CCCP<br>4 | G5<br>Sodium azide<br>1 | G6<br>Sodium azide<br>2 | G7<br>Sodium azide<br>3 | G8<br>Sodium azide<br>4 | G9<br>Menadione<br>1 | G10<br>Menadione<br>2 | G11<br>Menadione<br>3 | G12<br>Menadione<br>4 |
| H1<br>2-Nitroimidazole<br>1 | H2<br>2-Nitroimidazole<br>2 | H3<br>2-Nitroimidazole<br>3 | H4<br>2-Nitroimidazole<br>4 | H5<br>Hydroxyurea<br>1 | H6<br>Hydroxyurea<br>2 | H7<br>Hydroxyurea<br>3 | H8<br>Hydroxyurea<br>4 | H9<br>Zinc chloride<br>1 | H10<br>Zinc chloride<br>2 | H11<br>Zinc chloride<br>3 | H12<br>Zinc chloride<br>4 |

#### PM16A MicroPlate™

| | | | | | | | | | | | |
|---|---|---|---|---|---|---|---|---|---|---|---|
| A1<br>Cefotaxime<br>1 | A2<br>Cefotaxime<br>2 | A3<br>Cefotaxime<br>3 | A4<br>Cefotaxime<br>4 | A5<br>Phosphomycin<br>1 | A6<br>Phosphomycin<br>2 | A7<br>Phosphomycin<br>3 | A8<br>Phosphomycin<br>4 | A9<br>5-Chloro-7-iodo-8-hydroxy-quinoline<br>1 | A10<br>5-Chloro-7-iodo-8-hydroxy-quinoline<br>2 | A11<br>5-Chloro-7-iodo-8-hydroxy-quinoline<br>3 | A12<br>5-Chloro-7-iodo-8-hydroxy-quinoline<br>4 |
| B1<br>Norfloxacin<br>1 | B2<br>Norfloxacin<br>2 | B3<br>Norfloxacin<br>3 | B4<br>Norfloxacin<br>4 | B5<br>Sulfanilamide<br>1 | B6<br>Sulfanilamide<br>2 | B7<br>Sulfanilamide<br>3 | B8<br>Sulfanilamide<br>4 | B9<br>Trimethoprim<br>1 | B10<br>Trimethoprim<br>2 | B11<br>Trimethoprim<br>3 | B12<br>Trimethoprim<br>4 |
| C1<br>Dichlofluanid<br>1 | C2<br>Dichlofluanid<br>2 | C3<br>Dichlofluanid<br>3 | C4<br>Dichlofluanid<br>4 | C5<br>Protamine sulfate<br>1 | C6<br>Protamine sulfate<br>2 | C7<br>Protamine sulfate<br>3 | C8<br>Protamine sulfate<br>4 | C9<br>Cetylpyridinium chloride<br>1 | C10<br>Cetylpyridinium chloride<br>2 | C11<br>Cetylpyridinium chloride<br>3 | C12<br>Cetylpyridinium chloride<br>4 |
| D1<br>1-Chloro -2,4-dinitrobenzene<br>1 | D2<br>1-Chloro -2,4-dinitrobenzene<br>2 | D3<br>1-Chloro -2,4-dinitrobenzene<br>3 | D4<br>1-Chloro -2,4-dinitrobenzene<br>4 | D5<br>Diamide<br>1 | D6<br>Diamide<br>2 | D7<br>Diamide<br>3 | D8<br>Diamide<br>4 | D9<br>Cinoxacin<br>1 | D10<br>Cinoxacin<br>2 | D11<br>Cinoxacin<br>3 | D12<br>Cinoxacin<br>4 |
| E1<br>Streptomycin<br>1 | E2<br>Streptomycin<br>2 | E3<br>Streptomycin<br>3 | E4<br>Streptomycin<br>4 | E5<br>5-Azacytidine<br>1 | E6<br>5-Azacytidine<br>2 | E7<br>5-Azacytidine<br>3 | E8<br>5-Azacytidine<br>4 | E9<br>Rifamycin SV<br>1 | E10<br>Rifamycin SV<br>2 | E11<br>Rifamycin SV<br>3 | E12<br>Rifamycin SV<br>4 |
| F1<br>Potassium tellurite<br>1 | F2<br>Potassium tellurite<br>2 | F3<br>Potassium tellurite<br>3 | F4<br>Potassium tellurite<br>4 | F5<br>Sodium selenite<br>1 | F6<br>Sodium selenite<br>2 | F7<br>Sodium selenite<br>3 | F8<br>Sodium selenite<br>4 | F9<br>Aluminum sulfate<br>1 | F10<br>Aluminum sulfate<br>2 | F11<br>Aluminum sulfate<br>3 | F12<br>Aluminum sulfate<br>4 |
| G1<br>Chromium chloride<br>1 | G2<br>Chromium chloride<br>2 | G3<br>Chromium chloride<br>3 | G4<br>Chromium chloride<br>4 | G5<br>Ferric chloride<br>1 | G6<br>Ferric chloride<br>2 | G7<br>Ferric chloride<br>3 | G8<br>Ferric chloride<br>4 | G9<br>L-Glutamic-g-hydroxamate<br>1 | G10<br>L-Glutamic-g-hydroxamate<br>2 | G11<br>L-Glutamic-g-hydroxamate<br>3 | G12<br>L-Glutamic-g-hydroxamate<br>4 |
| H1<br>Glycine hydroxamate<br>1 | H2<br>Glycine hydroxamate<br>2 | H3<br>Glycine hydroxamate<br>3 | H4<br>Glycine hydroxamate<br>4 | H5<br>Chloroxylenol<br>1 | H6<br>Chloroxylenol<br>2 | H7<br>Chloroxylenol<br>3 | H8<br>Chloroxylenol<br>4 | H9<br>Sorbic acid<br>1 | H10<br>Sorbic acid<br>2 | H11<br>Sorbic acid<br>3 | H12<br>Sorbic acid<br>4 |

### PM15 (Chemical Sensitivity Bacteria)

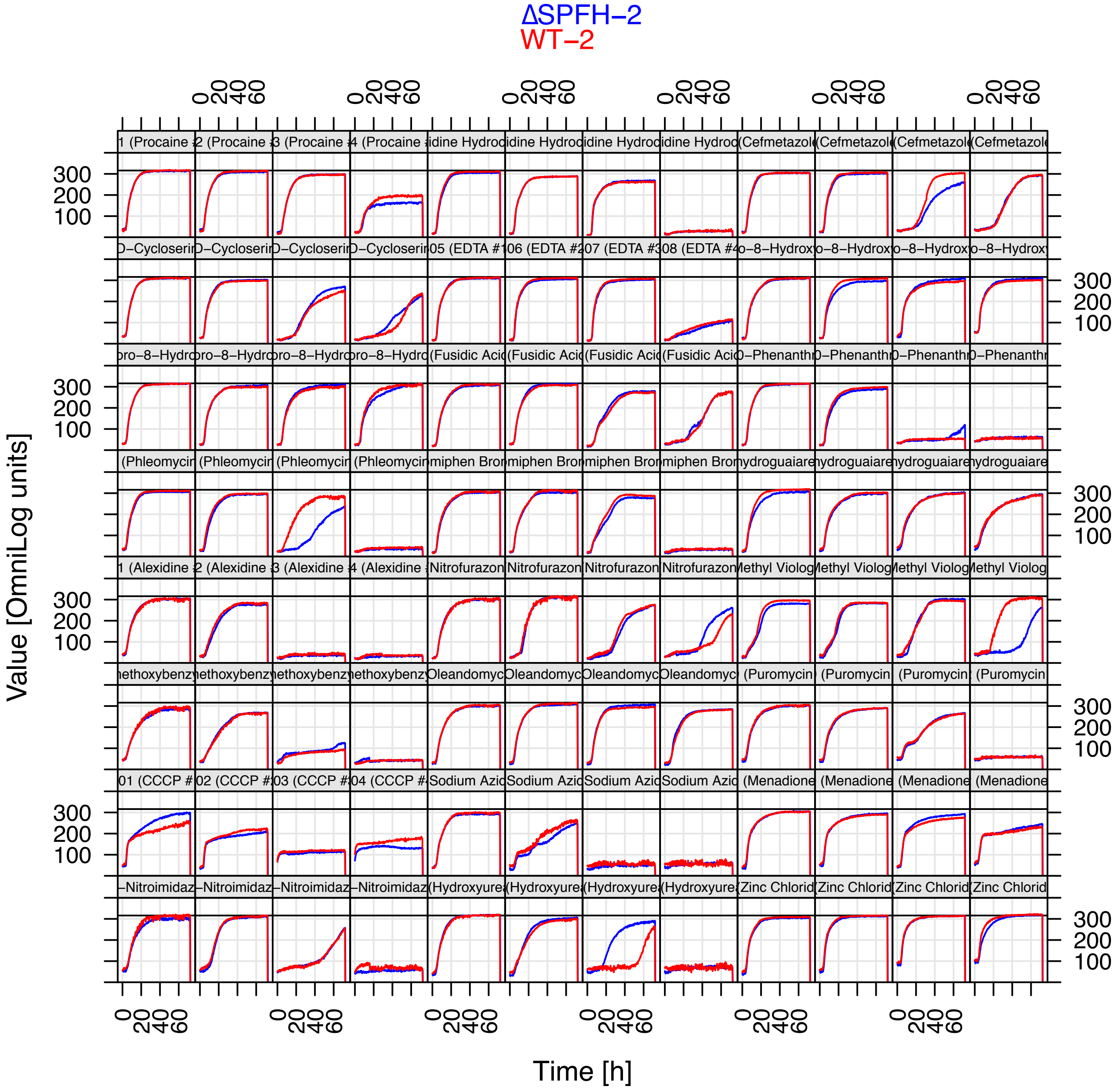

### PM15 (Chemicals)

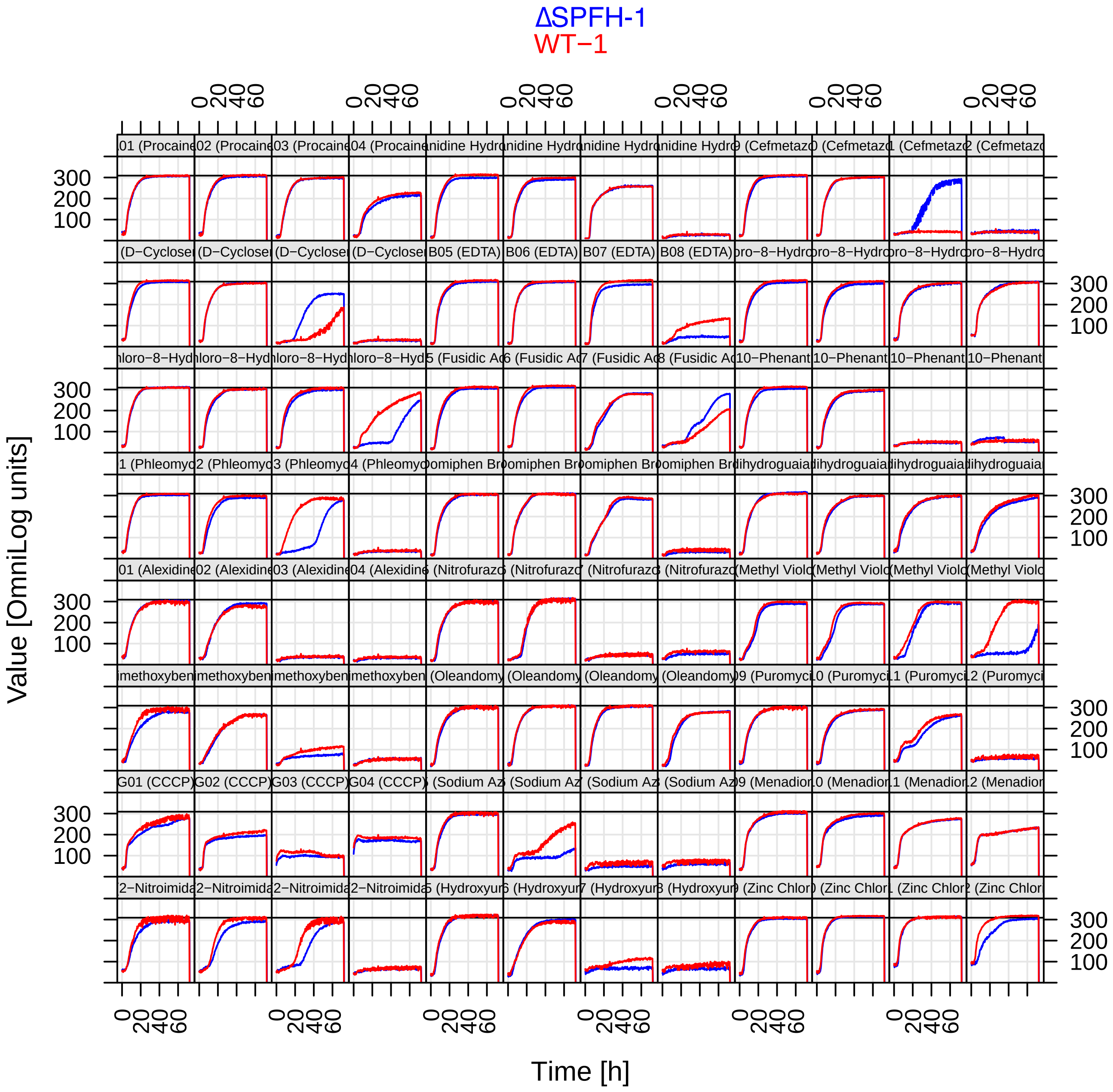

#### PM16 (Chemical Sensitivity Bacteria)

ΔSPFH−2
WT−2

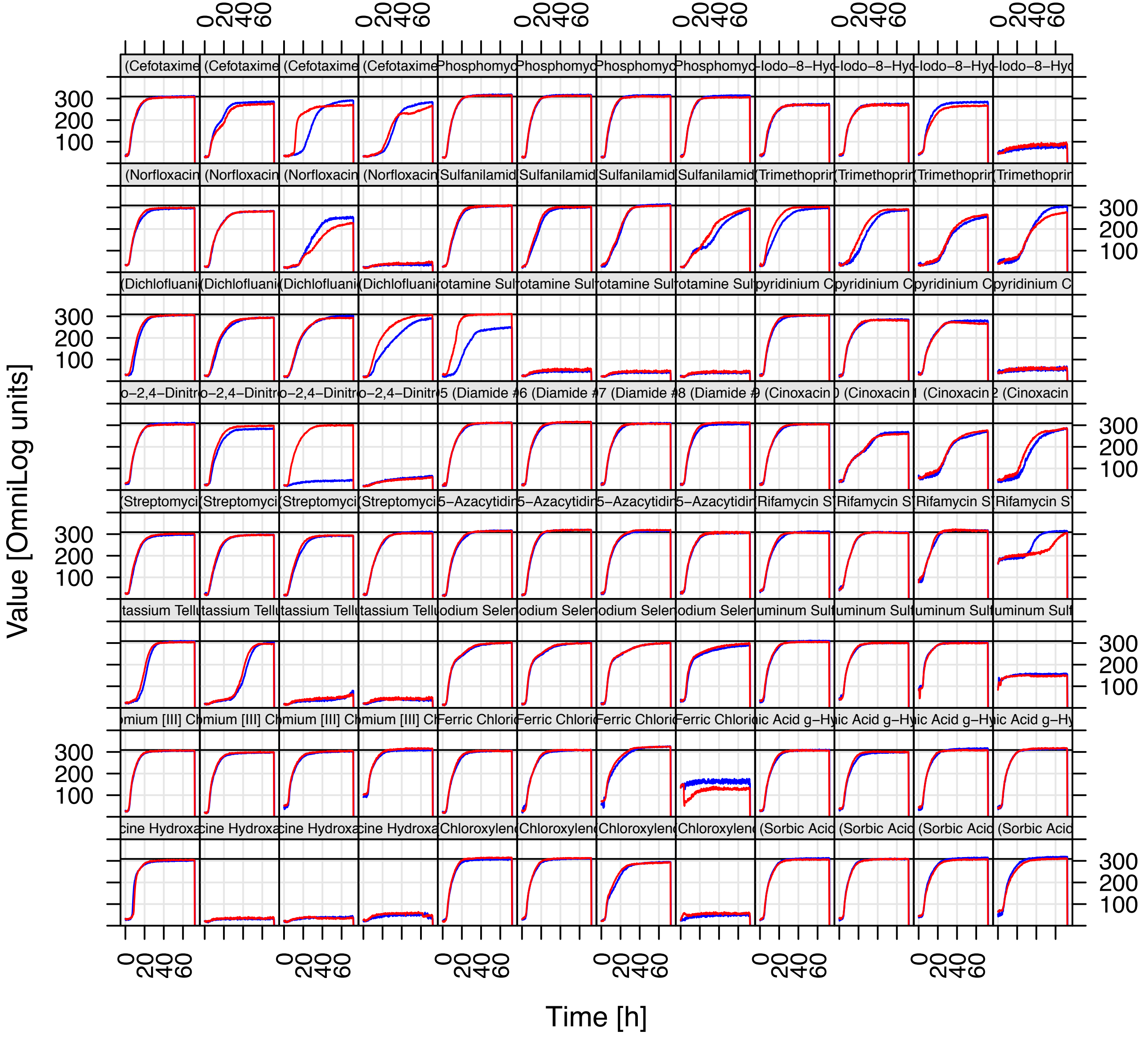

### PM16 (Chemicals)

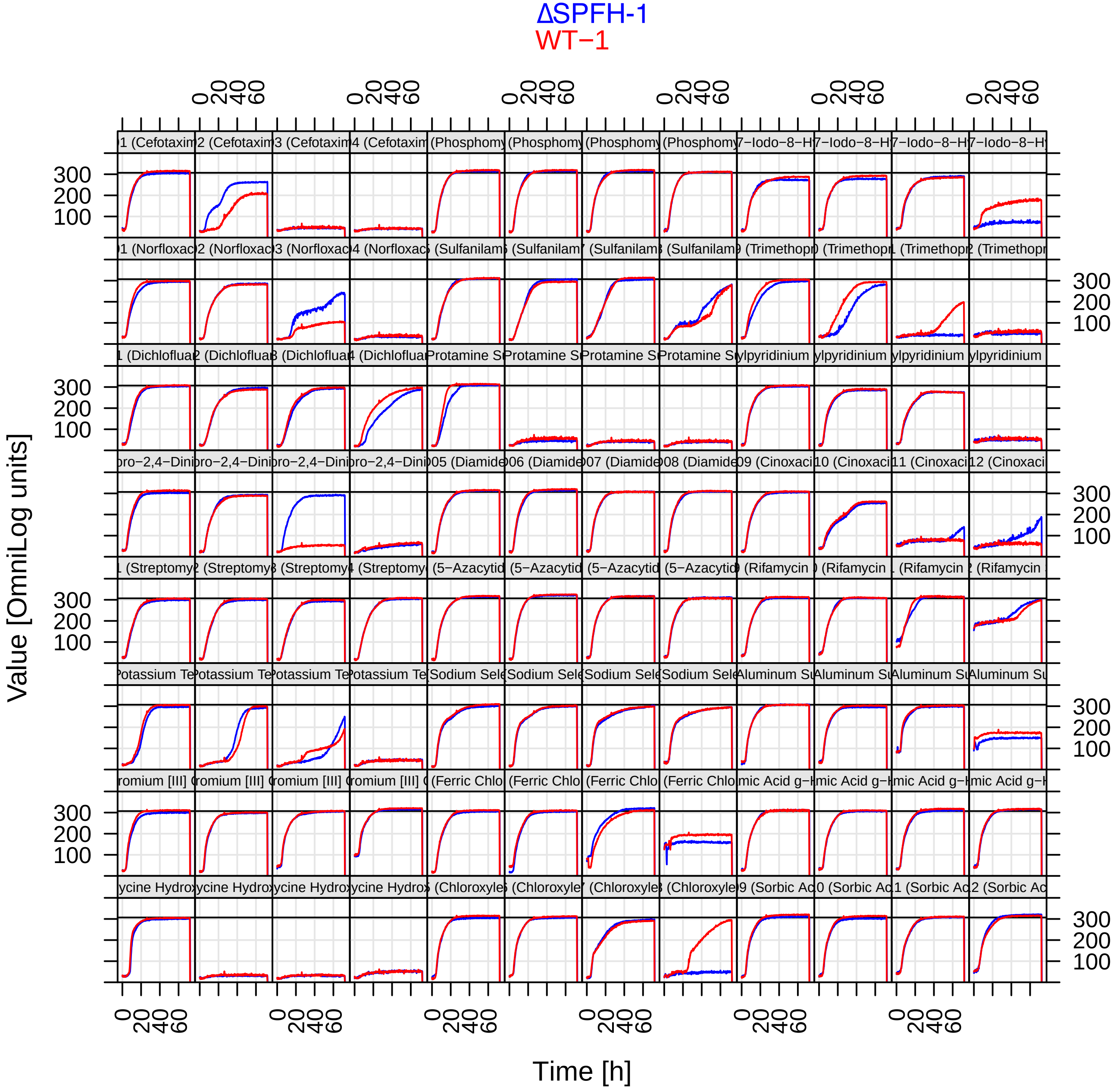

#### PM17A MicroPlate™

| | | | | | | | | | | | |
|---|---|---|---|---|---|---|---|---|---|---|---|
| A1<br>D-Serine<br>1 | A2<br>D-Serine<br>2 | A3<br>D-Serine<br>3 | A4<br>D-Serine<br>4 | A5<br>β-Chloro-L-alanine hydrochloride<br>1 | A6<br>β-Chloro-L-alanine hydrochloride<br>2 | A7<br>β-Chloro-L-alanine hydrochloride<br>3 | A8<br>β-Chloro-L-alanine hydrochloride<br>4 | A9<br>Thiosalicylic acid<br>1 | A10<br>Thiosalicylic acid<br>2 | A11<br>Thiosalicylic acid<br>3 | A12<br>Thiosalicylic acid<br>4 |
| B1<br>Sodium salicylate<br>1 | B2<br>Sodium salicylate<br>2 | B3<br>Sodium salicylate<br>3 | B4<br>Sodium salicylate<br>4 | B5<br>Hygromycin B<br>1 | B6<br>Hygromycin B<br>2 | B7<br>Hygromycin B<br>3 | B8<br>Hygromycin B<br>4 | B9<br>Ethionamide<br>1 | B10<br>Ethionamide<br>2 | B11<br>Ethionamide<br>3 | B12<br>Ethionamide<br>4 |
| C1<br>4-Aminopyridine<br>1 | C2<br>4-Aminopyridine<br>2 | C3<br>4-Aminopyridine<br>3 | C4<br>4-Aminopyridine<br>4 | C5<br>Sulfachloro-pyridazine<br>1 | C6<br>Sulfachloro-pyridazine<br>2 | C7<br>Sulfachloro-pyridazine<br>3 | C8<br>Sulfachloro-pyridazine<br>4 | C9<br>Sulfamono-methoxine<br>1 | C10<br>Sulfamono-methoxine<br>2 | C11<br>Sulfamono-methoxine<br>3 | C12<br>Sulfamono-methoxine<br>4 |
| D1<br>Oxycarboxin<br>1 | D2<br>Oxycarboxin<br>2 | D3<br>Oxycarboxin<br>3 | D4<br>Oxycarboxin<br>4 | D5<br>3-Amino-1,2,4-triazole<br>1 | D6<br>3-Amino-1,2,4-triazole<br>2 | D7<br>3-Amino-1,2,4-triazole<br>3 | D8<br>3-Amino-1,2,4-triazole<br>4 | D9<br>Chlorpromazine<br>1 | D10<br>Chlorpromazine<br>2 | D11<br>Chlorpromazine<br>3 | D12<br>Chlorpromazine<br>4 |
| E1<br>Niaproof<br>1 | E2<br>Niaproof<br>2 | E3<br>Niaproof<br>3 | E4<br>Niaproof<br>4 | E5<br>Compound 48/80<br>1 | E6<br>Compound 48/80<br>2 | E7<br>Compound 48/80<br>3 | E8<br>Compound 48/80<br>4 | E9<br>Sodium tungstate<br>1 | E10<br>Sodium tungstate<br>2 | E11<br>Sodium tungstate<br>3 | E12<br>Sodium tungstate<br>4 |
| F1<br>Lithium chloride<br>1 | F2<br>Lithium chloride<br>2 | F3<br>Lithium chloride<br>3 | F4<br>Lithium chloride<br>4 | F5<br>DL-Methionine hydroxamate<br>1 | F6<br>DL-Methionine hydroxamate<br>2 | F7<br>DL-Methionine hydroxamate<br>3 | F8<br>DL-Methionine hydroxamate<br>4 | F9<br>Tannic acid<br>1 | F10<br>Tannic acid<br>2 | F11<br>Tannic acid<br>3 | F12<br>Tannic acid<br>4 |
| G1<br>Chlorambucil<br>1 | G2<br>Chlorambucil<br>2 | G3<br>Chlorambucil<br>3 | G4<br>Chlorambucil<br>4 | G5<br>Cefamandole nafate<br>1 | G6<br>Cefamandole nafate<br>2 | G7<br>Cefamandole nafate<br>3 | G8<br>Cefamandole nafate<br>4 | G9<br>Cefoperazone<br>1 | G10<br>Cefoperazone<br>2 | G11<br>Cefoperazone<br>3 | G12<br>Cefoperazone<br>4 |
| H1<br>Cefsulodin<br>1 | H2<br>Cefsulodin<br>2 | H3<br>Cefsulodin<br>3 | H4<br>Cefsulodin<br>4 | H5<br>Caffeine<br>1 | H6<br>Caffeine<br>2 | H7<br>Caffeine<br>3 | H8<br>Caffeine<br>4 | H9<br>Phenylarsine oxide<br>1 | H10<br>Phenylarsine oxide<br>2 | H11<br>Phenylarsine oxide<br>3 | H12<br>Phenylarsine oxide<br>4 |

#### PM18C MicroPlate™

| | | | | | | | | | | | |
|---|---|---|---|---|---|---|---|---|---|---|---|
| A1<br>Ketoprofen<br>1 | A2<br>Ketoprofen<br>2 | A3<br>Ketoprofen<br>3 | A4<br>Ketoprofen<br>4 | A5<br>Sodium pyrophosphate decahydrate<br>1 | A6<br>Sodium pyrophosphate decahydrate<br>2 | A7<br>Sodium pyrophosphate decahydrate<br>3 | A8<br>Sodium pyrophosphate decahydrate<br>4 | A9<br>Thiamphenicol<br>1 | A10<br>Thiamphenicol<br>2 | A11<br>Thiamphenicol<br>3 | A12<br>Thiamphenicol<br>4 |
| B1<br>Trifluorothymidine<br>1 | B2<br>Trifluorothymidine<br>2 | B3<br>Trifluorothymidine<br>3 | B4<br>Trifluorothymidine<br>4 | B5<br>Pipemidic Acid<br>1 | B6<br>Pipemidic Acid<br>2 | B7<br>Pipemidic Acid<br>3 | B8<br>Pipemidic Acid<br>4 | B9<br>Azathioprine<br>1 | B10<br>Azathioprine<br>2 | B11<br>Azathioprine<br>3 | B12<br>Azathioprine<br>4 |
| C1<br>Poly-L-lysine<br>1 | C2<br>Poly-L-lysine<br>2 | C3<br>Poly-L-lysine<br>3 | C4<br>Poly-L-lysine<br>4 | C5<br>Sulfisoxazole<br>1 | C6<br>Sulfisoxazole<br>2 | C7<br>Sulfisoxazole<br>3 | C8<br>Sulfisoxazole<br>4 | C9<br>Pentachloro-phenol<br>1 | C10<br>Pentachloro-phenol<br>2 | C11<br>Pentachloro-phenol<br>3 | C12<br>Pentachloro-phenol<br>4 |
| D1<br>Sodium m-arsenite<br>1 | D2<br>Sodium m-arsenite<br>2 | D3<br>Sodium m-arsenite<br>3 | D4<br>Sodium m-arsenite<br>4 | D5<br>Sodium bromate<br>1 | D6<br>Sodium bromate<br>2 | D7<br>Sodium bromate<br>3 | D8<br>Sodium bromate<br>4 | D9<br>Lidocane<br>1 | D10<br>Lidocane<br>2 | D11<br>Lidocane<br>3 | D12<br>Lidocane<br>4 |
| E1<br>Sodium metasilicate<br>1 | E2<br>Sodium metasilicate<br>2 | E3<br>Sodium metasilicate<br>3 | E4<br>Sodium metasilicate<br>4 | E5<br>Sodium m-periodate<br>1 | E6<br>Sodium m-periodate<br>2 | E7<br>Sodium m-periodate<br>3 | E8<br>Sodium m-periodate<br>4 | E9<br>Antimony (III) chloride<br>1 | E10<br>Antimony (III) chloride<br>2 | E11<br>Antimony (III) chloride<br>3 | E12<br>Antimony (III) chloride<br>4 |
| F1<br>Semicarbazide<br>1 | F2<br>Semicarbazide<br>2 | F3<br>Semicarbazide<br>3 | F4<br>Semicarbazide<br>4 | F5<br>Tinidazole<br>1 | F6<br>Tinidazole<br>2 | F7<br>Tinidazole<br>3 | F8<br>Tinidazole<br>4 | F9<br>Aztreonam<br>1 | F10<br>Aztreonam<br>2 | F11<br>Aztreonam<br>3 | F12<br>Aztreonam<br>4 |
| G1<br>Triclosan<br>1 | G2<br>Triclosan<br>2 | G3<br>Triclosan<br>3 | G4<br>Triclosan<br>4 | G5<br>3,5-Diamino-1,2,4-triazole (Guanazole)<br>1 | G6<br>3,5-Diamino-1,2,4-triazole (Guanazole)<br>2 | G7<br>3,5-Diamino-1,2,4-triazole (Guanazole)<br>3 | G8<br>3,5-Diamino-1,2,4-triazole (Guanazole)<br>4 | G9<br>Myricetin<br>1 | G10<br>Myricetin<br>2 | G11<br>Myricetin<br>3 | G12<br>Myricetin<br>4 |
| H1<br>5-fluoro-5'-deoxyuridine<br>1 | H2<br>5-fluoro-5'-deoxyuridine<br>2 | H3<br>5-fluoro-5'-deoxyuridine<br>3 | H4<br>5-fluoro-5'-deoxyuridine<br>4 | H5<br>2-Phenylphenol<br>1 | H6<br>2-Phenylphenol<br>2 | H7<br>2-Phenylphenol<br>3 | H8<br>2-Phenylphenol<br>4 | H9<br>Plumbagin<br>1 | H10<br>Plumbagin<br>2 | H11<br>Plumbagin<br>3 | H12<br>Plumbagin<br>4 |

#### PM17 (Chemical Sensitivity Bacteria)

ΔSPFH−2
WT−2

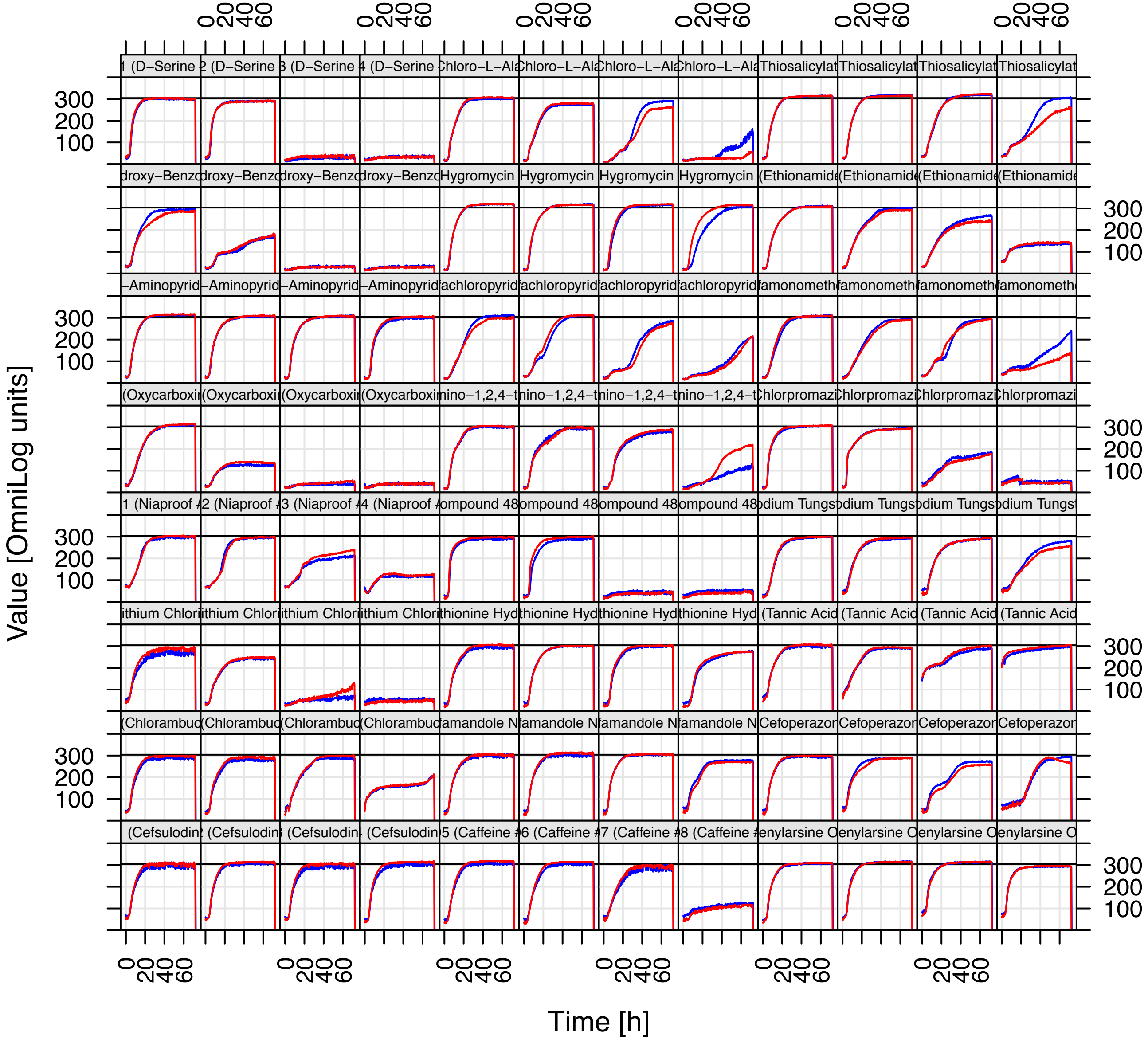

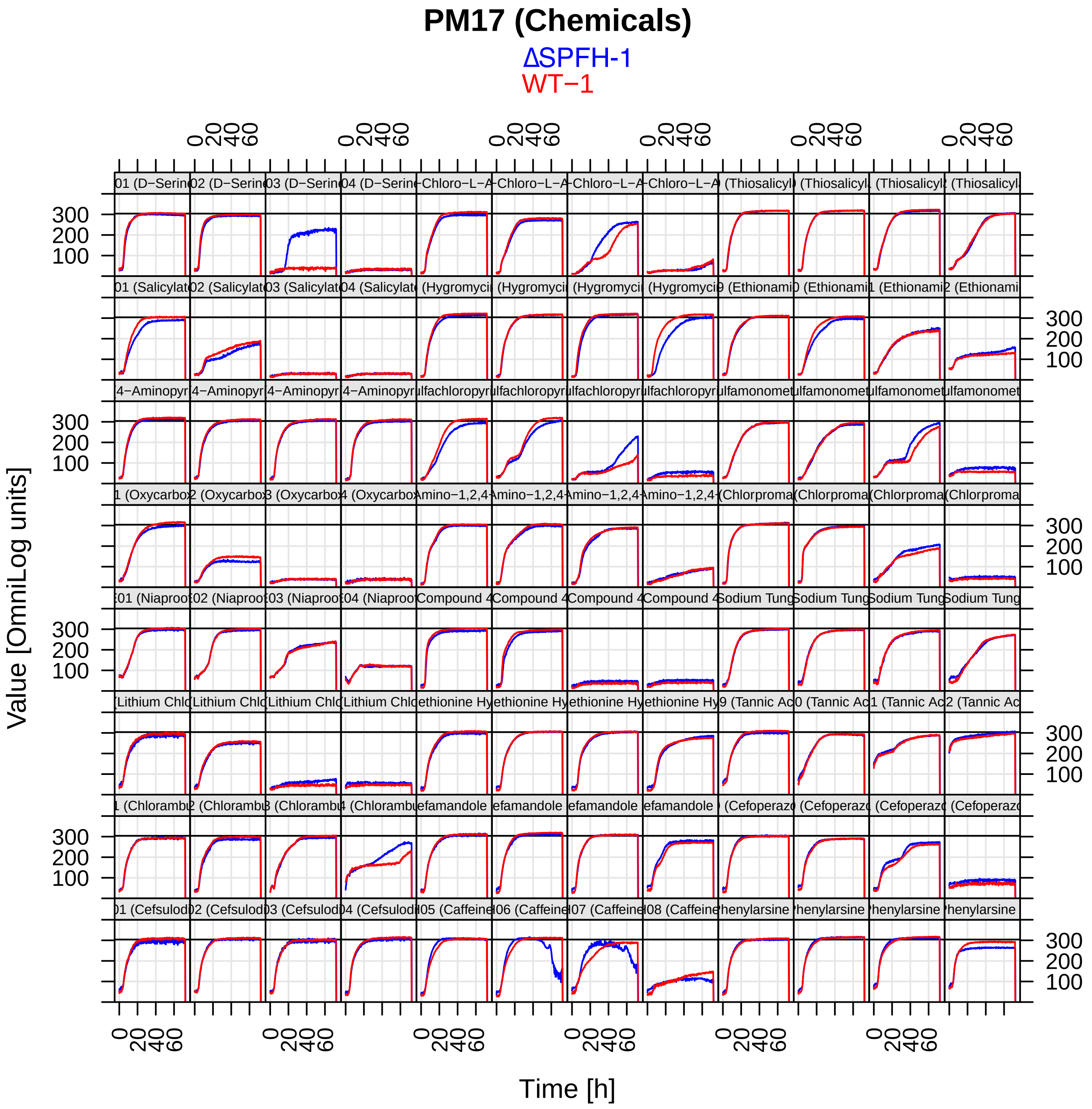
PM17 (Chemicals)
ΔSPFH-1
WT−1
Value [OmniLog units]
Time [h]

### PM18 (Chemical Sensitivity Bacteria)

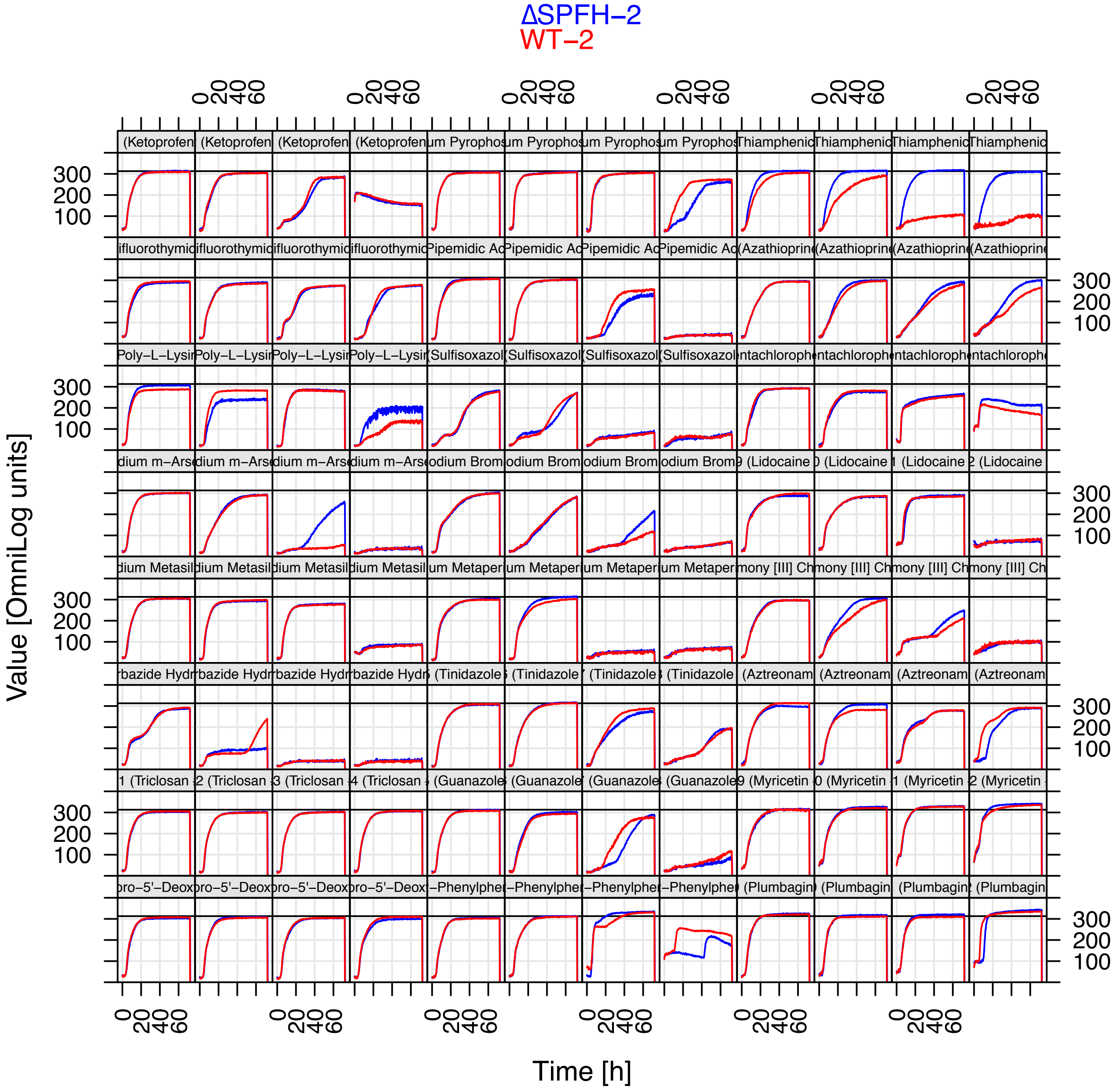

#### PM18 (Chemicals)

ΔSPFH-1
WT−1

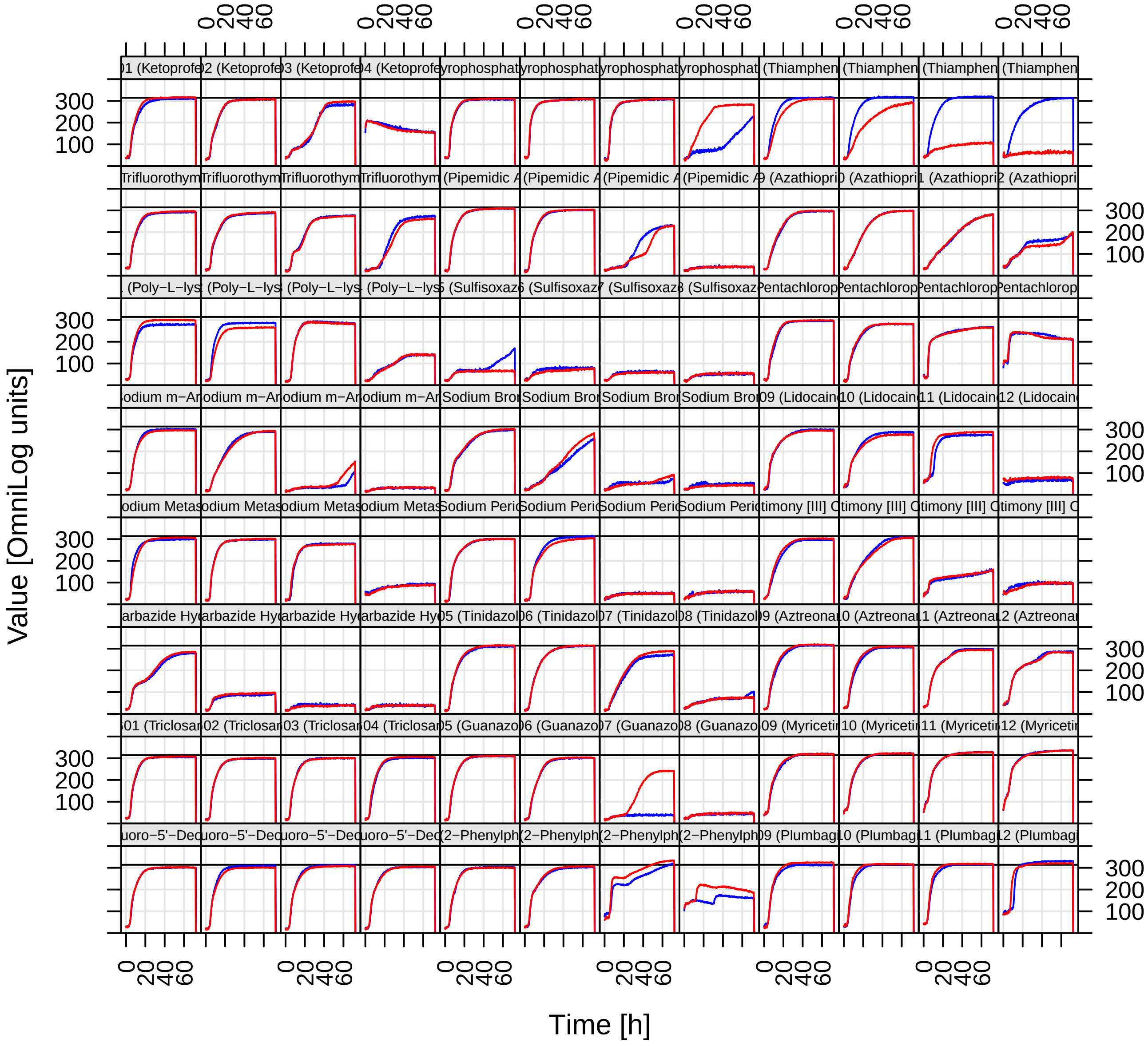

#### PM19 MicroPlate™

| | | | | | | | | | | | |
|---|---|---|---|---|---|---|---|---|---|---|---|
| A1<br>Josamycin<br>1 | A2<br>Josamycin<br>2 | A3<br>Josamycin<br>3 | A4<br>Josamycin<br>4 | A5<br>Gallic acid<br>1 | A6<br>Gallic acid<br>2 | A7<br>Gallic acid<br>3 | A8<br>Gallic acid<br>4 | A9<br>Coumarin<br>1 | A10<br>Coumarin<br>2 | A11<br>Coumarin<br>3 | A12<br>Coumarin<br>4 |
| B1<br>Methyltrioctyl-ammonium chloride<br>1 | B2<br>Methyltrioctyl-ammonium chloride<br>2 | B3<br>Methyltrioctyl-ammonium chloride<br>3 | B4<br>Methyltrioctyl-ammonium chloride<br>4 | B5<br>Harmane<br>1 | B6<br>Harmane<br>2 | B7<br>Harmane<br>3 | B8<br>Harmane<br>4 | B9<br>2,4-Dintrophenol<br>1 | B10<br>2,4-Dintrophenol<br>2 | B11<br>2,4-Dintrophenol<br>3 | B12<br>2,4-Dintrophenol<br>4 |
| C1<br>Chlorhexidine<br>1 | C2<br>Chlorhexidine<br>2 | C3<br>Chlorhexidine<br>3 | C4<br>Chlorhexidine<br>4 | C5<br>Umbelliferone<br>1 | C6<br>Umbelliferone<br>2 | C7<br>Umbelliferone<br>3 | C8<br>Umbelliferone<br>4 | C9<br>Cinnamic acid<br>1 | C10<br>Cinnamic acid<br>2 | C11<br>Cinnamic acid<br>3 | C12<br>Cinnamic acid<br>4 |
| D1<br>Disulphiram<br>1 | D2<br>Disulphiram<br>2 | D3<br>Disulphiram<br>3 | D4<br>Disulphiram<br>4 | D5<br>Iodonitro Tetrazolium Violet<br>1 | D6<br>Iodonitro Tetrazolium Violet<br>2 | D7<br>Iodonitro Tetrazolium Violet<br>3 | D8<br>Iodonitro Tetrazolium Violet<br>4 | D9<br>Phenyl- methyl-sulfonyl-fluoride (PMSF)<br>1 | D10<br>Phenyl- methyl-sulfonyl-fluoride (PMSF)<br>2 | D11<br>Phenyl- methyl-sulfonyl-fluoride (PMSF)<br>3 | D12<br>Phenyl- methyl-sulfonyl-fluoride (PMSF)<br>4 |
| E1<br>FCCP<br>1 | E2<br>FCCP<br>2 | E3<br>FCCP<br>3 | E4<br>FCCP<br>4 | E5<br>D,L-Thioctic Acid<br>1 | E6<br>D,L-Thioctic Acid<br>2 | E7<br>D,L-Thioctic Acid<br>3 | E8<br>D,L-Thioctic Acid<br>4 | E9<br>Lawsone<br>1 | E10<br>Lawsone<br>2 | E11<br>Lawsone<br>3 | E12<br>Lawsone<br>4 |
| F1<br>Phenethicillin<br>1 | F2<br>Phenethicillin<br>2 | F3<br>Phenethicillin<br>3 | F4<br>Phenethicillin<br>4 | F5<br>Blasticidin S<br>1 | F6<br>Blasticidin S<br>2 | F7<br>Blasticidin S<br>3 | F8<br>Blasticidin S<br>4 | F9<br>Sodium caprylate<br>1 | F10<br>Sodium caprylate<br>2 | F11<br>Sodium caprylate<br>3 | F12<br>Sodium caprylate<br>4 |
| G1<br>Lauryl sulfobetaine<br>1 | G2<br>Lauryl sulfobetaine<br>2 | G3<br>Lauryl sulfobetaine<br>3 | G4<br>Lauryl sulfobetaine<br>4 | G5<br>Dihydro-streptomycin<br>1 | G6<br>Dihydro-streptomycin<br>2 | G7<br>Dihydro-streptomycin<br>3 | G8<br>Dihydro-streptomycin<br>4 | G9<br>Hydroxylamine<br>1 | G10<br>Hydroxylamine<br>2 | G11<br>Hydroxylamine<br>3 | G12<br>Hydroxylamine<br>4 |
| H1<br>Hexammine cobalt (III) chloride<br>1 | H2<br>Hexammine cobalt (III) chloride<br>2 | H3<br>Hexammine cobalt (III) chloride<br>3 | H4<br>Hexammine cobalt (III) chloride<br>4 | H5<br>Thioglycerol<br>1 | H6<br>Thioglycerol<br>2 | H7<br>Thioglycerol<br>3 | H8<br>Thioglycerol<br>4 | H9<br>Polymyxin B<br>1 | H10<br>Polymyxin B<br>2 | H11<br>Polymyxin B<br>3 | H12<br>Polymyxin B<br>4 |

#### PM20B MicroPlate™

| | | | | | | | | | | | |
|---|---|---|---|---|---|---|---|---|---|---|---|
| A1<br>Amitriptyline<br>1 | A2<br>Amitriptyline<br>2 | A3<br>Amitriptyline<br>3 | A4<br>Amitriptyline<br>4 | A5<br>Apramycin<br>1 | A6<br>Apramycin<br>2 | A7<br>Apramycin<br>3 | A8<br>Apramycin<br>4 | A9<br>Benserazide<br>1 | A10<br>Benserazide<br>2 | A11<br>Benserazide<br>3 | A12<br>Benserazide<br>4 |
| B1<br>Orphenadrine<br>1 | B2<br>Orphenadrine<br>2 | B3<br>Orphenadrine<br>3 | B4<br>Orphenadrine<br>4 | B5<br>D,L-Propranolol<br>1 | B6<br>D,L-Propranolol<br>2 | B7<br>D,L-Propranolol<br>3 | B8<br>D,L-Propranolol<br>4 | B9<br>Tetrazolium violet<br>1 | B10<br>Tetrazolium violet<br>2 | B11<br>Tetrazolium violet<br>3 | B12<br>Tetrazolium violet<br>4 |
| C1<br>Thioridazine<br>1 | C2<br>Thioridazine<br>2 | C3<br>Thioridazine<br>3 | C4<br>Thioridazine<br>4 | C5<br>Atropine<br>1 | C6<br>Atropine<br>2 | C7<br>Atropine<br>3 | C8<br>Atropine<br>4 | C9<br>Ornidazole<br>1 | C10<br>Ornidazole<br>2 | C11<br>Ornidazole<br>3 | C12<br>Ornidazole<br>4 |
| D1<br>Proflavine<br>1 | D2<br>Proflavine<br>2 | D3<br>Proflavine<br>3 | D4<br>Proflavine<br>4 | D5<br>Ciprofloxacin<br>1 | D6<br>Ciprofloxacin<br>2 | D7<br>Ciprofloxacin<br>3 | D8<br>Ciprofloxacin<br>4 | D9<br>18-Crown-6 ether<br>1 | D10<br>18-Crown-6 ether<br>2 | D11<br>18-Crown-6 ether<br>3 | D12<br>18-Crown-6 ether<br>4 |
| E1<br>Crystal violet<br>1 | E2<br>Crystal violet<br>2 | E3<br>Crystal violet<br>3 | E4<br>Crystal violet<br>4 | E5<br>Dodine<br>1 | E6<br>Dodine<br>2 | E7<br>Dodine<br>3 | E8<br>Dodine<br>4 | E9<br>Hexa-chlorophene<br>1 | E10<br>Hexa-chlorophene<br>2 | E11<br>Hexa-chlorophene<br>3 | E12<br>Hexa-chlorophene<br>4 |
| F1<br>4-Hydroxy-coumarin<br>1 | F2<br>4-Hydroxy-coumarin<br>2 | F3<br>4-Hydroxy-coumarin<br>3 | F4<br>4-Hydroxy-coumarin<br>4 | F5<br>Oxytetracycline<br>1 | F6<br>Oxytetracycline<br>2 | F7<br>Oxytetracycline<br>3 | F8<br>Oxytetracycline<br>4 | F9<br>Pridinol<br>1 | F10<br>Pridinol<br>2 | F11<br>Pridinol<br>3 | F12<br>Pridinol<br>4 |
| G1<br>Captan<br>1 | G2<br>Captan<br>2 | G3<br>Captan<br>3 | G4<br>Captan<br>4 | G5<br>3,5-Dinitro-benzene<br>1 | G6<br>3,5-Dinitro-benzene<br>2 | G7<br>3,5-Dinitro-benzene<br>3 | G8<br>3,5-Dinitro-benzene<br>4 | G9<br>8-Hydroxy-quinoline<br>1 | G10<br>8-Hydroxy-quinoline<br>2 | G11<br>8-Hydroxy-quinoline<br>3 | G12<br>8-Hydroxy-quinoline<br>4 |
| H1<br>Patulin<br>1 | H2<br>Patulin<br>2 | H3<br>Patulin<br>3 | H4<br>Patulin<br>4 | H5<br>Tolylfluanid<br>1 | H6<br>Tolylfluanid<br>2 | H7<br>Tolylfluanid<br>3 | H8<br>Tolylfluanid<br>4 | H9<br>Troleandomycin<br>1 | H10<br>Troleandomycin<br>2 | H11<br>Troleandomycin<br>3 | H12<br>Troleandomycin<br>4 |

### PM19 (Chemical Sensitivity Bacteria)

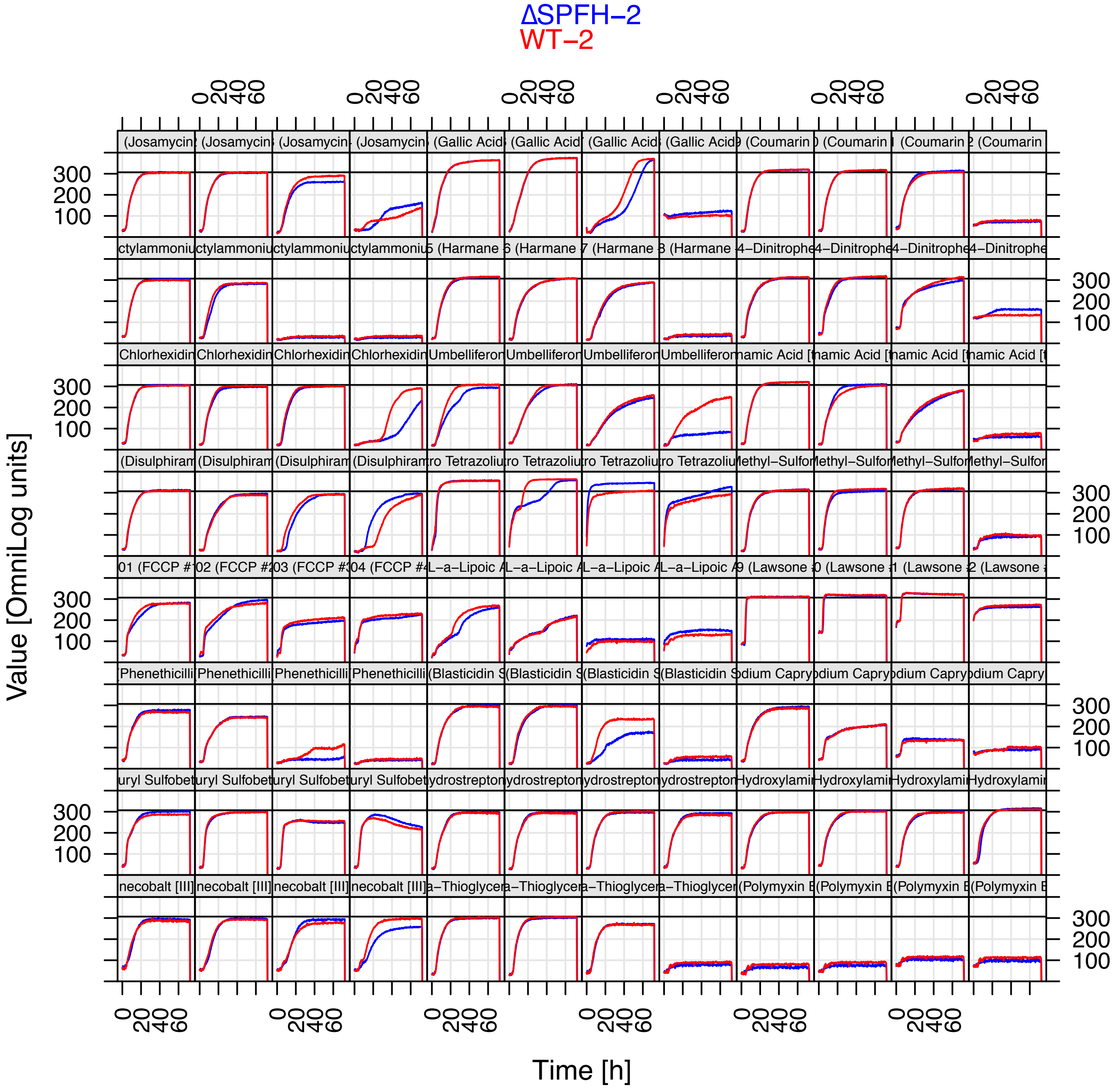

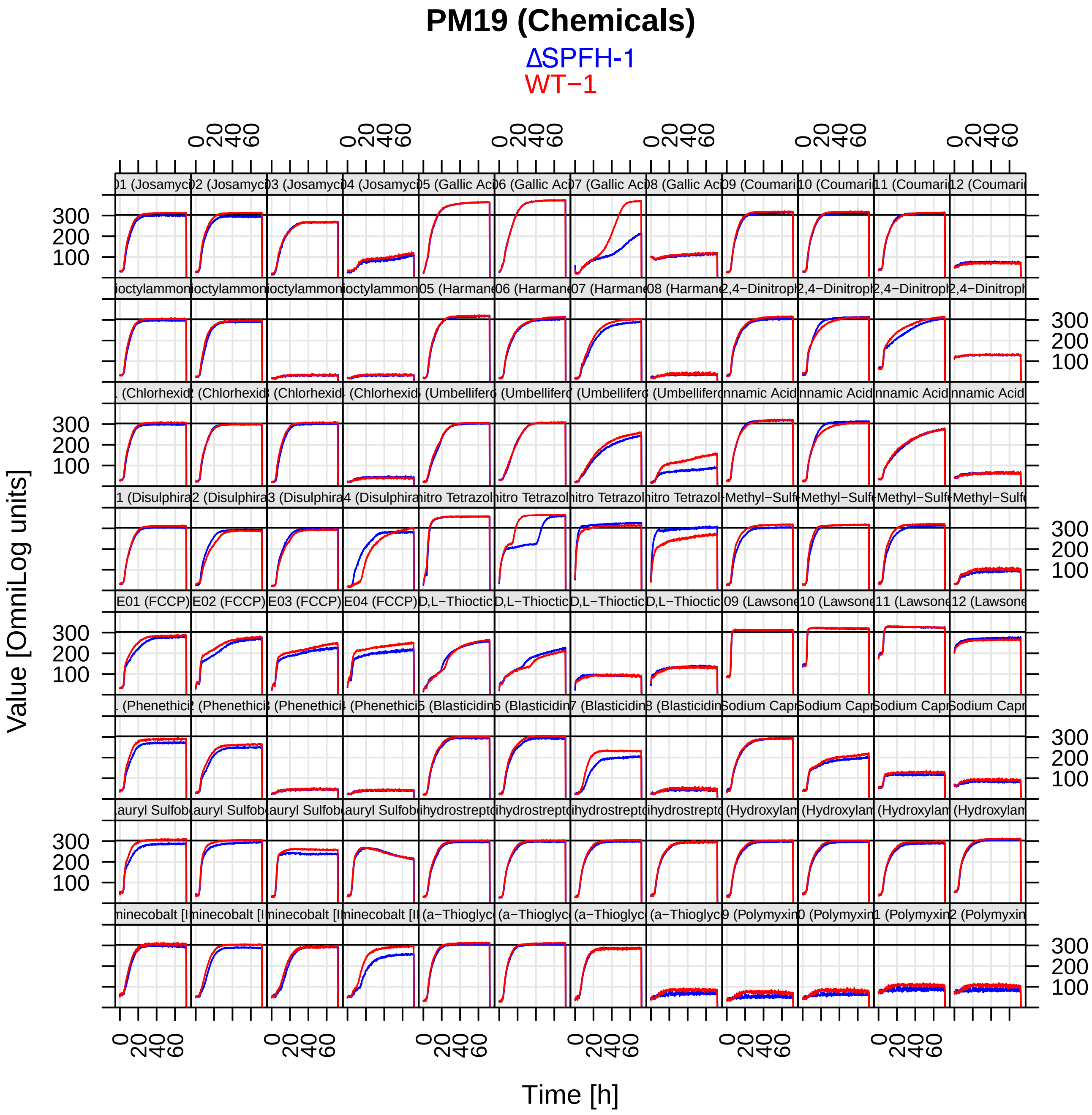

PM19 (Chemicals)
ΔSPFH-1
WT−1
Value [OmniLog units]
Time [h]

### PM20 (Chemical Sensitivity Bacteria)

ΔSPFH−2
WT−2

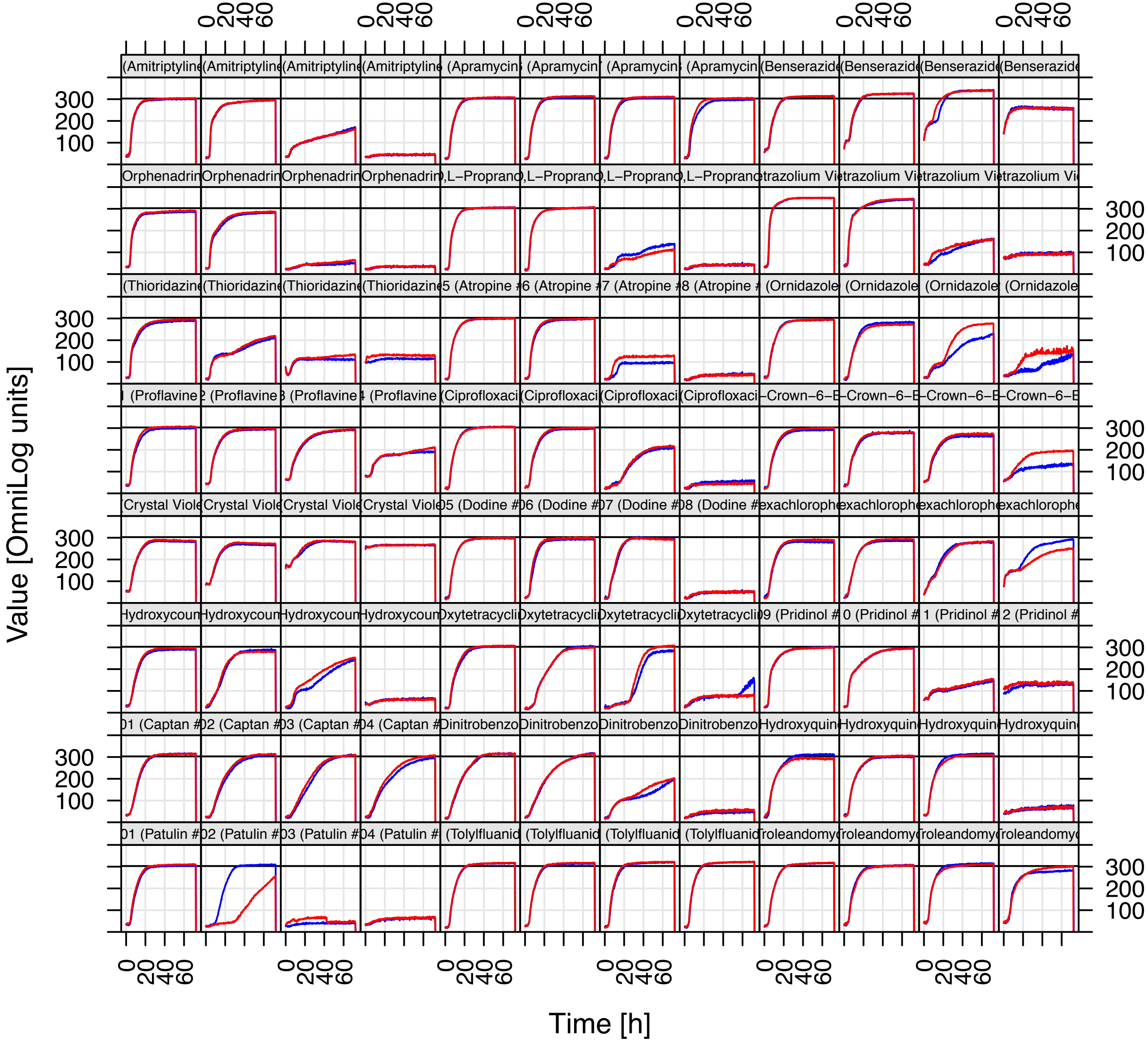

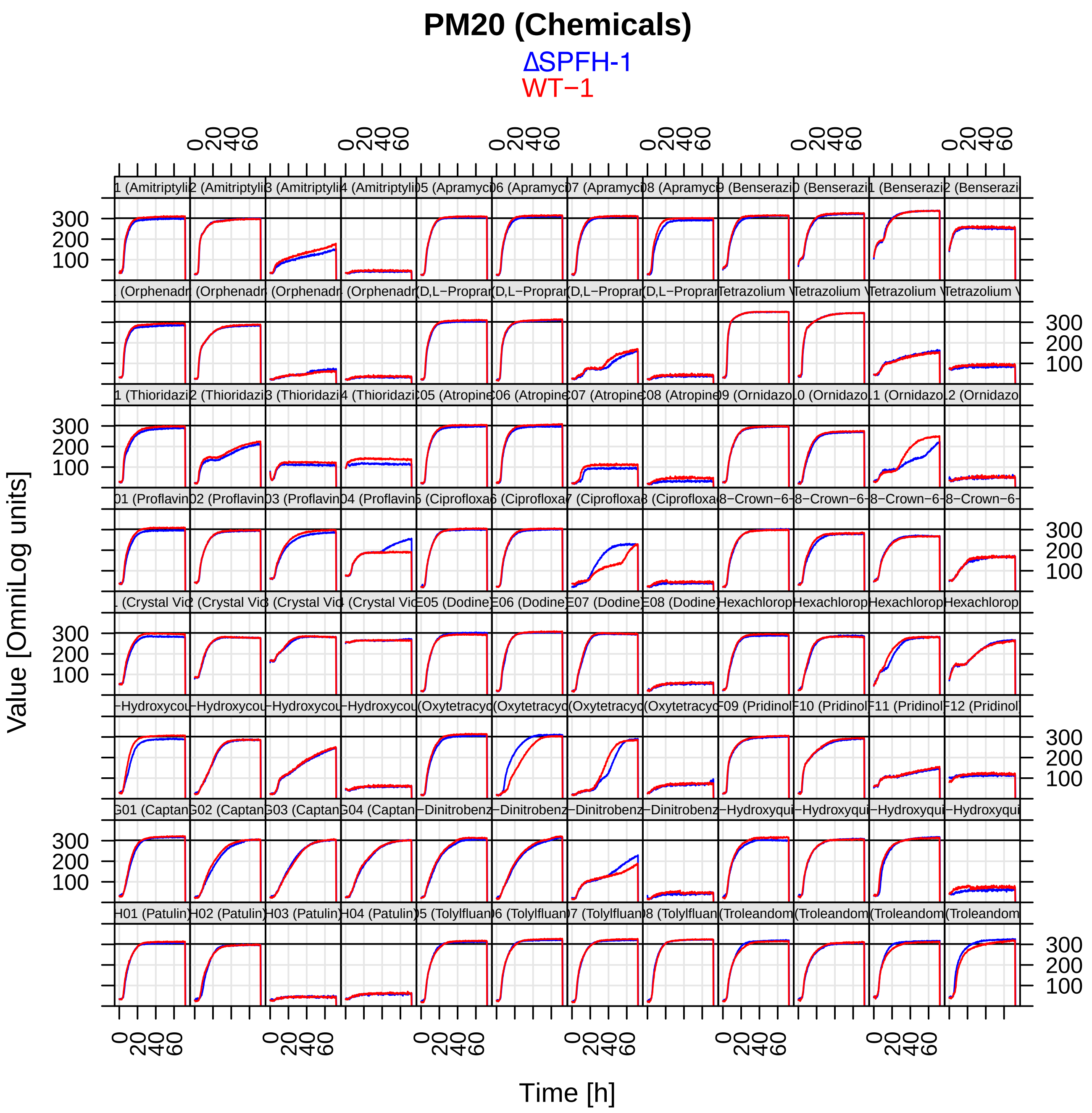
PM20 (Chemicals)
ΔSPFH-1
WT−1
Value [OmniLog units]
Time [h]
